# Stepwise emergence of recombination suppression precedes fissiparous asexuality in the *planarian Schmidtea mediterranea*

**DOI:** 10.1101/2025.09.12.675808

**Authors:** Jeremias N. Brand, Ajinkya Bharatraj Patil, Luca Pandolfini, Kira Zadesenets, Nikolay Rubtsov, Laura Robledillo, Meng Zhang, André Marques, Jochen C. Rink

## Abstract

A central paradox in evolutionary biology is the rarity of asexual reproduction, which is often attributed to developmental constraints and long-term costs. Yet, fissiparous asexuality—where animals split and regenerate—is widespread among planarians, hinting at genomic features predisposing them to asexuality. We investigated the genomic underpinnings and evolutionary consequences of asexuality in the planarian *Schmidtea mediterranea*, which exists as both obligately fissiparous and sexual strains. We generated a haplotype-phased genome assembly of the asexual strain and collected population genomic data from laboratory and wild populations to uncover extensive heterozygous chromosomal rearrangements affecting all chromosomes. We show that these rearrangements arose in a sexually reproducing ancestor without directly disrupting reproductive genes but instead progressively suppressing recombination across the genome. The asexual genome exhibits minimal deleterious mutation accumulation, indicating a low cost of asexuality. Population-genomic data show this strain lacks any detectable sexual reproduction and originated recently (0.17–0.4 Ma), but the young age is insufficient to explain the low mutational burden. Instead, planarians may be able to exploit the lack of a single-cell bottleneck in fissiparous reproduction to mitigate the costs of asexuality. Altogether, our results support a model in which stepwise recombination suppression due to structural rearrangements eroded the benefits of sex and enabled the emergence of fissiparous asexuality in *S. mediterranea*.

## Introduction

Sexual reproduction is predicted to be more costly than asexual reproduction^1,2^ due to three main factors: the two-fold cost of males^2^, the cost of recombination breaking favorable combinations of alleles^3^, and the costs associated with mating^4^. Consequently, asexual mutants should spread throughout a population and drive sexual lineages to extinction^1^. The explanation for why sexual reproduction remains the rule rather than the exception despite these costs is thought to be the evolutionary advantages sex confers through efficient selection^1,5^, especially in heterogeneous environments^6,7^ or during coevolution with parasites^8,9^. Sexual recombination can reduce negative epistasis^10^, avoid Hill-Robertson interference^10^ by allowing beneficial mutations to spread independently of harmful ones^11,12^, and prevent the accumulation of deleterious mutations in small populations (Muller’s ratchet)^11^, by restoring the least-loaded genotype. Consequently, asexual lineages deprived of the advantages of sexual reproduction should accumulate deleterious mutations over time and ultimately go extinct—a process known as mutational meltdown^5,11^. However, while numerous studies find increased accumulation of deleterious mutations in parthenogenetic asexuals^13–19^, some ancient asexual lineages are associated with equal^20^ or even more efficient purifying selection compared to their sexual relatives^21,22^. This suggests that some asexual lineages have evolved mechanisms to mitigate mutational meltdown or generate new genetic diversity^23^. In addition, alternative asexual reproduction strategies, e.g., via budding or fission of the parental organism into two new entities, a form of asexuality occurring in the majority of animal phyla^24^, have been poorly studied so far. Budding and fission are particularly interesting because they lack the single-cell bottleneck between generations that is thought to be essential for ensuring the clonality of the soma^24,25^. The increased intra-individual genetic diversity in such systems^26^ raises the potential for evolutionary conflicts, but also provides a conceptual possibility for reducing the effective mutation rate^27^, which could prevent mutational meltdown^24^.

Planarian flatworms are an excellent model to understand the emergence and consequences of fissiparous asexuality, because the hundreds of planarian species worldwide display a fascinating spectrum of reproductive strategies, ranging from hermaphroditic sexual reproduction, parthenogenesis, to asexual fissiparous reproduction^28^. In addition, the molecularly tractable laboratory model species *Schmidtea mediterranea* is of particular interest. Usually studied for its whole-body regeneration ability and abundant adult pluripotent stem cells^29,30^, this species exists in both asexual and sexual strains. The asexual strains reproduce exclusively by fission and subsequent whole-body regeneration and never develop sexual organs. The sexual strains develop hermaphroditic reproductive organs and lay egg capsules containing multiple fertilized zygotes. The planarian research community primarily studies a clonal strain (CIW4) originating from a single animal collected from a fountain in Montjuïc Park, Barcelona, Spain, in 1998^33–35,31–33^. The CIW4 strain is exclusively fissiparous and never develops reproductive organs. The somewhat less-studied exclusively sexual laboratory strains were established from specimens collected in Sardinia and do not undergo fission^34,35^. *S. mediterranea* has a restricted distribution in the western Mediterranean, with asexual populations on the Catalan coast and the Balearic Islands and sexual populations in Corsica, Sardinia, Sicily, and the Tunisian coast^36–44^. No co-occurrence between the two strains has been reported^36–44^ (Fig. 1a). The current distribution has been hypothesized to reflect the fragmentation of an ancestral population due to microplate tectonics^38,39^, which accounts for the low dispersal ability of planarians due to their high desiccation susceptibility^28,45^. Consistently, molecular dating analyses suggest a tentative split between the sexual and asexual populations ∼8.36 million years ago (Ma), but such analyses remain challenging in planarians due to the lack of accurate calibration data^46^. Overall, *S. mediterranea* offers both the comparative analysis of molecularly tractable sexual and asexual laboratory strains, as well as a biogeographical distribution range that may inform on the origins of asexuality.

**Figure 1.**
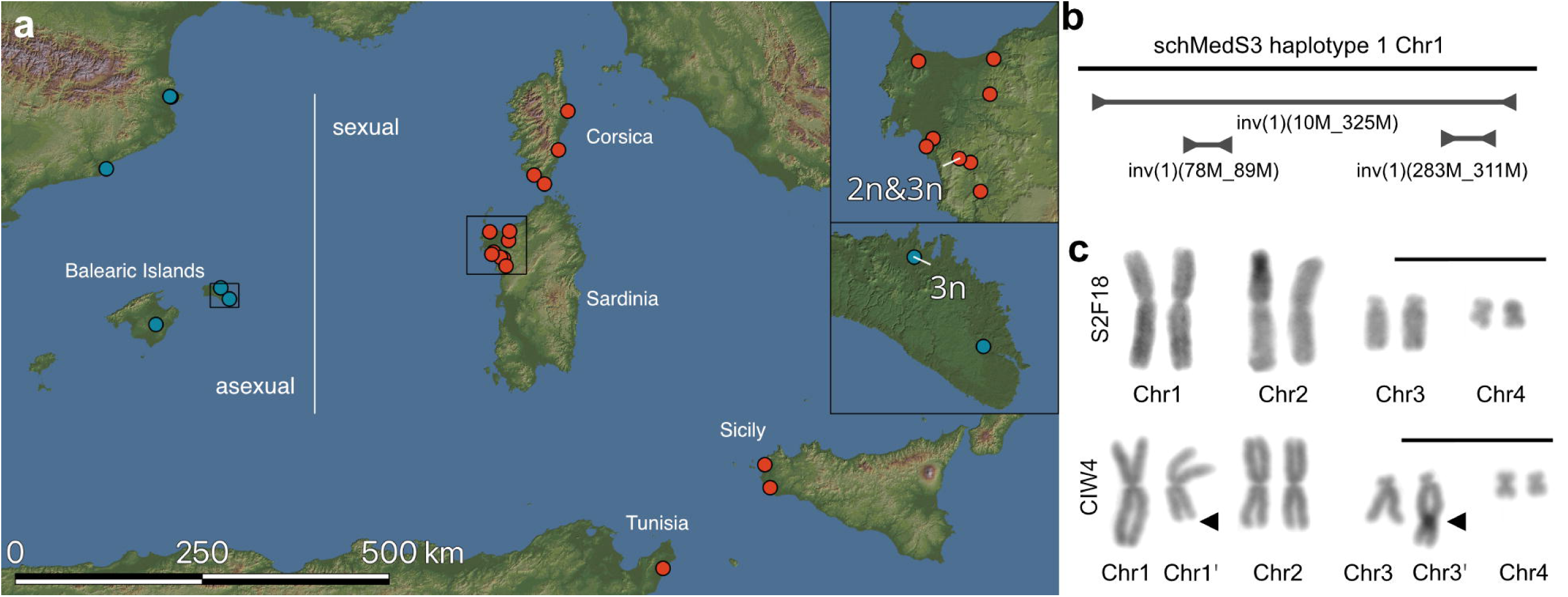
Geographic distribution and chromosomal characteristics of *Schmidtea mediterranea* sexual and asexual populations. **a** Map showing the localities where *S. mediterranea* has been found. Asexual populations (blue) have been found on the Spanish mainland and the Balearic Islands. Sexual populations (red) have been found on Corsica, Sardinia, Sicily, and Tunisia. Insets show details of collection sites on Sardinia (top) and Menorca (bottom). *S. mediterranea* is usually diploid (2n = 8) but, as indicated on the insets, there are one asexual and one sexual triploid population (3n = 12). **b** Schematic representation of Chromosome 1 (Chr1) in haplotype 1 of the schMedS3 genome assembly of the sexual laboratory strain (S2F18), indicating three chromosome inversions, one of which spans almost the entire length of Chr1^48^. Population heterozygosity data show that the large inversion is present in sexual populations from Corsica and Sardinia, but absent in Sicily and Tunisia^52^. **c** Chromosome spread of the sexual laboratory strain (S2F18) and the asexual laboratory strain (CIW4). Arrows indicate the reciprocal translocation between Chr1 and Chr3, characteristic of CIW4, resulting in a short, derived Chr1’ and a long, derived Chr3’. Scale bars represent 10 μm.

The genomic causes and consequences of asexuality in planarians remain understudied due to a lack of suitable genomic resources. Planarian genomes are difficult to assemble due to extreme compositional bias (70% A/T), the abundance of unusually large repeats, and inbreeding-resistant heterozygosity^43,47–51^. Chromosome-scale assemblies of sexually reproducing strains have only recently become available, collectively revealing rapid structural divergence in planarian genomes^5148^. The genome assembly of the sexual strain of *S. mediterranea* reveals three large inversions on chromosome 1^48,52^, which repress recombination on that chromosome^43^ (Fig. 1b), and have been interpreted as a first step in the evolution of a sex chromosome^52^. Similarly, the asexual strain of *S. mediterranea* is known to carry a cytologically visible reciprocal translocation between chromosomes 1 and 3^36^ (Chr1/3 translocation, Fig. 1c), which has been proposed as a driver of asexuality, either by disrupting reproductive system development^37^ or by inducing fission behavior^38^. However, without a genome assembly for the asexual strain, both the genomic causes and the consequences of asexuality in the system cannot be elucidated.

Here, we present the first chromosome-scale diploid genome assembly of the asexual *S. mediterranea* strain, alongside extensive population genomic analyses of wild and laboratory populations, to probe its evolutionary history. We demonstrate that three successive inversions and a reciprocal translocation, originating in a sexual ancestor, progressively led to the genome-wide suppression of recombination without directly disrupting reproductive genes. Surprisingly, both laboratory and wild asexual populations exhibit minimal accumulation of deleterious mutations or other expected genomic costs. Occasional sex in the wild and a recent origin of asexuality (0.17–0.4 Ma versus the previously inferred 8.4 Ma) do not fully explain this low mutational load. This raises the possibility that selection processes among adult pluripotent stem cells may purge deleterious variants. Together, our findings support the stepwise evolution of asexuality in S. mediterranea, whereby recombination suppression through structural rearrangements gradually undermines the benefits of sex and facilitates the emergence of fissiparous asexuality to mitigate its costs.

## Results

### Genome assembly and annotation of the asexual CIW4 strain

Toward our goal of examining the genomic causes and consequences of asexuality in *S. mediterranea*, we utilized our previous assembly strategy with PacBio HiFi and Hi-C sequences to generate a high-quality reference assembly of the asexual laboratory strain (CIW4, see Table 1 for details on all strains used in this study). As shown in Table 2, the new assembly (schMedA2) shows a high contiguity with ∼90% of the sequence occurring on the four chromosome scaffolds. A mercury evaluation of the k-mer content with three independent short-read datasets reveals a high base-pair accuracy of the entire assembly (QV of 46.1 and 46.9, of haplotype 1 and 2) and even higher accuracy when only including the chromosome scaffolds (QV of 47.9 and 48.2). Furthermore, the k-mer analysis shows that heterozygosity is well reflected and efficiently phased between the two haplotypes (Supporting Information Genome QC), and the assembly is of similar completeness as our previous planarian genome assemblies^48^ as assessed using Benchmarking Universal Single-Copy Orthologs (BUSCO, Table 2). However, while the assembly is chromosome-scale, it is not telomere to telomere. 88.4 Mb and 90.7 Mb could not be scaffolded in haplotype 1 and haplotype 2, respectively. This level of contiguity is similar to our previous scaffolding success in *S. polychroa* but slightly lower than for the schMedS3 assembly of the sexual laboratory strain^48^. Consequently, the ends of chromosome scaffolds were not capped by telomere repeats. Instead, these repeats were found on unincorporated scaffolds. To investigate the repetitive content, we annotated transposable elements and other highly repetitive regions using EDTA and RepeatExplorer. Similar to the sexual genome, the overall repeat content was high (haplotype 1: 61.7%; haplotype 2: 60.6%) and we were able to resolve large stretches of abundant tandem repeat elements on all Chromosomes. These tandem repeat clusters often spanned > 1 Mb and comprised several distinct repeat sequences, of which some occurred in multiple blocks on multiple chromosomes (Table 3; Tables S1-2; see below). In addition, we improved upon our previous hybrid transcriptome assembly strategy by leveraging the complementary gene annotations from both alleles to annotate the gene content of the genome (See Methods). As is common with planarian genomes, we found that BUSCO completeness increased substantially when running in transcriptome mode (Table 2). Overall, the new *S. mediterranea* genome assembly provides an annotated reference assembly of an asexual planarian strain and a valuable resource for the analysis of the CIW4 laboratory strain as the workhorse of the planarian research community.

**Table 1.** *S. mediterranea* strains, populations, and genome assemblies used in this study. Two laboratory populations with genome assemblies and two wild populations were used. The asexual CIW4 genome assembly (schMedA2) complements the previously published sexual S2F18 assembly.

| **ID** | **genome** | **sample type** | **origin** | **reproduction** | **reference** |
| --- | --- | --- | --- | --- | --- |
| S2F18 | schMedS3 | laboratory | Sardinia | sexual | <sup>48</sup> |
| CIW4 | schMedA2 | laboratory | Barcelona | fission | This study |
| SAR | - | wild population | Sardinia | sexual | This study |
| MEN | - | wild population | Menorca | fission | This study |

**Table 2.** Summary statistics for the genome assemblies and annotations of both *S. mediterranea* CIW4 haplotypes. Quality Value and Base calling accuracy are the average of three independent short-read datasets of the laboratory culture.

| **Genome assembly statistics** | **haplotype 1** | **haplotype 2** |
| --- | --- | --- |
| Quality value | 46.1 | 46.9 |
| Quality value chromosomes | 47.9 | 48.2 |
| Base accuracy (%) | 99.9975 | 99.9979 |
| Base accuracy chromosomes (%) | 99.9984 | 99.9984 |
| Scaffolds (#) | 1120 | 550 |
| Contigs (#) | 1258 | 699 |
| Total length (bp) | 853,691,854 | 836,358,679 |
| Scaffolded (bp) | 765,292,180 | 758,381,961 |
| Scaffolded (%) | 89.6 | 90.7 |
| Unscaffolded (Mb) | 88.4 | 78 |
| GC (%) | 29.66 | 29.49 |
| Scaffold N50 | 272,298,851 | 224,520,763 |
| Contig N50 | 8,105,788 | 7,438,653 |
| BUSCO complete (%)<br>(single copy, duplicated) | 68.1 (62.6, 5.5) | 68.9 (63.1, 5.8) |
| BUSCO fragmented | 9.3 | 9.4 |
| BUSCO missing | 22.6 | 21.7 |
| **Transcriptome statistics** |  |  |
| Genes (#) | 60,931 | 60,595 |
| Transcripts (#) | 104,163 | 103,465 |
| Coding sequences (#) | 44,796 | 44,533 |
| Total gene length (bp) | 355,698,801 | 355,185,528 |
| Total transcript length (bp) | 337,843,474 | 336,471,699 |
| Median gene length (bp) | 791 | 790 |
| Median transcript length (bp) | 1635 | 1633 |
| *longest gene (bp)* | 338,920 | 314,464 |
| *shortest gene (bp)* | 80 | 80 |
| *Gene coverage (%)* | 41.7 | 42.5 |
| *Coding sequence coverage (%)* | 3.4 | 3.5 |
| *BUSCO complete (%)* | 79.1 | 80 |
| *BUSCO fragmented (%)* | 3 | 3.4 |
| *BUSCO missing (%)* | 17.9 | 16.6 |

**Table 3.** The number and size of large tandem repeat elements detected in the asexual (schMedA2) and sexual (schMedS3) genome assemblies. The largest and most abundant repeat consisted of a 158 bp k-mer (158mer) and the second largest repeat consisted of a 159 bp long k-mer (159mer). In addition, we identified four tandem repeats as putative centromeric because they were chromosome-specific and located as expected from karyology (CL66: 424 bp; CL73: 544 bp; CL75: 693 bp; CL79: 766 bp).

| **Genome** | **Tandem repeat** | **N** | **N >1Mb** | **min. (Mb)** | **max. (Mb)** | **total (Mb)** |
| --- | --- | --- | --- | --- | --- | --- |
| *schMedA2*<br>*haplotype 1* | 158mer | 12 | 5 | 0.16 | 2.41 | 11.65 |
|  | 159mer | 3 | 1 | 0.31 | 1.45 | 2.11 |
|  | pericentromeric | 6 | 2 | 0.45 | 1.9 | 6.29 |
| *schMedA2*<br>*haplotype 2* | 158mer | 9 | 3 | 0.26 | 2.54 | 8.56 |
|  | 159mer | 4 | 0 | 0.12 | 0.33 | 1.02 |
|  | pericentromeric | 5 | 0 | 0.11 | 0.67 | 2.18 |
| *schMedS3*<br>*haplotype 1* | 158mer | 14 | 4 | 0.12 | 3.34 | 13.5 |
|  | 159mer | 5 | 3 | 0.72 | 5.31 | 10.67 |
|  | pericentromeric | 6 | 3 | 0.19 | 1.87 | 5.39 |
| *schMedS3*<br>*haplotype 2* | 158mer | 11 | 6 | 0.23 | 2.26 | 12.84 |
|  | 159mer | 4 | 3 | 0.44 | 5.1 | 9.48 |
|  | pericentromeric | 4 | 2 | 0.21 | 1.38 | 3.04 |

### Structural analysis of the asexual genome assembly

Planarian genomes are generally known to undergo rapid structural evolution^48^ and the asexual strain of *S. mediterranea* has previously been shown to carry a reciprocal Chr1/3 translocation^36^. With the asexual schMedA2 genome assembly in hand, we first analyzed its structure in comparison to the sexual strain reference genome assembly (schMedS3;^48^) via Hi-C chromosome conformation capture (Supporting Information: Hi-C QC). The mapping of the asexual Hi-C data on schMedA2 haplotype 1 revealed multiple clear off-diagonal signals as typical signatures of structural rearrangements (Fig. 2a). The off-axis signal between Chr1 and Chr3 (Fig. 2a, solid box 1) was consistent with the known translocation, indicating its successful recovery in the schMedA2 assembly. Two additional off-axis signals (Fig. 2a, solid box 1) suggested large inversions on Chr1 and Chr2 (Fig. 2a, solid boxes 2,3). Interestingly, we further noticed a pronounced increase in contact signal between the centromeres (Fig. 2a, dashed boxes), suggesting centromere clustering typical of a Type I architecture, which has also been observed in the parasitic flatworm *Clonorchis sinensi*^53^. To complement the Hi-C maps, we directly compared the two schMedA2 haplotypes by whole-genome alignment. As expected, the whole-genome alignments reflected the structural differences suggested by Hi-C (Fig. 2b). Specifically, the size and position of the chromosome segments affected by the Chr1/3 translocation matched well with predictions from the literature (Supporting Information: Karyology literature) and large inversions on Chr1, Chr2 and possibly Chr4 were also apparent. However, the haplotype differences on Chr4 were not analyzed further due to its particularly high repeat content and correspondingly lower confidence in its long-range contiguity (Supporting Information: Hi-C Chromosome 4). Surprisingly, our whole-genome alignments revealed that the large inversion on Chr1 was identical with the sexual schMedS3 assembly, indicating that the inversion was already present in the common ancestor of the asexual CIW4 and sexual S2F18 strains (Fig. 2a, Supporting Information Hi-C comparison S2F18 vs CIW4; see below). However, the two additional inversions on Chr1 in the sexual assembly (Fig. 1b) were not detected in the asexual assembly, suggesting continued genome rearrangements in both strains after their divergence (Supporting Information: Hi-C comparison S2F18 vs CIW4). Overall, our new asexual strain reference genome assembly accurately reflects the known Chr1/3 heterozygous translocation and additional genome rearrangements, thereby generalizing our previous findings of extensive structural genome divergence between *Schmidtea* species^48^ to different strains of *S. mediterranea*.

**Figure 2.**
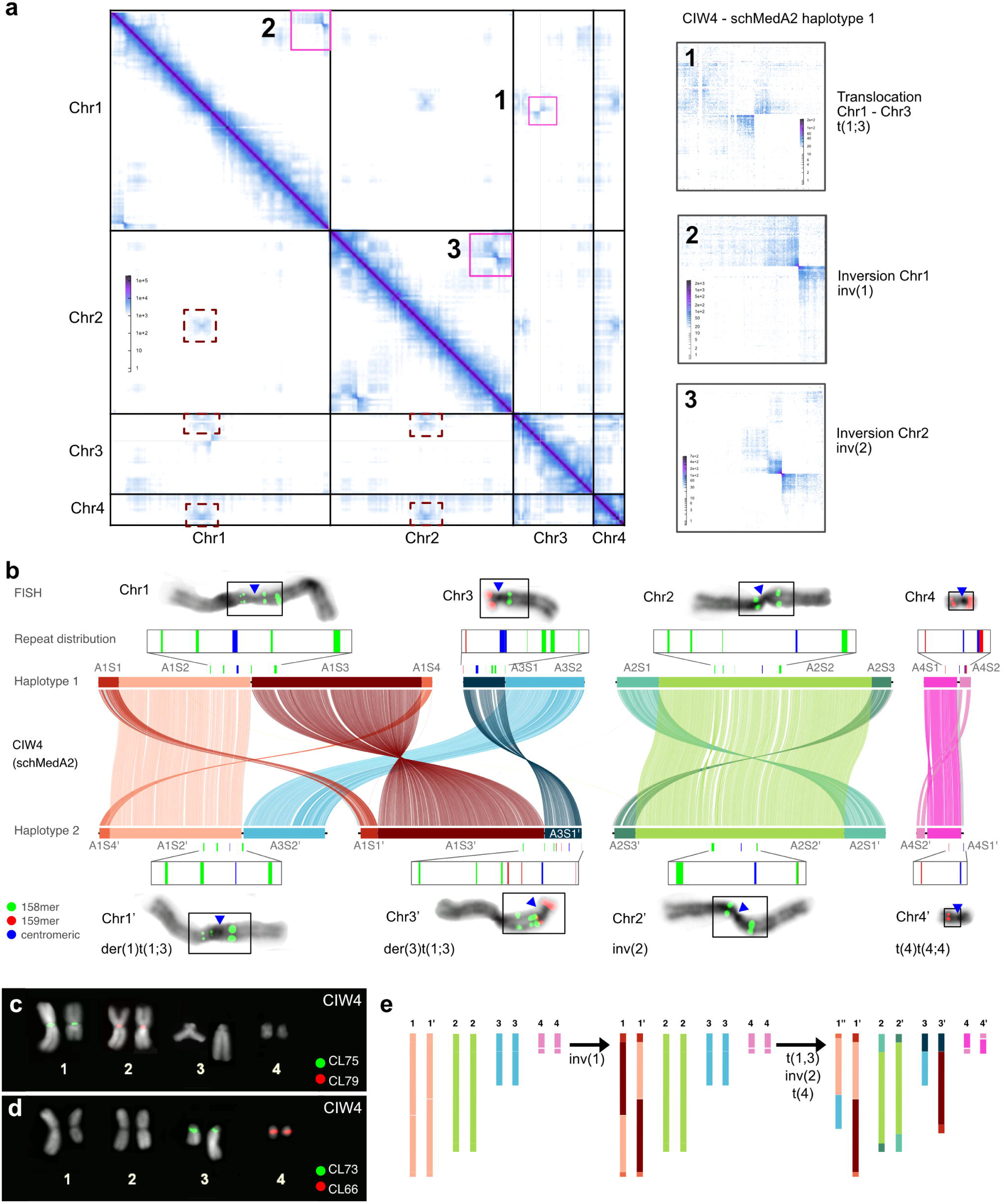
Haplotype-phased genome assembly of the asexual *Schmidtea mediterranea* laboratory strain CIW4. **a** Hi-C chromatin interaction heatmap of haplotype 1 of the schMedA2 assembly of CIW4. The heatmap is symmetric and shows high contiguity within chromosome scaffolds. Boxes in the bottom triangle highlight increased inter-chromosomal contact at the centromeres. Numbered boxes in the top triangle highlight long-distance contacts characteristic of structural variation between haplotypes, and the corresponding numbered panels show these regions at higher resolution. 1) contact between Chr1 and Chr3 due to the Chr1/3 translocation, 2) contact between ends of Chr1 due to a large inversion, 3) contact between ends of Chr2 due to a large inversion. Abbreviations of the structural variations follow the ISCN nomenclature. **b** Comparison between haplotype 1 and haplotype 2 of the schMedA2 assembly. The middle diagram represents the synteny between haplotypes based on whole-genome alignment, showing for each haplotype the genomic strata and their alignment (colored bars and lines connecting them) and the distribution of tandem repeat regions (158mer, 159mer) and putative pericentromeric repeat regions. The top and bottom chromosome images show fluorescent *in situ* hybridization using probes targeting the 158mer and 159mer repeats. Blue triangles indicate the location of the centromeres. See “Supporting Information: FISH” for the original images. Black lines below the syntenic blocks represent the genome sequence. The last row indicates the ISCN description of the derived chromosomes in haplotype 2. **c-d** Fluorescent *in situ* hybridization using directly labeled probes targeting the putative pericentromeric repeats. **e** Schematic representation of the most parsimonious history of chromosome rearrangements. inv(1) likely preceded t(1;3), but inv(2) and t(4) could also have occurred earlier. However, since inv(1) is shared with the sexual strain (S2F18), which does not have these rearrangements, it is most likely that inv(2) and t(4) occurred after inv(1) (Supporting Information: Comparison of Hi-C signals on Chr1).

### Assembly verification by *in situ* hybridization

To independently confirm the various indications of chromosomal rearrangements in the asexual schMedA2 assembly, we thought to exploit the distinct repeat sequences of the large tandem repeat clusters and their predicted spacing in individual chromosomes. *In situ* hybridization (FISH) probes designed against the consensus sequence of two abundant tandem repeats (referred to as 158mer and 159mer) yielded bright and distinct banding patterns on metaphase chromosomes of *S. mediterranea* (Fig. 2b, see Methods). The quantitative analysis of observed versus predicted patterns as a structural bar code in 50 metaphase plates from 10 sexual and asexual specimens each largely confirmed the expected patterns shown in Fig. 2b, despite some variation in signal intensity due to variable chromosome compaction. Specifically, the weaker 158mer signal on Chr1 compared to Chr1′ (Supporting Information: Chromosome FISH) and higher signals for both the 158mer and 159mer on Chr3′ confirmed the mapping of the Chr1/3 translocation. In addition, the FISH patterns confirmed the predicted loss of 159mer signal on Chr4′, which together confirms the structural accuracy of the schMedA2 assembly.

During the course of these analyses, we noticed large tandem repeat clusters at the approximate position of the centromeres of each of the four chromosomes, but interestingly, the repeat sequence was different for each chromosome (Fig. 2b; Table 3). FISH against each of the four repeats indeed resulted in bright labelling in the vicinity of the centromere, yet with remarkable specificity for individual chromosomes (Fig. 2c-d). While further studies will be necessary for the functional definition of centromere sequences in *S. mediterranea*, the chromosome-specific pericentromeric repeats enabled us to link genome sequences to corresponding chromosomes and provided additional experimental confirmation of the long-range contiguity of the schMedA2 genome.

Accordingly, schMedA2 haplotype 1 contains chromosomes similar to chromosomes of the sexual S2F18 strain (Chr1-4), while haplotype 2 contains rearranged chromosomes with the reciprocal balanced Chr1/3 translocation resulting in a shorter derived Chr1’ and a longer derived Chr3’. On the basis of sequence homology alone, the rearranged Chr3’ would align with Chr1 due to the translocated long arm of Chr1, which illustrates the utility of the unique pericentromeric regions in understanding the structural genome organization. Furthermore, haplotype 2 contains an inverted Chr2’ and a derived Chr4’ due to a transposition of a distal region and loss of a 159mer repeat region (Fig. 2b). As previously noted, Chr4’ is misscaffolded in the current assembly and further efforts such as ultra-long sequencing will be needed to resolve the correct structure of Chr4’. Our data suggest the succession of chromosomal translocations and inversions depicted in Fig. 2e as the most parsimonious evolutionary explanation for the current asexual genome structure. The depicted occurrence of the Chr1 inversion before the Chr1/3 translocation is the most parsimonious explanation, as alternative scenarios would require three interchromosomal translocations instead of one. This finding is conceptually important, as it links the evolution of the asexual CIW4 strain to the gradual accumulation of genomic rearrangements (see discussion).

### Translocation breakpoint analysis

Given previous speculations that the reciprocal translocation between Chr1 and Chr3 might be causally linked to the loss of reproductive system development in the asexual strain^37,38^, we next examined the breakpoints in more detail. Using Hi-C mapping and genome alignment, we identified the breakpoints in the asexual schMedA2 assembly and located the homologous regions in the sexual schMedS3 assembly. Interestingly, in both assemblies, the breakpoints clearly map to tandem repeat clusters. In the schMedS3 assembly, the breakpoint region spans a 1.9 Mb region consisting of 158mer repeats on Chr1 and a 1.4 Mb region of 158mer repeats on Chr3 (Fig. 3a–b; Supporting Information: Strata liftover). After identifying the breakpoint, we selected haplotype 1 of schMedS3 as our reference, allowing for direct comparison of its synteny with haplotype 2 and schMedA2 via whole-genome alignment. As expected, schMedS3 haplotype 2 aligned in inverted orientation across the repeat stretch on Chr1 (Fig. 3a, first row). Similarly, schMedA2 haplotype 1 aligned across the breakpoint, although downstream sequences were on unincorporated scaffolds, likely due to a flanking ∼250 kb low-mappability region. The alignment of schMedA2 haplotype 2 revealed the Chr1/3 translocation with a transition from Chr1‘ to unincorporated scaffolds and then to Chr3‘. On Chr3, alignments around the breakpoint were simpler, with clean connections across the breakpoint for schMedS3 haplotype 2 and schMedA2 haplotype 1, and a switch from Chr3’ to Chr1’ in schMedA2 haplotype 2 due to the Chr1/3 translocation (Fig. 3b). Having narrowed the Chr1/3 translocation breakpoint to the 158mer repeat region, we examined the repeat variation between haplotypes by similarity heatmaps with 2 kb resolution to further home in on the precise location of the break point. As shown in Fig. 3c, we detected strong self-similarity between the repeat regions on Chr1 and Chr3. On the rearranged Chr1’, two blocks were apparent: the upstream block was more similar to Chr1 and the downstream block more similar to Chr3, indicating that the Chr1/3 translocation occured at the border between these regions (Fig. 3c) and hence deeply within the Mb-scale gene deserts of the 158mer tandem repeat clusters on Chr1 and Chr3.

**Figure 3.**
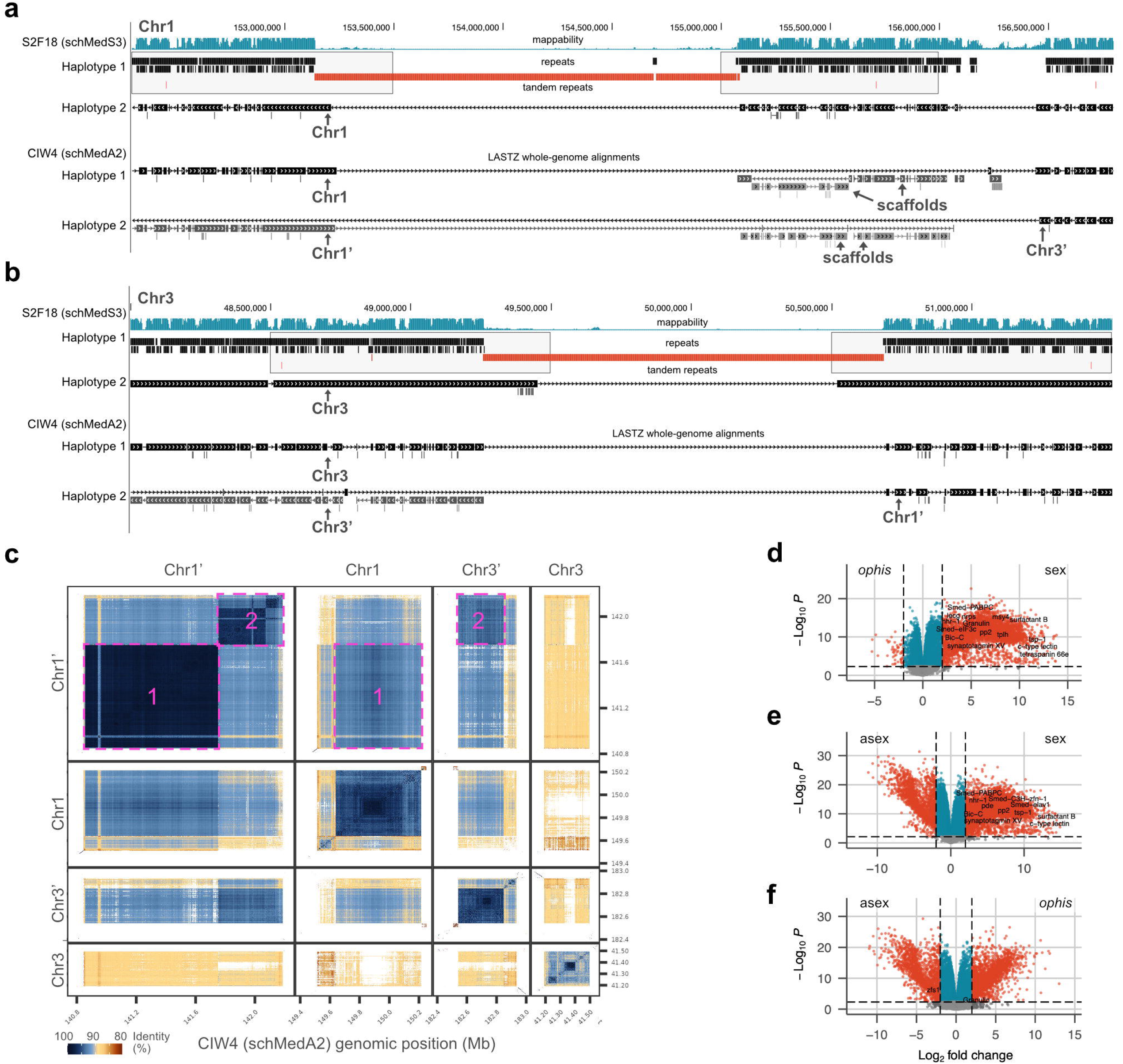
Genomic context of the Chr1/3 translocation. **a-b** UCSC genome browser views with haplotype 1 of the schMedS3 genome assembly of the sexual laboratory strain S2F18 as reference. Tracks show genome mappability, repeat content, and whole genome alignment tracks of the region identified as homologous to the breakpoint in the asexual schMedA2 assembly on (a) Chr1 and (b) Chr3. Note that the apparent extension of the Chr1’ alignment across the breakpoint in (a) reflects netting of repetitive elements (2x BURRO, 1x RND-4-family) rather than true synteny. Grey boxes outline the regions evaluated for gene loss and/or changes in gene regulation in Supporting Information: Gene loss. **c** Similarity heatmap of 158mer repeats in the schMedA2 assembly showing the high similarity within and between the breakpoint regions. Each box shows similarity in 2kb windows of regions to themselves and between them. Numbered boxes highlight increased similarity of the Chr1’ repeats with (1) Chr1 and (2) Chr3’, indicating that the translocation breakpoint lies between them. **d-f** Volcano plots showing differential expression analysis used to identify reproduction-related genes by comparing (d) sexual control (wild-type & eGFP RNAi) vs. *ophis* RNAi, (e) sexual control vs. asexuals wild-type, and (f) *ophis* RNAi vs asexual wild-type. The 2,136 high-confidence reproduction-related genes are defined as those that are significantly upregulated in sexual control in both (d) and (e).

To validate the lack of gene loss, we examined genes near the breakpoints using the annotation tool TOGA^54^ (Supporting Table S3). We also identified high-confidence reproduction-related genes near the breakpoints by generating gene expression profiles from sexual and asexual wild-type animals, as well as from RNAi-treated sexual animals (eGFP control and *ophis* RNAi), the latter of which blocks early reproductive system development^55,56^ (Fig. 3d-f, Supporting Information: Reproduction-related genes; Supporting Tables S4-8). We found no clear coding gene losses within 1 Mb of the Chr1/3 breakpoints, nor were reproduction-related genes located within 250 kb of them (Supporting Table S9; Supporting Information: Gene Loss). Likewise, no coding gene losses or nearby reproduction-related genes were detected around the large inversions on Chr1, and only a single exon loss was observed in one gene in CIW4 haplotype 2 due to the Chr2 inversion (Supporting Table S9). These results suggest that the large-scale structural rearrangements themselves did not disrupt coding genes. Thus, the genomic basis of asexuality in *S. mediterranea* likely lies among the many other genetic and epigenetic differences between the strains, offering a promising direction for future research.

### Genomic costs of asexuality

Next, we explored the possible genomic consequences of asexuality in the schMedA2 assembly. Reduced purifying selection is a genomic signature frequently associated with the loss of recombination^13–19^, which can be assayed by measuring the ratio of non-synonymous (dN; amino-acid changing) to synonymous (dS; non-amino acid changing) substitutions in genes. We inferred pairwise d_N_/d_S_ for both the asexual and sexual strains of *Schmidtea mediterranea* in comparison with each of the three other *Schmidtea* species, based on 10,479 single-copy genes (see Methods; Fig. 4a; Supporting Table S10). Specifically, for each sister species, we compared d_N_/d_S_ of the asexual strain to that of the sexual strain to assess whether the two strains show differences in the strength of purifying selection.

**Figure 4.**
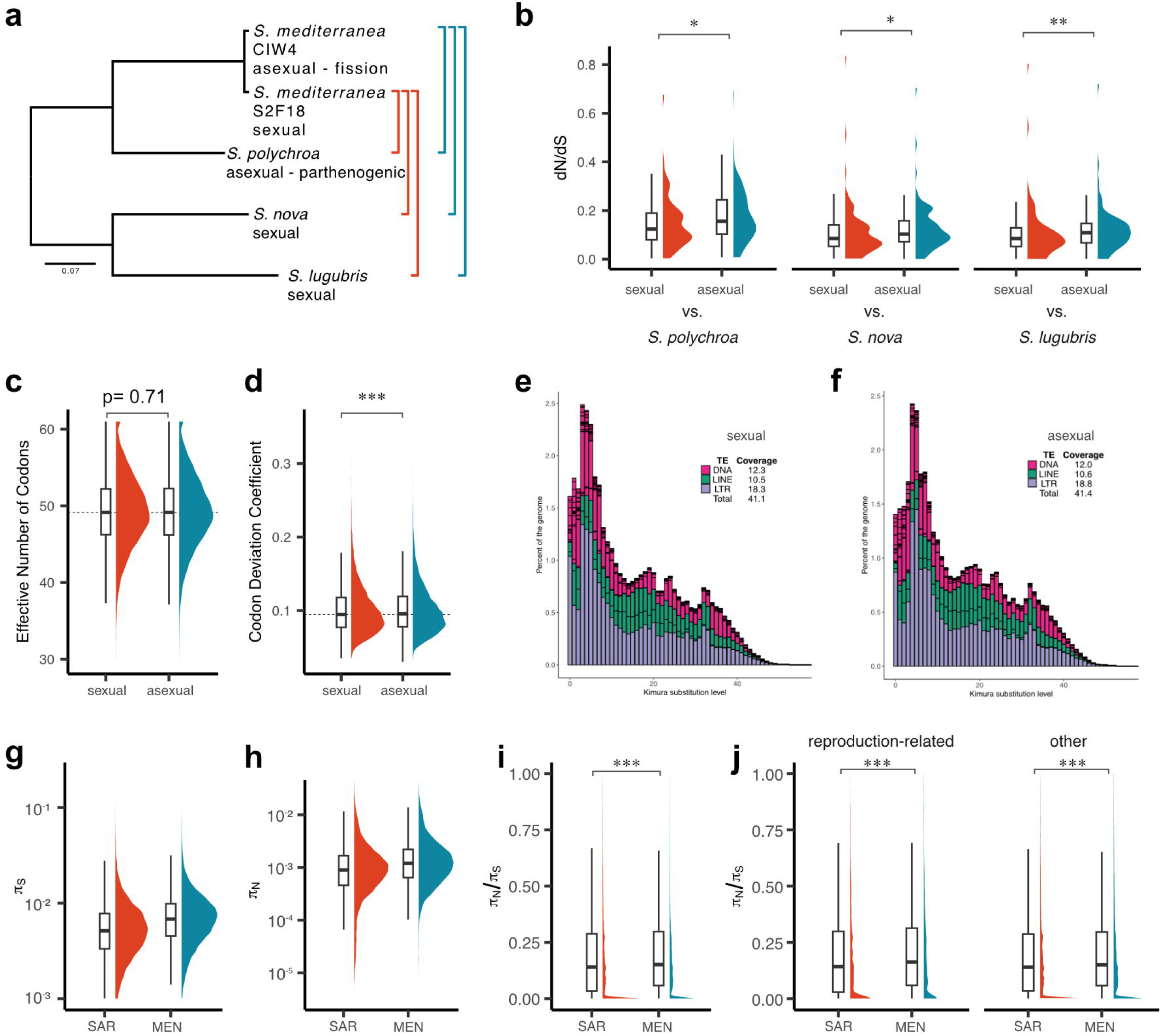
Detection of costs of asexuality in *S. mediterranea*. **a** Maximum-likelihood phylogeny of the genus *Schmidtea* used for dN/dS analysis. The phylogeny is based on codon alignments of 10,556 single-copy genes (16,684,518 positions–no missing data) and was inferred in IQ-TREE using gene-wise partitions and best-fit models from ModelFinder. All nodes show maximal support (500 bootstraps). Colored brackets indicate pairwise comparisons in (b). **b** Pairwise d_N_/d_S_ comparisons for 89 genes with sufficient divergence (dS > 0.1), between the sexual (S2F18) and asexual (CIW4) strains. Asterisks denote significant differences; d_N_/d_S_ was slightly elevated in CIW4 across all comparisons. **c** Effective number of codons (ENC), a measure of codon usage bias (range: 20 = maximum bias to 61 = no bias), did not differ significantly between strains. d Codon deviation coefficient (CDC), quantifying bias relative to background composition, was significantly higher in S2F18, suggesting slightly stronger selection in sexuals. e-f Kimura distance landscapes of repeats in S2F18 (**e**) and CIW4 (**f**), showing repeat copy number (y-axis) vs. divergence from consensus (x-axis). Lower distances indicate recent TE activity; higher values reflect older elements. g-j Tests for costs of asexuality using whole-genome sequencing of 49 individuals from wild asexuals from Menorca (20 x MEN) and wild sexuals from Sardinia (29 x SAR). g Nucleotide diversity (π) at synonymous codon positions. h π at non-synonymous codon positions. i Ratio of π at non-synonymous and synonymous codon positions (πN/πS) of 12,470 genes compared between SAR and MEN. πN/πS was significantly larger in MEN, indicating reduced purifying selection in asexuals. j πN/πS of all genes from g split into 1,507 high-confidence reproduction-related genes and the 10,963 remaining genes. Boxplots show interquartile range (IQR) and the median. Whiskers extend to 1.5 × IQR. Statistical results are in Table S12. All statistical tests were two-sided.

The pairwise d_N_/d_S_ values of both sexual and asexual *S. mediterranea* compared to each of the three other *Schmidtea* species were almost all <<1, consistent with the general conservation of protein sequence by selection (Supporting Table S11). Interestingly, d_N_/d_S_ in the asexual strain was consistently higher than in the sexual strain, indicating less efficient selection and thus a potential cost of asexuality. However, the effect size was extremely small (Cohen’s *d* = 0.01, Supporting Table S12), suggesting that while statistically significant, the difference is biologically negligible. Furthermore, we noted that synonymous divergence (d_S_) between the sexual and asexual strains was generally low, which results in low power to detect differences in d_N_/d_S_ (Supporting Table S11). When applying a common filter to retain genes with d_S_ > 0.1 between the sexual and asexual strains, only 89 genes were retained. These genes also had high d_S_ across the genus, all being among the top 15% of d_S_. Although we found no significant GO-term enrichment of these genes, they were significantly enriched for genes on our previously defined reproduction-related gene list (Fig. 3d-f; Supporting Information: Reproduction-related genes; odds ratio = 2.22, 95% CI: 1.3–3.7, p = 0.003, two-sided Fisher’s exact test), which is expected since reproduction-related genes are known to evolve at an accelerated rate^57,58^. When repeating the d_N_/d_S_ analysis with this subset, we found that the increase in d_N_/d_S_ in asexuals was slightly larger (Cohen’s *d* = –0.18), again hinting at less efficient purifying selection in the asexual strain (Fig. 4b; Supporting Table S12).

Further genomic signatures of asexuality can include changes in codon usage bias due to reduced purifying selection^59^ or the proliferation of repetitive elements due to the lack of recombination^60^. Interestingly, neither metric was significantly altered in the asexual genome assembly: Two independent measures of codon usage bias detected no or negligible differences between the two genomes (Fig. 4c–d; Supporting Tables S12-13). In addition, the global transposon landscapes of the two strains were highly similar, suggesting that the emergence of asexuality was not associated with overt transposon proliferation (Fig. 4e–f). In conclusion, our genome-wide comparisons revealed surprisingly few costs of asexuality in the CIW4 laboratory strain, which is all the more surprising in the face of the current estimate of the split between those two lines of ∼ 8.4 Ma^39^.

### Population genomics of natural *S. mediterranea* populations

Since the generally low divergence between the sexual and asexual laboratory strains restricted the use of d_N_/d_S_ to only a few genes, we instead chose to elucidate the strength of purifying selection using the distribution of segregating sites in natural populations. Because the source population of the asexual laboratory strain in Barcelona has likely become extinct (pers. observations), we sampled the asexual populations on Menorca, which were described to comprise distinct diploid and triploid populations^37,38,40^. We successfully collected the diploid population (MEN), but the original locality had dried up due to agricultural use. We only found *S. mediterranea* in a concrete canal further downstream, suggesting that this population may be threatened in the wild. In addition, we collected wild sexual *S. mediterranea* from Sardinia (SAR) and performed short-read whole-genome sequencing at 30x coverage of 49 individuals across both populations (20 x MEN, 29 x SAR). We supplemented these data with equivalent sequencing samples from the two laboratory populations (3 x CIW4, 3 x S2F18). Principal component analysis and k-means clustering of genotype likelihoods of the sequencing data revealed that the laboratory strain CIW4 was highly similar to MEN and the S2F18 laboratory population clustered with SAR, consistent with the origins of the founder animals^35^ (Supporting Information: Population Genomics). To further corroborate these results, we genotyped SAR, MEN, and laboratory individuals at the COI barcoding locus and compared the sequencing reads with publicly available data^39^. A phylogenetic analysis showed MEN specimens clustering with CIW4 and SAR specimens grouping both with COI isolates from Corsica/Sardinia, but also showing some similarity to Sicily and Tunisia, albeit with low node support (Supporting Information: Population Genomics). This again indicates that MEN and CIW4 are very similar. Therefore, the MEN and SAR population genomics data sets are well-suited for complementing our previous results with the laboratory strains.

To investigate the costs of asexuality in the natural populations, we first inferred nucleotide diversity (π) within our MEN and SAR population genomics data at each nucleotide position and identified 12,470 protein-coding genes that had sufficient diversity in both populations for the following analysis (Supporting Table S14; Note, that these do not completely overlap with the single-copy genes, See Supporting Information: Dataset intersection). We then calculated the ratio of the diversity at non-synonymous (π_N_) and synonymous (π_S_) positions. The interpretation of the π_N_/π_S_ ratio is similar to d_N_/d_S_, with an expectation for the ratio to be below 1 for most proteins in the presence of efficient purifying selection. Indeed, we found that the likely sexual SAR population had significantly lower π_N_ and π_S_ compared to the likely asexual MEN population (Fig. 4g-h), yet again with a negligible effect size across the complete gene set (Cohen d: -0.053, Fig. 4i; Supporting Table S12). However, when partitioning the dataset based on our reproduction-related gene annotation (Fig. 3d-f), we again obtained higher π_N_/π_S_ values in MEN as compared to SAR, but this time with a 50% larger effect size (Cohen d: -0.073, Fig. 4j; Supporting Table S12). Our results therefore confirm the expected reduced purifying selection in the wild asexual population, especially in sexual development genes that are likely experiencing relaxed selection in asexuals. However, the effect sizes were still modest, which indicates that even wild asexual *S. mediterranea* are not in the process of undergoing mutational meltdown.

### Population history of asexuality in *S. mediterranea*

One plausible explanation for the unexpectedly low cost of asexuality in *S. mediterranea* could be rare events of sexualization and sexual reproduction within wild asexual populations^61,62^. In fact, a case of sexualization of asexual *S. mediterranea* has been reported^63^ and seasonal switching between fissiparous and sexual reproduction is common in planarian species, including the sister genus *Dugesia*^64,65^. To test for the possibility of rare sex in the wild MEN population, we analyzed linkage disequilibrium (LD) as a function of genomic distance. The resulting LD decay curves can reveal recombination events in the form of breakdown of the associations between distant genetic variants^66^. While the exact shape of the LD curves can be influenced by selection, demography, and drift^67,68^, asexual populations without recombination are expected to display high LD regardless of distance^66^, but rapid distance-dependent decline can be expected in recombining sexual populations (Fig. 5a). Consistent with this prediction, the LD decay pattern in MEN shows little distance-dependent decay, supporting strict asexuality, while LD decay in SAR is steep as expected in a sexual population (Fig. 5b). Furthermore, the SAR population exhibited a higher inbreeding coefficient indicative of high inbreeding (Fig. 5c). Therefore, the MEN population is likely truly asexual and unlikely to undergo periodic events of sexualization.

**Figure 5.**
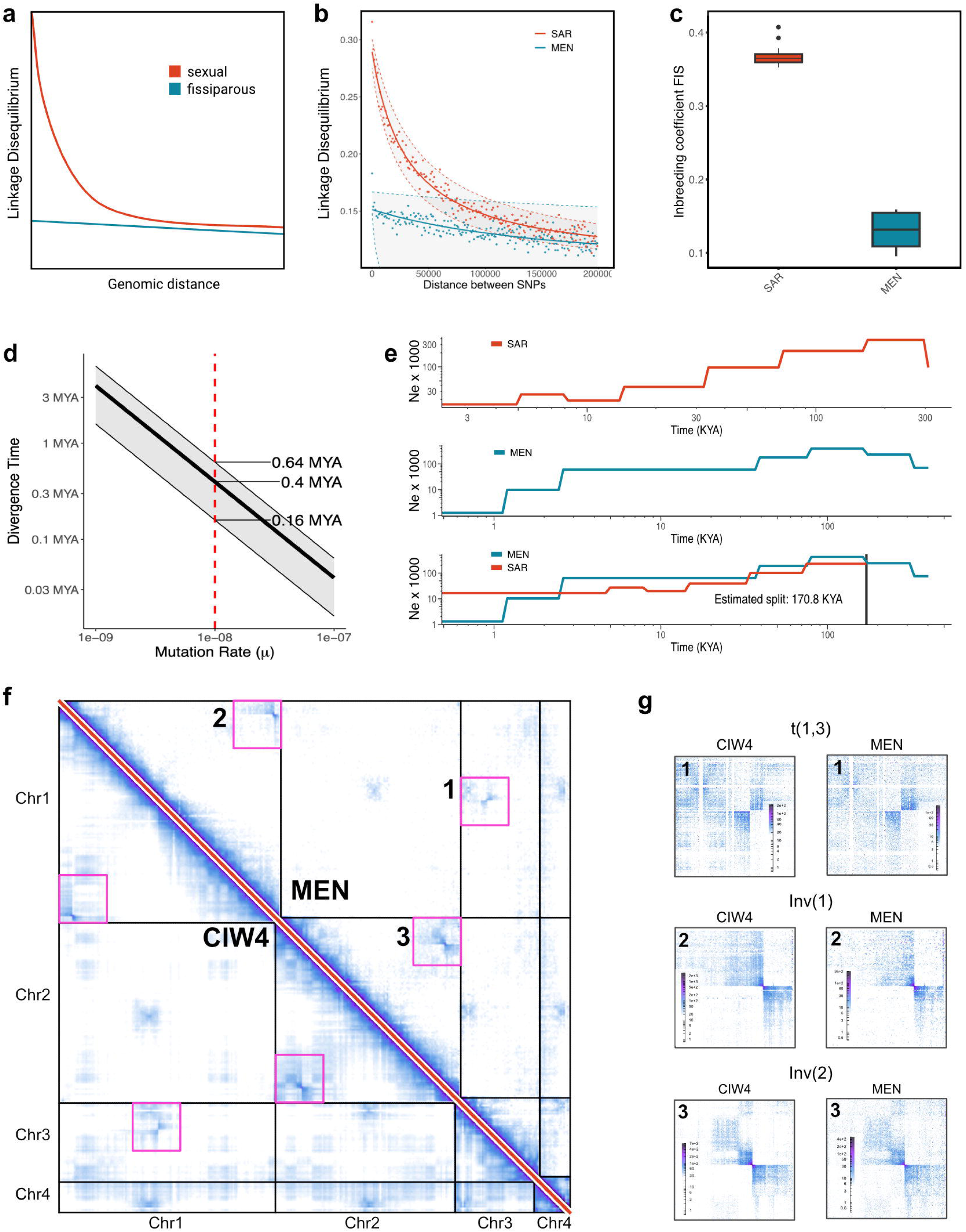
Sexual-asexual divergence estimation and evidence for persistent asexuality in wild *S. mediterranea*. **a** Schematic representation of the expected LD decay patterns in a sexual or asexual population. LD should drop sharply with distance in sexual populations, but be largely independent of genetic distance in asexual populations. **b** LD decay in wild asexual Menorca (MEN) and wild sexual Sardinia (SAR) populations inferred using ngsLD. Envelopes represent 95% confidence intervals based on 100 non-parametric bootstraps. **c** Inbreeding coefficient of SAR and MEN populations showing higher inbreeding in SAR. **d** Maximum-likelihood divergence estimation based on 1,961,630 four-fold degenerate positions across 10,556 single-copy proteins. The red line indicates the empirical estimate of per-generation mutation rate in *S. mediterranea* (Guo et al. 2020). The diagonal line and confidence envelope represent average divergence rate and minimum and maximum divergence μ across a range of generation times (1-4 generations/year). **e** Sexual-asexual divergence estimation using pairwise coalescent modeling of MEN and SAR population. **f** Hi-C matrix on haplotype 1 of the asexual schMedA1 assembly for CIW4 (bottom triangle) and MEN (top triangle), highlighting the shared structural variation between MEN and CIW4. Pink boxes outline off-diagonal signals shown in detail in (g). **g** Details of the Hi-C signal of (1) the Chr1/3 translocation, (2) the Chr1 inversion, and (3) the Chr2 inversion. In all cases, the Hi-C signal of CIW4 and MEN closely match. The signal was also qualitatively identical between three clonal lines generated from Menorca individuals (Supporting Information: Hi-C Menorca isolines). Boxplots indicate the interquartile range (IQR) and the median. Whiskers extend to 1.5 × IQR.

A second plausible explanation for the apparent low genomic costs of asexuality in *S. mediterranea* could be a more recent origin of the asexual strain than the previous estimate of ∼8.4 Ma, which was based on a two-step dating approach, with the initial calibration point based on the biogeographical split between the African and South American continents^39^. Instead of a biogeographical calibration, we used a strict molecular clock to estimate the divergence time between sexual and asexual strains. We estimated neutral divergence rates based on 1,961,630 four-fold degenerate sites in 10,556 single-copy genes and applied the formula *t = k/μ*, where *t* is the number of generations since divergence, *k* is the number of neutral substitutions per site, and *μ* is the mutation rate per site per generation. Using a pedigree-based estimate of μ = 1×10⁻⁸ for the sexual laboratory strain S2F18^52^ and assuming 1–4 generations per year, we estimated the sexual-asexual divergence at ∼0.4 Ma (0.16–0.64 Ma). Allowing two orders of magnitude variation in μ widened the estimate to 0.01-3.4 Ma (Fig. 5d). In addition, we estimated divergence using Pairwise Sequentially Markovian Coalescent (PSMC) modeling, which infers historical changes in effective population size from genome-wide polymorphism patterns. Using all genomes from the MEN and SAR populations and again assuming μ = 1×10⁻⁸, we estimated the split at ∼0.17 Ma (Fig. 5e). Finally, we estimated the median age of the last independent LTR transposon insertion to be 0.21 Ma in the asexual genome and 0.42 Ma in the sexual genome, adding another line of evidence to the recent split (Supporting Information: LTR insertion age). Therefore, three divergence time estimates independently suggest a relatively recent split between the asexual and sexual strains, which is also consistent with the k-means clustering analysis grouping MEN and CIW4 together (Supporting Information: Population genomics).

### Impact of chromosomal rearrangements on genetic diversity

A recent sexual-asexual divergence is especially surprising given the broad geographic distribution of the asexual strain across the Balearic Islands and mainland Spain, and the genome-wide structural divergence between the lineages. Therefore, we explored if the observed chromosomal rearrangements were shared between MEN and CIW4, and if they might be linked to the origin of asexuality. To achieve this, we generated clonal isolines from individual MEN animals by cutting and regeneration and performed Hi-C sequencing on them. The Hi-C signal of four biological replicates from three isolines was highly similar, indicating a shared chromosome organization. When comparing MEN to CIW4, we found that they shared the breakpoints for the Chr1/3 translocation, the Chr1 inversion, and the Chr2 inversion (Fig. 5f-g). This again confirms the remarkably high similarity between the Menorca and Barcelona strains and provides important confirmation that the many chromosomal rearrangements in the schMedA2 assembly are not an artefact of laboratory culture.

To ask how the chromosomal rearrangements impact genetic diversity in the wild populations and to obtain hints at the mechanisms by which asexuality might have evolved, we inferred the population mutation rate θ for MEN and SAR using the diversity estimator θ_π_ and the Watterson estimator θ*_W_* across the genome in 50kb windows. We found that for both estimators, genome-wide genetic diversity was significantly higher in MEN than in SAR (Fig. 6a-b, both p-values < 2.2e-16, Supporting Information: Population genomics), consistent with the expected independent divergence of the two haplotypes in an asexual population. Interestingly, we found a difference in diversity estimates between chromosomes. In SAR θ_π,_ diversity was significantly higher in Chr1 but not different between the other chromosomes, as expected if the Chr1 inversion would repress recombination (Fig. 6c). An even more striking pattern was observed in MEN: The highest diversity was observed on Chr1, followed by Chr2 and Chr3, which had similarly high diversity, and Chr4 showed substantially lower diversity, approaching the range observed in SAR (Fig. 6c, similar results obtained using the θ*_W_* estimator, Supporting Information: Population genomics). Finally, we found that observed heterozygosity was high in MEN and CIW4 compared to SAR, and the lowest in the inbred S2F18 laboratory strain (Fig. 6d). This observation is consistent with the substantial structural variation on Chr1, Chr2, and Chr3. Taken together, these findings suggest a link between structural rearrangements and elevated genetic diversity, potentially driven by recombination suppression.

**Figure 6.**
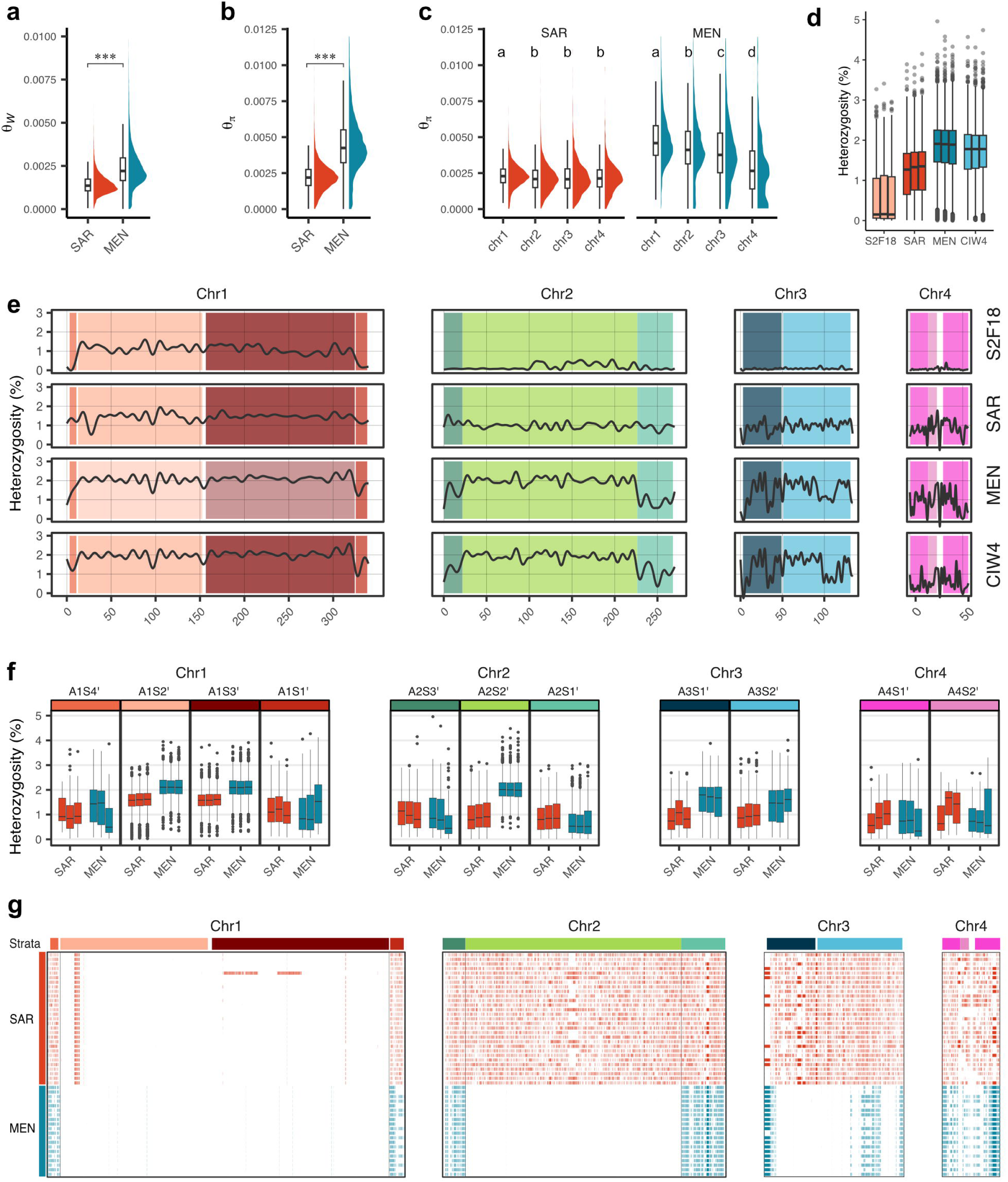
Genetic diversity and individual heterozygosity assessed using whole genome sequencing of asexual (Menorca, MEN) and sexual (Sardinia, SAR) populations. All analyses used the sexual schMedS3 genome assembly as a reference. **a** Population mutation rate (θ = 4N_e_μ) estimated using the Watterson’s estimator (θ_W_), which is based on the number of polymorphic sites. θ_W_ is significantly higher in MEN compared to SAR. (p < 0.001, Mann-Whitney U test). **b** Population mutation rate estimated using nucleotide diversity (θ_π_), which is based on the average number of pairwise differences between individuals. θ_π_ is significantly higher in MEN compared to SAR. (p < 0.001, Mann-Whitney U test). **c** Chromosome-specific θ_π_ estimates for SAR and MEN populations. Different letters indicate significant differences between groups (all p < 0.001, pairwise Wilcoxon rank-sum tests with Bonferroni correction). **d** Percentage of heterozygous sites among all callable sites, calculated in 50kb windows across three representative samples from each population (SAR and MEN). Box plots show the distribution of heterozygosity levels across the genome. **e** Average heterozygosity patterns from panel (d) displayed by chromosome (Chr1-4). **f** Heterozygosity levels across genomic strata in each chromosome. Box plots show the distribution of heterozygosity within each defined genomic region. Colored bars above indicate genomic strata inferred via alignment to the asexual genome schMedA2 haplotype 2 (See Fig. 1). **g** Runs of homozygosity (ROH) on the sexual reference genome inferred for 29 SAR and 20 MEN individuals. Each row represents one individual. ROH are shown as colored segments. Colored bars above indicate genomic strata inferred via alignment to the asexual genome schMedA2 haplotype 2 (See Fig. 1). All statistical tests were two-sided.

To investigate how individual heterozygosity was related to structural rearrangements, we examined heterozygosity along the chromosomes. We found that, across all strains, Chr1 consistently showed higher heterozygosity, whereas the other chromosomes were more homozygous in the highly inbred S2F18. Mapping the genomic strata from CIW4 revealed that regions of elevated heterozygosity in MEN and CIW4 aligned with the identified inversions and translocations (Fig. 6e). Specifically, MEN showed higher heterozygosity than SAR within inversions on Chr1 (strata A1S2’ & A2S3’), the inversion in Chr2 (strata A2S2’), and Chr3 (A3S1’ & A3S2’) which was affected by the Chr1/3 translocation (Fig. 6f, Supporting Information: Population genomics). These elevated levels of heterozygosity suggest suppression of recombination specifically in those regions, but recombination in other parts of the genome (i.e., signatures of sexuality). To understand this further, we inferred runs of homozygosity (ROH), which are a tell-tale sign of inbreeding, for SAR and MEN. SAR had almost no ROH in the Chr1 inversion but many ROH across all other parts of the genome, suggesting heavy inbreeding. In contrast, ROH in the MEN population were uniform between individuals and restricted to genomic regions not implicated in structural rearrangements (Fig. 6g), suggesting these ROH tracks were inherited from a sexual ancestor rather than the result of recent accumulation under asexuality (Fig. 6g). Overall, our results show that the population history of asexual *S. mediterranea* around the Mediterranean is intimately linked to the accumulation of chromosomal rearrangements and the associated suppression of recombination.

## Discussion

Here, we investigate the emergence of asexual reproduction in the planarian model species *S. mediterranea* from a comparative genomics perspective. The high-quality, haplotype-phased schMedA2 genome assembly of the CIW4 laboratory strain represents the first genome of an asexually reproducing planarian, enabling analysis of the genomic causes and consequences of asexuality. Surprisingly, our results reveal only weak traces of the predicted genomic consequences of asexuality, e.g., reduced purifying selection, changes in codon bias or transposon accumulation (Fig. 4b-j), despite obligate asexuality even in wild populations (Fig. 5b). Our results suggest asexual and sexual lineages split much more recently than the previously suggested ∼ 8.36 Ma^39^ and reveal a history of chromosomal inversions and translocations that indicates a role of chromosomal dynamics in sexual system evolution.

The previous age estimate for the divergence between the asexual and sexual strains was based on a strict molecular clock applied to a 309 bp fragment of the COI gene, calibrated using the Gondwanaland split (∼100 Ma) as a biogeographic event. Although fossil-based calibrations are generally more reliable, the complete absence of a fossil record in planarians makes this unfeasible^46^. With the availability of the *schMedA2* assembly and a pedigree-based estimation of the germline mutation rate in *S. mediterranea*^52^, we could reexamine this question using genome-wide data. A phylogenetic approach based on ∼2 million four-fold degenerate sites yielded an age estimate of 0.4 Ma (Fig. 5d). Coalescence modeling, which infers divergence from patterns of genetic variation across both coding and non-coding regions, produced a slightly younger estimate of 0.17 Ma (Fig. 5e). A third approach, based on the neutral accumulation of transposon insertions, estimated divergence between 0.21 and 0.42 Ma (Supporting Information: LTR insertion age). Finally, Guo et al. estimated the origin of the large inversion on chromosome 1 to be ∼0.32 Ma^52^. We find this inversion to be shared between the sexual and asexual strains (Fig. 5f–g; Supporting Information: HiC comparison S2F18 vs CIW4), further constraining the divergence time and contradicting earlier marker-based phylogenies that placed the asexual lineages as a sister group to all sexual lineages^39^. Together, these four orthogonal approaches converge on a much younger divergence between the asexual and sexual strains than previously thought, challenging the prevailing microplate tectonics model for the evolution and dispersal of *S. mediterranea* around the Mediterranean basin^38,39^.

Instead, our results are consistent with the following two population history scenarios. In the “island scenario”, sexual strains bearing the Chr1 inversion first colonized Corsica and Sardinia, then dispersed to the Balearic Islands. There, factors such as small effective population sizes and inbreeding may have favored additional chromosomal rearrangements and the evolutionary emergence of asexuality, followed by a recent migration to the mainland (Fig. 7a). The “ghost population scenario” in contrast, posits a now-extinct coastal ancestor that harbored the original inversion. Its descendants would have split into the Corsican/Sardinian and Spanish coastal populations, with the latter acquiring further rearrangements and asexuality before reaching the Balearic Islands (Fig. 7b). We favor the island scenario since the formerly known mainland asexual populations were found in artificial habitats (e.g., a fountain in Barcelona and irrigation canals in Girona), whereas the populations on Menorca were in more natural streams and wetlands. This makes anthropogenic dispersal, possibly via ornamental aquatic plants, from Menorca to the mainland more plausible than the reverse and thus favors the island scenario. Furthermore, the presence of both diploid and triploid populations on Menorca^37,40^, contrasting with exclusively diploid mainland populations, also suggests the origin of mainland populations from Menorca. Taken together, our results indicate that *S. mediterranea* has dispersed from Sardinia to the Balearic Islands and subsequently to the Spanish mainland, explaining the low genetic distance between asexual strains.

**Figure 7.**
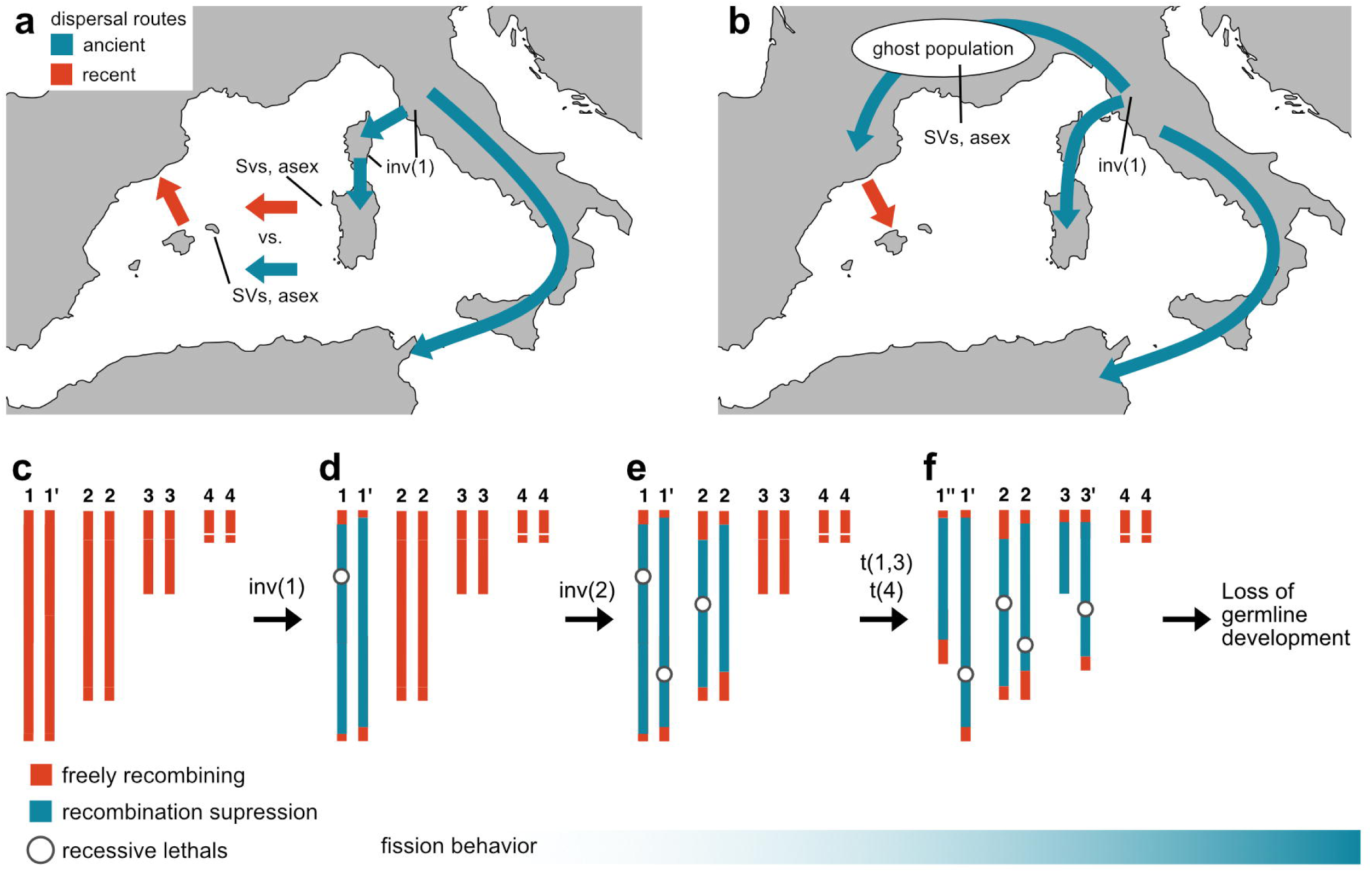
Biogeographical scenarios and evolutionary model for the origin and spread of asexuality in *S. mediterranea*. **a** Island scenario: The Chr1 inversion, inv(1), occurred on Corsica or the mainland, then other structural variants (SVs) occurred followed by the origin of asexuality (asex). If dispersal to Menorca was recent (red arrow) then SVs and asexuality likely originated on Sardinia, but if dispersal was ancient (blue arrow), it is more likely that they originated on Menorca. **b** Ghost population scenario: inv(1) originated in a now-extinct ghost population on the Italian mainland. This population subsequently migrated along the Mediterranean coast and dispersed the Balearic Islands, carrying both the other SVs and asexual reproduction. **c-f** Proposed model of evolution of asexuality in *S. mediterranea*. **c** Ancestral genome without heteromorph structural variations and no recombination suppression. **d** Inversion inv(1) on Chr1 causing recombination suppression and establishment of a lethal or balanced lethal system through capture of deleterious alleles or *de novo* mutations. **e** Secondary inversion inv(2) on Chr2, extending recombination suppression to >60% of the genome. **f** Reciprocal translocation t(1,3) and additional rearrangements affect nearly the entire genome through heteromorphic changes. Homologous chromosome pairing becomes increasingly difficult and may require multivalent complex formation (Chr3’-Chr1’-Chr1’’-Chr3’). During this evolutionary process, fission behavior evolved, enabling asexual reproduction through fission. This reproductive mode can co-occur with sexual reproduction, as observed in many *Dugesia* species, but eventually a mutation blocks germline development arose, leading to obligate asexuality.

Irrespective of which of the dispersion scenarios applies, our results link the evolution of the asexual strain to structural chromosome rearrangements. It has long been known that the asexual strain carries a reciprocal translocation^36–38^. Our haplotype-phased genome assembly and FISH verifications demonstrate that this translocation is balanced with breakpoints located within two of the large tandem-repeat clusters on the *S. mediterranea* chromosomes (Fig. 2a-b, Fig. 3c). This is a significant finding, as the Chr1/3 translocation has previously been speculated to be causative of the lack of sexual development in the asexual strain. While both strains harbor *piwi^+^/nanos^+^* primordial germ cells (PGCs)^56,69–71^, their development into gonads and accessory reproductive organs is blocked in the asexual strain, resulting in a complete absence of the reproductive system^56^. Our mapping of the Chr1/3 translocation breakpoints to gene deserts (Fig. 3a-c), along with the absence of gene loss or disruption in their vicinity and the lack of reproduction-related genes within 250 kb of the breakpoints (Supporting Information: Gene loss), suggests that neither the Chr1/3 translocation nor the other structural alterations we could characterize are likely to be the primary cause of asexuality. Thus, the ultimate cause of the developmental reproductive system formation arrest is likely caused by changes elsewhere in the genome and thus unrelated to the pervasive chromosomal rearrangements.

Instead, our results suggest that these chromosomal rearrangements emerged in an ancestral sexual population and facilitated the eventual transition to asexuality. The Chr1 inversion that has been previously characterized in the sexual laboratory strain and that we find to be shared by the asexual strain is instructive here^43,52^. Inbreeding experiments show that the Chr1 inversion represses recombination and therefore, we expect the inverted region to be shielded from heterozygosity loss due to inbreeding. We confirm this prediction in the sexual SAR population, which exhibit reduced genetic diversity and frequent runs of homozygosity (ROH) in all regions of the genome not shielded by the Chr1 inversion (Fig. 6e-g), consistent with its high inbreeding coefficient (Fig. 5c). We observed a strikingly similar pattern in the asexual MEN population, where again we see high genetic diversity and minimal ROH specifically in rearranged regions but no differences outside them suggesting they were also shielded from inbreeding (Fig. 5c, Fig. 6g, a reduction of ROH on Chr1 was also previously observed in the CIW4 strain^72^). Crucially, alternative sources of high heterozygosity are insufficient to explain the observed pattern. If the pattern stemmed from the independent accumulation of mutations in both haplotypes—a phenomenon known as the Meselson effect^73,74^, which has been described in asexual Rotifers and found in a few other asexual species^20,26,73,75,76^—we would expect to see relatively uniform divergence along the chromosomes. Similarly, hybridization of two populations with highly divergent chromosome structures, a common origin for many asexual lineages^77^, would also be inconsistent with the rearrangement-dependent pattern we observe. Instead, the pattern indicates the accumulation of heterozygosity in a sexually reproducing ancestor of the asexual MEN population and the asexual laboratory strain.

Based on these observations, we propose the following model for the evolution of asexuality in *S. mediterranea*. The first step was the inversion on Chr1, which suppressed recombination across ∼35% of the genome (Fig. 7c-d). The recombination suppression likely shielded recessive deleterious alleles from inbreeding^78^ and potentially linked locally adapted alleles, thereby protecting them from recombination with maladapted migrant alleles^78,79^ during the evolution of these island populations. The subsequent inversion on Chr2 extended recombination suppression across most of that chromosome, thereby affecting ∼60% of the genome (Fig. 7e). Finally, the Chr1/3 translocation extended recombination suppression to almost the entire genome (Fig. 7f). Although these rearrangements drastically limited recombination, the heterozygosity pattern shaped by inbreeding (see above) and the continued genetic propagation of the rearrangements suggest that sexual reproduction via egg-laying likely persisted through the later stages of this process. Indeed, several sexually reproducing populations of *Dugesia* subtentaculata in the sister genus *carry distinct* reciprocal translocations similar to the Chr1/3 translocation in *S. mediterranea*^26,80,81^, and flatworms are generally known to reproduce sexually despite varying ploidy^61^, mixoploidy^82,83^, and karyotype instability^84–86^. The last stage in the evolution of asexuality in *S. mediterranea* were the mutation(s) that abolished reproductive system development and enabled fissiparous reproduction. While the identification of the causative mutation(s) is an important objective for further studies, the drivers for the transition to behavioral asexuality likely included substantial fertility reductions caused by the preceding inversions and translocations. Specifically, the chromosomal rearrangements gave rise to balanced-lethality systems^87^, as is likely present on Chr1, since homozygous Chr1/Chr1 or Chr1’/Chr1’ embryos in *S. mediterranea* are rarely viable^43^. The Chr1/3 translocation may also have necessitated a modified meiotic chromosome pairing mechanism, such as the multivalent complexes in primroses, which reduce gamete viability^88^. In addition, the block of reproductive system development likely removed important costs of sexual reproduction (e.g., metabolic costs of yolk/gamete formation; behavioral costs of mating), while also providing a dispersal advantage by offering reproductive assurance^89^. Taken together, our model posits that adaptive chromosomal rearrangements progressively undermined the efficiency and genomic benefits of sex, thereby facilitating the ultimate transition to fissiparous asexuality to remove its associated costs.

Our data also provide a new perspective on the interpretation of the large inversion on Chr1 in sexual *S. mediterranea* from Corsica and Sardinia. Previous research described this inversion—with its recombination suppression and haplotype-biased expression of some reproduction-related genes—as a “sex-primed autosome” potentially evolving toward a sex chromosome^52^. While this remains a plausible hypothesis, it is important to note that separate sexes are extremely rare among planarians, possibly occurring in only two of many hundred species worldwide (*Sabussowia dioica* and *Cercyra teisseri*^90^). Our finding that the asexual strain also carries the Chr1 inversion, in addition to independently acquired similar recombination-impairing structural changes, suggests an alternative interpretation: rather than constituting a sex chromosome precursor, the Chr1 inversion may represent an early step in the evolutionary trajectory toward asexuality, as outlined in our model (Fig. 7c–f). This suggests that the genetic architecture often associated with nascent sex chromosomes might, in some systems, represent a predisposition toward asexuality.

The final question remains: How can *S. mediterranea* tolerate the costs of asexuality and reduced recombination, and more broadly, why are asexual reproductive strategies so common among planarians? On the one hand, the lack of strong signatures of reduced selection, even among fast-evolving reproduction-related genes^57,91,92^, could simply reflect the relatively recent origin of asexuality in these lineages. Under this scenario, the asexual *S. mediterranea* strain and other fissiparous planarian strains represent evolutionary dead ends that arise sporadically within parental sexual populations, compete and often outcompete their parental strains due to their reproductive advantages, but ultimately succumb to the cumulative effects of mutation load and other long-term disadvantages inherent to asexuality. On the other hand, the lack of mutational meltdown in our data and the abundance of asexually reproducing strains and species might indicate the existence of particularly effective means of purging deleterious mutations in flatworms. Tentative evidence for gene conversion in *S. mediterranea* supports the possibility that somatic purging mechanisms may operate in planarians^93^; however, such mechanisms would likely require either strong selection or a population bottleneck to be effective^94,95^. The uniquely abundant adult pluripotent stem cells of planarians^29^, with their continuous divisions and consequent potential for clonal dynamics and the dual nature of fissiparous reproduction—as both a mechanism for transmitting somatic mutations to the next generation and a bottleneck that sub-samples the stem cell pool during each fission—create intriguing opportunities for both the purging of deleterious mutations and ongoing evolution, even in obligately fissiparous planarians^24,25,27,96,97^. Therefore, key questions for future research include the extent of intra-animal mosaicism, intra-animal clonal dynamics and bottlenecking effects of fission/regeneration^49^. In addition, population genomics of fissiparous planarian strains and species will be informative for comparisons with pioneering work on clonal evolution in seagrass or corals^94–96^. The genomic resources that we provide lay the foundation for addressing the mechanistic and evolutionary correlates of fissiparous asexuality in planarians.

## Methods

### Biological samples

Asexual *Schmidtea mediterranea* were collected from a stream along Rafal Colom Road, Menorca, Spain (39.90317° N, 4.23302° E) in May 2022. Sexual *S. mediterranea* were collected from the Temo River, Sardinia, Italy (40.398667° N, 8.559492° E) in April 2024. The asexual strain used for genome sequencing, genome assembly, and RNA-seq experiments was the long-term laboratory strain commonly used by most research groups, referred to as CIW4 (internal ID: GOE00071). For shotgun whole-genome sequencing and RNA-seq experiments using sexual worms from the lab, we used the previously established genome strain S2F18^47,48^ (internal ID: GOE00500). S2F18 derives from the S2F2 line originally collected in Sardinia and subsequently inbred through 18 rounds of self-fertilization^43,47^.

### HiFi sequencing

High molecular weight DNA for HiFi sequencing was extracted as previously described^48^. In brief, planarian mucus was stripped using buffered 0.5% (w/v) N-acetyl-L-cysteine (NAC) solution for 10 minutes, followed by 30 minutes of lysis in ice-cold guanidinium thiocyanate buffer (GTC). Lysates underwent phase separation via phenol/chloroform/isoamyl alcohol extraction (25:24:1 ratio), followed by NaCl-induced clarification and isopropanol precipitation. DNA was washed with 70% ethanol and dissolved in TE buffer overnight at 4 °C. Post-extraction, DNA was further purified using a CTAB-based protocol and treated with RNase A. To remove residual contaminants, streptomycin precipitation was performed, followed by PEG/NaCl washing. Finally, DNA was resuspended in pre-dialysis buffer and dialyzed for 4–6 hours using 0.1 μm pore-size membranes. The integrity and yield of the extracted DNA were assessed by pulse field gel electrophoresis (Pippin Pulse, Sage Science) and fluorometric quantification (Qubit, Thermo Fisher).

High-molecular-weight genomic DNA was processed for HiFi sequencing using the low-input SMRTbell Express Template Prep Kit 2.0 (PacBio), as previously described ^48^. Briefly, DNA was sheared to 14–22 kb using the MegaRuptor (Diagenode), and 12–18 μg of sheared DNA was used for library construction. Size selection was performed using BluePippin™. Library preparation and sequencing followed PacBio’s standard protocols using the Sequel® II Binding Kit 2.2 and Sequencing Kit 2.0, with sequencing carried out on SMRT® Cell 8M chips on a Sequel® II system.

### Hi-C sequencing

To obtain the 1 million crosslinked nuclei used for Hi-C library preparation we prepared 2 x 100 animals from our CIW4 laboratory population and 3 x 100 animals from three separate clonal lines we generated from single specimens collected from Menorca (GOE00553 56, GOE00553 9a, GOE00553 9f). Hi-C sequencing was performed as previously described^48^. Briefly, nuclei were isolated using a dounce tissue grinder followed by chromatin conformation capture using the ARIMA-Hi-C High Coverage Kit (Article Nr. A101030-ARI), and Illumina library preparation using the Kapa Hyper Prep kit. The libraries were sequenced using 2 x 150 bp cycles on an Illumina NovaSeq 6000 at the DRESDEN Concept Genome Center, Dresden.

### Hi-C processing

We mapped all Hi-C sequencing reads to haplotype 1 of the sexual (schMedS3) and asexual (schMedA2) assembly using the Arima SV pipeline (v: 1.3, https://github.com/ArimaGenomics/Arima-SV-Pipeline). For each genome, the necessary digested genome files were produced using Juicer (v: 1.6^98^, ‘generate_site_positions.py Arima’) and HiCUP (v: 0.8.0^99^, ‘hicup_digester –arima’). Since automatic detection of structural variations with the state-of-the-art tools relies on the availability of a normalized map of background interactions, which are not available for *S. mediterranea*, we ran the pipeline without the Hi-C Breakfinder module (‘Arima-SV-Pipeline-v1.3.sh -W 1 -B 0 -J 1 -H 0’). Next, we applied the Knight-Ruiz matrix balancing to the interaction map of all datasets (CIW4, S2F18, and Menorca) using HiGlass^100^ and inspected them by locking resolution and zoom level between datasets and manually searching for off-diagonal signals that were exclusive to the sexual or asexual strains.

### Phased *de novo* assembly

PacBio HiFi and Hi-C reads were used to assemble phased contigs with hifiasm v0.7 with default settings (v0.7^101^). Next, Hi-C reads were mapped to the contigs with bwa mem (v0.7.17) with parameter “-5SP -T0”. Reads with mapping quality no less than 10 (-q 10) were further utilized to scaffold the contigs from each haplotype by SALSA (v2^102^) following the hic-pipeline (https://github.com/esrice/hic-pipeline). Scaffolding errors were then manually curated based on the interaction frequency indicated by the intensity of Hi-C signals.

### Identification of genomic strata and breakpoints

We used Minimap2 (v2.28^103^) to align both haplotypes of the S2F18 and CIW4 assemblies in all pairwise combinations (‘-x asm5 -c –eqx --secondary=no’). Then we first identified the genomic strata, defined as sections of the genome with a shared rearrangement history, via the alignment of haplotype 1 and 2 of the CIW4 genome. For this, we parsed the whole-genome alignments using SyRI (v.1.6^104^) run with default parameters and visualized and manually inspected all indicated duplications, translocations, and syntenic alignments. Since we saw that the large parent elements of the alignments explained the observed karyological differences and agreed with the Hi-C signal (see above) we labelled them as genomic strata naming them based on their location on haplotype 1 (which is more similar to the sexual genome) (AXSY where A = Asexual followed by chromosome number S=strata followed by strata number. We named the strata on haplotype 2 based on their homology with haplotype 1 and applying a ‘ to indicate that it is derived (e.g., A1S1 and A1S1’). The same process was used to define strata in the sexual genome. Then we used the other alignments to lift over the strata in the asexual haplotype 2, which represents the rearrangement from one haplotype to both haplotypes of the sexual assembly. Again, care was taken to manually inspect and select the appropriate liftovers to avoid greedy assignment of regions due to short alignments that could represent duplications due to transposable elements or translocations. Breakpoints between strata were double-checked with HiC, dotplots of the alignments, minimap2 alignments generated using R with functions adapted from dotPlotly (https://github.com/tpoorten/dotPlotly), and inspection of LASTZ syntentic chains in our local instance of the UCSC genome browser. Finally, for the Chr1/3 translocation breakpoint we compared similarity of the repetitive region using StainedGlass with a segment size of 2000bp.

### Genome QC

k-mer spectra of short-read or HiFi reads were constructed using meryl (v1.4.1) with k-mer length of 20 and 21, respectively. We assessed the genome phasing and quality using merqury (v1.3^105^) in diploid mode, and performed reference-free quality control and heterozygosity estimation of the HiFi reads using GenomeScope2.0^106^. We assessed genome and transcriptome completeness using BUSCO (v5.3.2^107^) with the metazoan_odb10 reference set, which contains 954 genes. Transcriptome completeness was done using the predicted coding sequences.

### Genome Annotation

We improved upon our previously published gene annotation approach combining Nanopore direct RNA and cDNA long-reads, Illumina cDNA short-reads, and 3P-seq of transcription termination sites (TTS)^108^. For this purpose, we employed long read RNA-sequencing runs from Ivanković et al., 2024 (SRX23002382-SRX23002387), as well as newly sequenced data (See Supporting Table S15). RNA extraction, Oxford Nanopore cDNA library preparation and hybrid genome annotation were all performed as described before^48^. We then extended the strategy to include annotation liftover between haplotypes to avoid inferring haplotype-specific gene loss due to annotation errors. Additionally, we used homology information via alignment of the dd_smed_v6 reference transcriptome to add coding DNA sequence models (CDS) for short proteins and to detect potential chimeric gene models, as we had previously observed a small number of chimeras in the S2F18 annotation. To facilitate reference-based inference of gene loss, we also applied these improvements to our previous annotation of the sexual assembly. For details see Supporting Information: Gene annotation.

### Functional Annotation

To annotate the transcriptome for potential function, we used eggNOG-mapper (v2.1.10^109^) with the options --sensmode ultra-sensitive --report_orthologs -m diamond --dmnd_iterate yes --pfam_ralign realign, and the eggNOG 5.0 database. We also ran InterProScan (v5.54_87.0) with the options -goterms --pathways using all available databases. Additionally, we used DIAMOND with the options --very-sensitive --evalue 1e-5 to search against published protein sequences for Schmidtea mediterranea from NCBI, retrieved via the efetch utility using the query “Schmidtea mediterranea[Organism] AND cds[Feature key]”.

### De novo repeat discovery and annotation

Tandem repeats and transposable elements were annotated as described previously^48^. To identify repetitive sequences that were not annotated using the other approaches, we estimated the genome mappability using genmap (v1.3.0-2,^110^). We used a kmer length of 150 and an error rate of 2. The mappability score is calculated as 1/N, where N represents how often a kmer can be mapped to the genome.

### Chromosome fluorescence in situ hybridization (FISH)

We selected two highly abundant tandem repeats based on their genomic distribution—one of which is located at the Chr1/3 translocation breakpoint—and because *in silico* predictions suggested that the observed structural variation would lead to differences in hybridization signal. Specifically, we selected a 158 bp (158mer) and a 159 bp (159mer) repeat (Supporting Table S1) and designed directly labeled probes for oligo-FISH. The 158mer showed partial similarity to a sequence identified Guo et al. 2022^52^, and both the 158mer and the sequence from Guo et al. match a short probe identified as located close to the centromere of Chr2 by Chretien 2011^111^ (Supporting Information: FISH). We also selected putative centromeric repeats based on their position in the genome assembly, identifying them as the only highly abundant tandem repeats in the expected centromeric region, and designed PCR primers for their amplification. Oligo-FISH using the 158mer and 159mer was performed on both sexual (S2F18) and asexual (CIW4) strains, while standard FISH was conducted only on CIW4. Chromosome spreads were prepared from regenerating tail fragments, and FISH was carried out using established protocols^85^. Details on probe generation, chromosome preparation, and hybridization conditions are provided in Supplementary Information: Chromosome FISH and all primers and oligo DNA probes used in this study are listed in Supporting Table S2.

Chromosome microimages were captured using a CCD camera mounted on an Axioplan 2 compound microscope (Zeiss, Germany) equipped with filter cubes #49, #10, and #15, and controlled via ISIS4 software (METASystems GmbH, Germany) at the Center for Microscopic Analysis of Biological Objects, SB RAS (Novosibirsk, Russia).

### RNA interference experiment

To identify reproduction-related genes, we compared gene expression profiles between sexual (S2F18) and asexual (CIW4) wild-type animals, and between *eGFP* and *ophis* RNAi treatments. Sexual wild-type and RNAi animals underwent a standard regeneration assay. Animals were fed twice with liver paste, which was supplemented with 2 µg/µl of dsRNA in the RNAi treatments. Then heads were amputated anterior to the ovaries but posterior to the eyes, to ensure removal of the reproductive system. After complete regeneration of the head fragments, the animals were fed dsRNA-supplemented liver paste twice a week until control animals had laid cocoons and exhibited visible gonopores. Each treatment was replicated in five 9 cm petri dishes with ten animals each, ensuring balanced incubator distribution and monitoring of food intake. Animals were maintained in Montjuïc water supplemented with 5 μg/mL ciprofloxacin. CIW4 animals were maintained identically, and size-matched individuals were collected. Three animals per group were stained with DAPI to confirm the presence or ablation of the reproductive system.

### Whole-mount DAPI straining

Worms were processed similarly to the standard whole-mount in situ protocol^112^, omitting the staining with riboprobes. Briefly, worms were killed in 7.5% N-Acetyl-L-cysteine in 1X PBS for 5 min with agitation, rinsed with 4% formaldehyde in 0.5X PBSTx0.3, then fixed in fresh fixative for 45 min. After three washes in 1X PBSTx0.3, specimens were incubated in reduction solution at 37°C for 10 min. Following three additional washes, samples were dehydrated through a methanol series and stored at -20°C. For staining, specimens were rehydrated, bleached with 6% H_2_O_2_, PBSTx0.3 for 3h on a light table, and treated with Proteinase K (2 μg/ml) for 10-25 min. After post-fixation, samples were prepared for hybridization, incubated in hybridization buffer at 56°C overnight, and subsequently washed. Specimens were then stained with DAPI (1:1000) overnight at 4°C or for 2.5h at room temperature, washed extensively, and mounted for imaging. Whole-mounts were imaged using an Olympus IX83 microscope equipped with a Yokogawa CSUW1-T2S Spinning Disk system and a Hamamatsu Orca Flash4.0 V3 camera. Images were acquired with an Olympus UPLXAPO 20X air objective (NA = 0.8), with the emission 447/50 filter and excitation at 405 nm.

### Double-stranded RNA production

For RNAi-mediated knock-downs, dsRNA for *ophis* (NCBI accession: KX018822.1, forward primer: ATTGTTAGGATATATTTTGAAACAATTGATG, reverse primer: TGCAGTATTCCATGCATGGC) and *eGFP* was synthesized in vitro and mixed with liver paste as described in Rhouhana et al.^113^. Briefly, linear DNA templates were produced by PCR with T7-AA18 and PR244 primers with pPRT4P-Smed-ophis or pPRT4P-eGFP plasmids as templates. DNA templates were purified and used in in vitro transcription reactions with the T7 RNA Polymerase (Thermo Fisher, EP0111). The dsRNA was then purified by NaCl / PEG-8000 precipitation, diluted to 8 µg/µl, mixed with liver paste (final concentration of 2 µg/µl) and stored at -80 °C until feeding.

### RNA extraction and sequencing

For each treatment, six samples were collected, each consisting of a single worm. Animals were lysed in TRIzol reagent (Invitrogen), and total RNA was extracted using the Direct-zol RNA MiniPrep Plus Kit (Zymo Research, Cat. No. R20700), following the protocol described in ^114^. The libraries were sequenced to an approximate depth of 55 million reads per sample using 2 x 150 bp cycles on an Illumina NovaSeq 6000 at the DRESDEN Concept Genome Center, Dresden.

### Differential expression analysis

Raw reads were screened for contamination with Kraken2 and trimmed using Trimmomatic (v0.39, ^115^) with parameters ILLUMINACLIP:[…]:2:30:10 LEADING:20 TRAILING:20 SLIDINGWINDOW:4:20 MINLEN:35. Trimmed reads were aligned to the sexual reference genome (GenBank accession: GCA_045838265.1) using STAR (v2.7.9a, ^116^) in quantification mode to generate gene-level count matrices. Genes with low raw counts were filtered using the *filterByExpr* function in edgeR (v4.4.0), retaining genes with 5 counts in at least 6 samples and a minimum of 20 counts over all. Library size normalization was performed using the TMM method, and the *voom* transformation was applied to estimate mean-variance trends and generate precision weights. Differential expression analysis was conducted using limma (v3.62.1, ^117^) with a linear model accounting for the three experimental conditions. The following contrasts were tested: *ophis* RNAi vs. asexual wild-type (O_vs_A), sexual control vs. asexual wild-type (S_vs_A), and sexual control vs. *ophis* RNAi (S_vs_O). Genes were considered differentially expressed if they had an absolute log2 fold change ≥ 2 and an adjusted *p*-value < 0.01 (Benjamini-Hochberg correction). Volcano plots were generated using EnhancedVolcano (v1.24.0), with reproduction-related genes highlighted based on curated sexual marker annotations. All downstream analyses were performed in R (v4.4.2) using the tidyverse (v2.0.0) packages.

### Divergence time estimation using four-fold degenerate sites

Our aim was to estimate the divergence time between S2F18 and CIW4 in the absence of fossil or geographic calibration points by approximating neutral divergence using four-fold degenerate sites from single-copy genes. These sites, while more conserved than truly neutral positions, offer the advantage of reliable homologous alignment. To infer orthologous genes, we leveraged the genome assemblies of the three species most closely related to *S. mediterranea*, which we recently published^48^. We used GENESPACE (v1.0.8^118^), which takes advantage of protein similarity via OrthoFinder(v2.5.4^119^) and DIAMOND (v2.0.14^120^) in combination with synteny information inferred using MCSanX (v1.0.0^121^). Then we extracted syntenic single-copy genes using a custom script.

For each single-copy gene, we extracted four-fold degenerate sites using degenotate.py (v1.3^122^) and then concatenated all sites across the single-copy genes into a supermatrix using AMAS^123^. Substitution rates were estimated using IQ-TREE (v2.3.6^124^) on the unpartitioned supermatrix using the GTR+F+I+G model. The cophenetic distance between S2F18 and CIW4 was extracted and halved to obtain the maximum likelihood estimate of substitutions per site since S2F18 and CIW4 diverged. Divergence time (*t*, in generations) was calculated using the equation *t = k / μ*, where *k* is the estimated neutral divergence and *μ* is the mutation rate. We used a pedigree-based estimate of μ = 1×10⁻⁸ mutations per site per generation from *S. mediterranea* as a baseline^52^, and varied this rate by one order of magnitude in either direction to explore the resulting range of divergence time estimates.

### Estimation of LTR insertion age

*Schmidtea* species have abundant and active long terminal repeats (LTR) transposons^47,48^. We estimated the age of LTR insertions restricted to either the sexual or asexual genome to place an upper bound on their divergence. We used SubPhaser (v1.2.6^125^), which identifies LTR insertions and uses phylogenetics to estimate the divergence between them. We ran SubPhaser with default parameters but set the mutation rate to  1×10⁻⁸ (see above).

### non-synonymous/synonymous substitutions and codon usage

Since the available gene annotations for *Schmidtea mediterranea* include genes with low expression and/or repetitive content, we restricted our analyses to genes that passed the minimal expression filter described in the differential expression analysis section. Of the 10,560 single-copy genes inferred using GENESPACE (see above), 10,470 passed this filter and were used for analyses of non-synonymous to synonymous substitution rates (d_N_/d_S_) and codon usage. For each orthogroup, protein sequences were aligned using MAFFT (v 7.480^126^, ‘--globalpair --maxiterate 1000’), and corresponding gap-free codon alignment was generated using PAL2NAL (v14.1^127^, ‘-nogap’). We then used CODEML from the PAML package (v4.9^128^) to estimate all pairwise d_N_/d_S_ ratios (‘runmode = -2, model = 0, fix_kappa = 0, fix_omega = 0’). To assess differences in purifying selection between the sexual and asexual strains, we focused on conserved genes (d_N_/d_S_ < 1) and compared d_N_/d_S_ values between S2F18 and CIW4 relative to other *Schmidtea* species (e.g., S2F18 vs. *S. polychroa* and CIW4 vs. *S. polychroa*). We also evaluated dS between S2F18 and CIW4 to ensure sufficient neutral divergence and applied a filter of dS ≥ 0.1 for a subset of genes.

Codon usage was characterized by estimating the effective number of codons using codonw (v1.4.4^129^, ‘-all_indices’) and the codon deviation coefficient using CAT (v1.3^130^, ‘-b 10000 -c 1’).

Paired, two-sided permutation tests were performed for all parameters to assess differences between S2F18 and CIW4 using the *oneway_test* function from the coin package (v1.4-3) with rank transformation. The test statistic and p-value were calculated based on 1,000,000 resampling iterations. Cohen’s *d* effect size was calculated using the effsize package (v0.8.1), and mean differences with 95% confidence intervals were estimated via bootstrap resampling using the boot package (v1.3-31). In the d_N_/d_S_ analysis, we corrected for multiple testing using the Benjamini-Hochberg procedure, setting the false discovery rate to 0.05. All statistical analyses were conducted in R (v4.4.2) using tidyverse packages (v2.0.0).

### Gene loss analysis

The TOGA^54^ gene annotation pipeline was used to identify genes lost in the asexual schMedA2 assembly. *TOGA was run with default parameters and the improved schMedS3h1 annotation (see above) as reference. We identified genes that were flagged as lost or potentially lost in both haplotypes of* schMedA2*. Additionally, we annotated a gene as conserved if it was present in at least 2/3 of the other Schmidtea assemblies.*

### COI barcoding & short-read sequencing

DNA for barcoding and whole-genome sequencing was extracted as described for HiFi sequencing, but without the post-extraction cleanup steps. To confirm species identity of each sample, we performed PCR amplification followed by Sanger sequencing for an 880 bp fragment of the mitochondrial cytochrome c oxidase subunit I (COI) as described in ^131^ and compared it against the transcriptome of S2F18 and CIW4, and COI haplotype diversity of ^39^. We then removed redundant identical sequences, aligned them using MAFFT (v7.480^126^, ‘--globalpair --maxiterate 1000’), identified HKY + F + I as the best fitting substitution model using ModelFinder^132^, and then inferred a maximum likelihood phylogeny using PHYML (v2.2.4^133^) with 100 non-parametric bootstraps to determine branch support.

Based on these barcoding results, we selected samples for whole-genome sequencing (WGS). For the Menorca sample, DNA from 20 individuals was purified using the Zymo Research Genomic DNA Clean kit. Libraries were prepared with the Illumina Nextera DNA Flex Library Prep Kit and sequenced using 2 × 150 bp cycles on an Illumina NovaSeq 6000 at the Leibniz Institute of Plant Genetics and Crop Plant Research, Gatersleben. For the Sardinia samples, DNA extracts from 29 individuals were prepared with the Illumina Nextera DNA Flex Library Prep Kit and sequenced using 2 × 150 bp cycles on an Illumina NovaSeq 6000 at Genewiz Europe, Leipzig. The three CIW4 and three S2F18 samples were prepared with the KAPA HyperPlus Library Prep Kit and sequenced using 2 × 100 bp cycles on an Illumina NovaSeq 6000 at the DRESDEN Concept Genome Center, Dresden.

### Variant calling

Raw sequencing reads were pre-processed as described for the RNA-seq, aligned to the reference genome using BWA-MEM, and processed following the GATK Best Practices workflow^134,135^. We employed joint genotyping for the πN/πS analysis and per-sample genotyping to assess observed heterozygosity. For joint genotyping, per-sample GVCFs were generated, consolidated by scaffold using GenomicsDBImport, and jointly genotyped with GenotypeGVCFs. For per-sample genotyping, per-sample GVCF files were called using GenotypeGVCFs ‘--all-sites’ option, resulting in detailed calls for each genomic position.

### Heterozygosity

We filtered the per-sample GVCF file using BCFtools by excluding sites with missing genotypes (GT=“mis”), high strand bias (FS > 30), low depth (FMT/DP ≤ 10), or indels (TYPE=“indel”). A custom script based on the R package GenomicRanges was used to calculate the number of variant sites and the total number of callable sites within 100 kb sliding windows, using a 50 kb step size. Windows with fewer than 50,000 callable positions were excluded from further analysis. Observed heterozygosity was calculated as the ratio of variant sites to total callable sites within each retained window.

### πN/πS

Nucleotide diversity parameters (πN and πS) were calculated for both sexual (MEN) and asexual (SAR) populations using SNPGenie (v1.0^136^). Biallelic SNPs found in protein-coding exons were extracted from the joint GVCF using BCFtools (–v snps –T exons.bed –AA –a) and filtered to remove sites with high strand bias (FS > 30) or low depth (FMT/DP ≤ 10). To identify differences in purifying selection, only those genes with πS > 0.001 and πN/πS < 1 were retained.

### PCA and admixture

We used 55 samples for genotype calling, consisting of Menorca (20), Sardinia (29), sexual S2F18 (3), and asexual CIW4 (3). We used ANGSD (v: 0.940 ^137^) to call genotype likelihoods with stringent filtering criteria (-SNP_pval 1e-6 -minMapQ 20 -minQ 20 -uniqueOnly 1 -remove_bads 1 -only_proper_pairs 1 -trim 0 -C 50 -baq 1 -setMinDepthInd 10 -minInd 42) to maintain the high quality of the data. The called genotype likelihoods were then used as input for PCAngsd (v: 1.10 ^138^) to perform Principal Component Analysis (PCA) and NgsAdmix ^139^ for estimating individual admixture proportions. To determine the best K (expected number of genetic clusters or ancestral populations) for admixture proportions, 10 replicates per K were run in NgsAdmix, and log-likelihood values were extracted. Best K was identified by Evanno’s method ^140^ implemented in CLUMPAK ^141^.

### Inbreeding coefficients

The BAM alignments of Menorca and Sardinia were used separately to call genotype likelihoods using ANGSD with additional flags (-doMajorMinor 1 -doMaf 1 -skipTriallelic 1 -doGlf 3) to generate Beagle files for both populations. ngsF-HMM^142^, a tool to estimate inbreeding coefficients (F_IS_) and IBD tracts efficiently from low coverage sequencing data, was used to calculate individual genome-based F_IS_ for both populations.

### Linkage disequilibrium

The BAM alignments of Menorca and Sardinia were used separately to call genotype likelihoods using ANGSD, similar to what was mentioned above, except they were only restricted to the sites within coding regions. The number of sites called was then prioritized for estimating pairwise LD values using ngsLD ^143^ with additional flags—-probs to specify input type and—-max_kb_dist 1000 to specify the distance required to consider the sites for LD calculations. The results were then subsetted randomly to a smaller portion to plot the exponential LD curves for both populations.

### SMC++

The BCFtools version 1.19 ^144^ was used to call variants using all Menorca (20) and Sardinia (29) individuals with filtering (-C50 -q30 -Q30) to retain better quality called variants in the resultant vcf file. SMC++ ^145^ vcf2smc module was used to convert the vcf data into smc++ input data. SMC++ estimate was used to calculate effective population size (Ne) estimates for both populations separately using a mutation rate of 1e^-08^ per generation, followed by an SMC++ plot to visualize the temporal population size trends for each population. SMC++ split module was used using both populations together to calculate the estimate of joint demography, which gives an indication of the split time between the two populations.

### Genetic diversity and differentiation

The site frequency spectrum (SFS) was estimated for Menorca and Sardinia populations by obtaining genotype likelihoods using ANGSD. To maintain high-quality variants we used uniquely mapped and properly mate-mapped reads with mapping quality adjustment (-C 50), adjusting q-scores around indels (-baq 1), minimum mapping quality of 20 (-minMapQ 20), minimum base quality of 20 (-minQ 20), at least 75% of individuals considered (-minInd Nx0.75), depth of at least 10 bases at each site in each individual (-setMinDepthInd 10). The obtained genotype likelihoods in the form of sample allele frequency (SAF) were then used to estimate SFS using realSFS module of ANGSD. The theta estimations were obtained using saf2theta module of ANGSD using SAF. SFS and theta estimations were further used to get estimates of diversity over 50 KB non-overlapping windows using thetastat module of ANGSD. The mean of Watterson’s estimator (θ_w_), Pairwise nucleotide diversity (θ_π_) and Tajima’s D of the 50 KB windows normalized by the number of sites in each window for each population was calculated for all chromosomes.

## Data Access

All sequencing data generated for this study and all code used in this analysis will be made available together with the peer-reviewed version of this article. To request pre-publication access contact the corresponding author.

## Acknowledgements

We thank Rick Kluiver, Jens Krull, Claudia Koch, and the MPI-NAT animal caretakers for worm care; Delia Niehaus, Fruzsina Ficze, and Til Schubert for DNA extractions; Fruzsina Ficze also for Hi-C preparations and COI barcoding; the IIT Genomics Facility (Diego Vozzi, Yeraldin Castillo Spelorzi, and Edoardo Henzen) for Nanopore sequencing support; Sylke Winkler and Nils Stein for supervising sequencing experiments; Tom Brown and Martin Pippel for assistance with genome assembly; Andrei Rozanski, Ferenc Kagan, and Elham Bavafaye Haghighi for bioinformatics support; Miquel Vila-Farré and Thomas Brochier for field collection support; and Leonard Drees for dsRNA production. We thank Leushkin Evgeny and Michael Hiller for help with the TOGA analysis. We also thank the Leibniz Institute of Plant Genetics and Crop Plant Research and the Dresden Concept Facilities for supporting sequencing experiments. Field collections in Menorca were conducted under GOIB permit 37077. JNB was supported by Swiss National Science Foundation (SNFS) Grant P500PB_206673. JCR received funding from the European Research Council (ERC) under the European Union’s Horizon 2020 research and innovation program (grant agreement number 649024), from the Max Planck Society and from the German Research Foundation (DFG) graduate school GRK 2984: Evolutionary Genomics: Consequences of Biodiverse Reproductive Systems (EvoReSt) and personal project RI 2449/51. LP was supported by intramural funding of the Istituto Italiano di Tecnologia. KSZ was supported by Ministry of Education and Science of Russian Federation (projects FWNR-2022-0015 and FSUS-2024-0018) and the results included in this manuscript were obtained before 02/24/22.

## Author contributions

JNB and JCR conceptualized the study, JNB designed and performed all experiments and analyses not otherwise specified. JNB and JCR wrote the manuscript. ABP performed genotype-likelihood-based population genomic analyses. LP conducted nanopore-based genome annotation. KZ and NR performed FISH experiments. LR performed repeat analysis. MZ and AM performed genome assembly. All authors read and approved the final manuscript.

