## Supporting Information for "Stepwise emergence of recombination suppression precedes fissiparous asexuality in the *planarian Schmidtea mediterranea*"

#### Genomic rearrangements trace the stepwise evolution of asexuality in the planarian *S. mediterranea*

### 1 Genome quality control

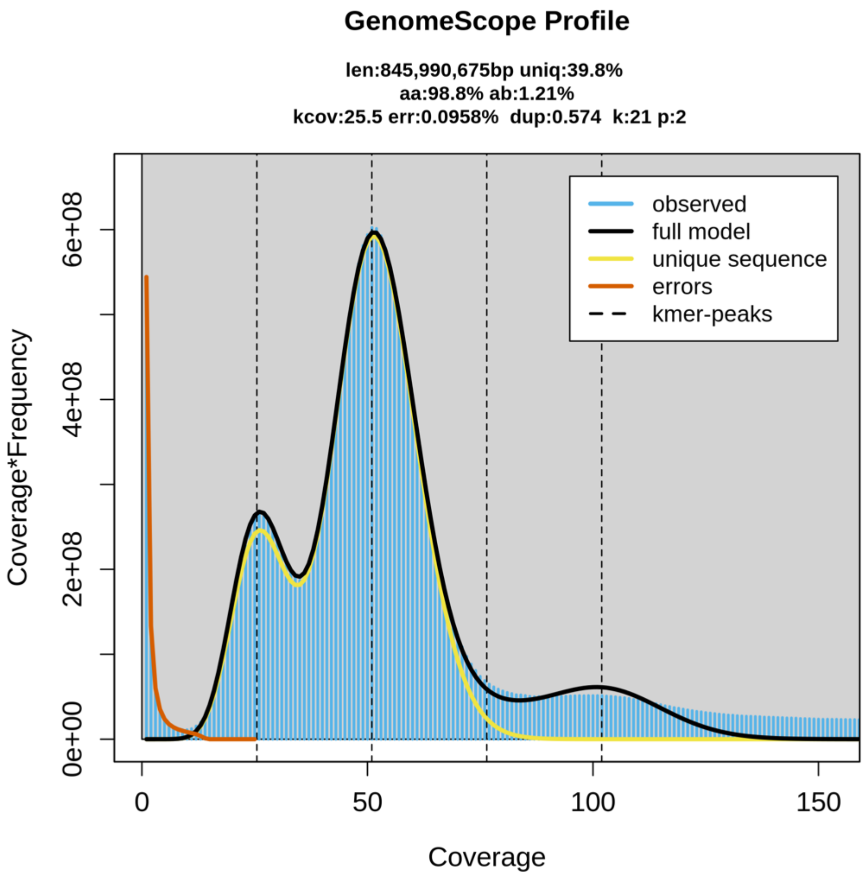

Figure 1 K-mer distribution in the asexual HiFi reads and the inferred genome size, and heterozygosity based on the model of GenomeScope 2.0.

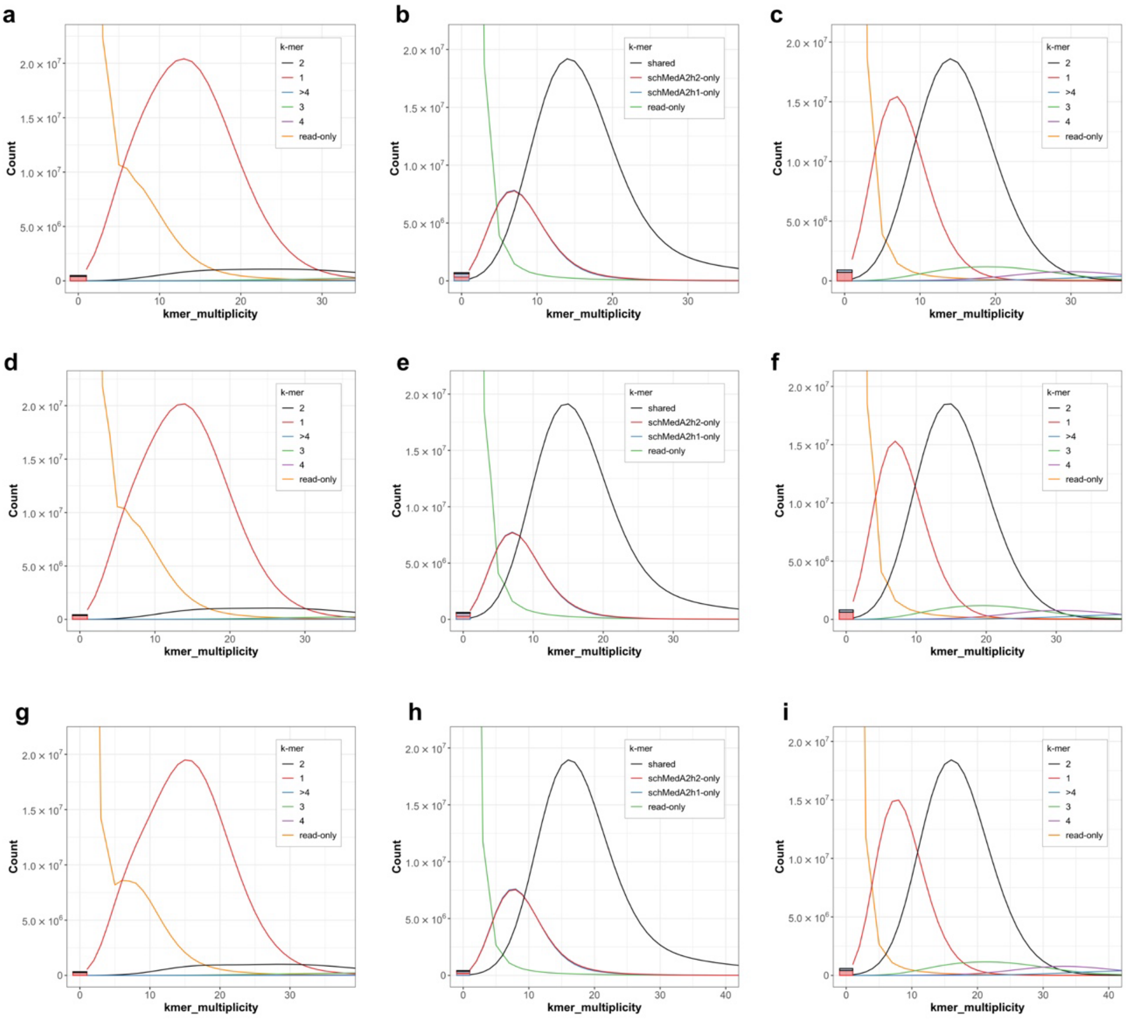

Figure 2 Merqury analysis of the schMedA2 assembly using three independent short read datasets generated from the genome strain. Rows are results from the three datasets (a-c:, d-f:,g-i:,**XXX Accessions will be added before publication XXX**). First column shows k-mer spectra for haplotype 1 clearly indicating a substantial proportion of k-mers with a high coverage occur only in the reads. Second column shows, that when considering both haplotypes these reads are divided between the assemblies for each haplotype, thus indicating good phasing efficiency. Finally, the last column shows the k-mer spectrum of the diploid assembly. We can see that the first peak (shown in red), which represents the haplotype specific k-mers is of a similar height than the second peak, which represents the shared diploid k-mers. Therefore, heterozygosity is high.

### 2 Gene annotation

For liftover, transcripts were aligned against the genome using minimap2 (v2.28-r1209[1],-cx splice:hq --cs -G 100k -L -u b -N 1), and CDS were predicted for both the existing annotation and the alignments. CDS were predicted using TransDecoder (v5.7.1) configured to detect short proteins (-m 40) and retain low-scoring models with homology to PFAM or UniRef90 databases as assessed using MMseqs2 (v16.747c6[2]). Next we collected extensive statistics on the liftovers using an all-vs-all sequence search of all inferred CDS using MMseqs2, assessed exon structure comparison using GFFcompare (v0.12.2[3]), assessed transcript coverage using minimap2, and collected statistics on splice junction coverage, expression, and overlap with Start-seq and transcription termination signals using SQANTI3 (v5.2.1[4], --CAGE_peak start peaks, --polyA_peak TTS, --coverage STAR file, --skipORF).

Transcripts were filtered using stringent quality criteria, with high-confidence annotations requiring at least 90% transcript identity and matching exon structure. CDS models with >80% identity and >80% coverage of the query were assessed as matches. When the primary CDS failed to meet thresholds, alternative CDS predictions were evaluated. Overlapping CDS were simply annotated, while non-overlapping CDS constituted potential chimeras. These were split if separated by a termination signal; otherwise, they were flagged as potentially chimeric. For liftovers without reference matches, novel coding genes were created if their predicted CDS passed filtering criteria. For mono-exon transcripts, we required a TSS ratio >5, start location within a START-Seq peak, and polyA site within 10bp of the TTS. Novel non-coding transcripts required a TSS ratio >5 and polyA motif within 10bp of TTS, with mono-exons additionally requiring either start within a START-seq peak or end within a TTS site. Following liftover completion, transcripts were re-clustered into final gene models.

### 3 Karyology literature

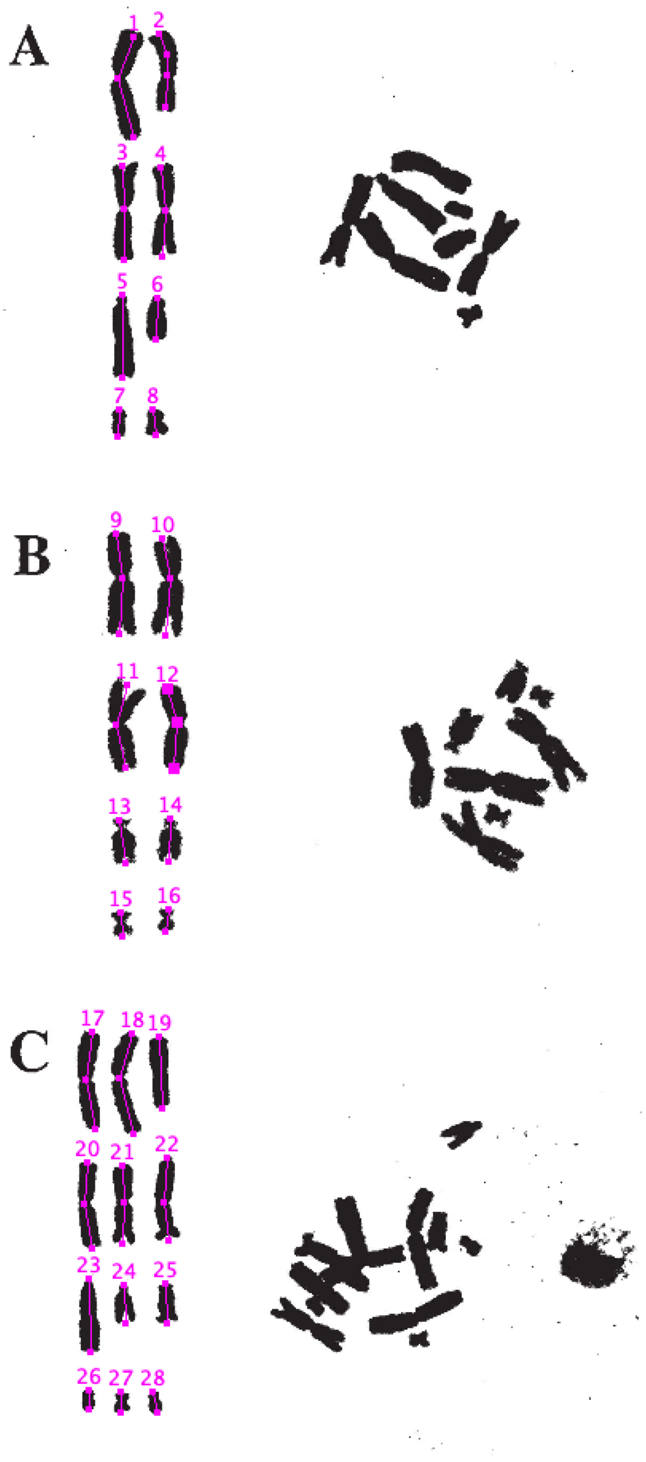

Fig 1 - Mitotic metaphase plates and karyotypes of *Schmidtea mediterranea*. A) 2n = 8, asexual diploid; B) 2n = 8, sexual diploid; C) 3n = 12, asexual triploid.

Figure 3 Estimation of relative chromosome size based on the publication of Baguñà et al. 1999.

Table 1 Relative size estimates of S. mediterranea chromosomes from the literature. Values for Baguñà et al. 1999 were extracted as shown in Figure 3.

| population | chr | chr_len (relative) | publication |
|---|---|---|---|
| asexual 2n | Chr1 | 197.7 | Baguñà et al. 1999 |
| asexual 2n | Chr1' | 140.7 | Baguñà et al. 1999 |
| asexual 2n | Chr2 | 176.0 | Baguñà et al. 1999 |
| asexual 2n | Chr2' | 168.8 | Baguñà et al. 1999 |
| asexual 2n | Chr3' | 156.0 | Baguñà et al. 1999 |
| asexual 2n | Chr3 | 78.1 | Baguñà et al. 1999 |
| asexual 2n | Chr4 | 52.0 | Baguñà et al. 1999 |
| asexual 2n | Chr4' | 50.4 | Baguñà et al. 1999 |
| sexual 2n | Chr1 | 190.8 | Baguñà et al. 1999 |
| sexual 2n | Chr1' | 186.1 | Baguñà et al. 1999 |
| sexual 2n | Chr2 | 160.2 | Baguñà et al. 1999 |
| sexual 2n | Chr2' | 152.2 | Baguñà et al. 1999 |
| sexual 2n | Chr3 | 82.6 | Baguñà et al. 1999 |
| sexual 2n | Chr3' | 82.0 | Baguñà et al. 1999 |
| sexual 2n | Chr4 | 46.0 | Baguñà et al. 1999 |
| sexual 2n | Chr4' | 44.4 | Baguñà et al. 1999 |
| Menorca 3n | Chr1 | 185.9 | Baguñà et al. 1999 |
| Menorca 3n | Chr1' | 194.6 | Baguñà et al. 1999 |
| Menorca 3n | Chr1'' | 134.8 | Baguñà et al. 1999 |
| Menorca 3n | Chr2 | 163.5 | Baguñà et al. 1999 |
| Menorca 3n | Chr2' | 148.7 | Baguñà et al. 1999 |
| Menorca 3n | Chr2'' | 155.5 | Baguñà et al. 1999 |
| Menorca 3n | Chr3 | 136.0 | Baguñà et al. 1999 |
| Menorca 3n | Chr3' | 74.1 | Baguñà et al. 1999 |
| Menorca 3n | Chr3'' | 74.1 | Baguñà et al. 1999 |
| Menorca 3n | Chr4 | 38.0 | Baguñà et al. 1999 |
| Menorca 3n | Chr4' | 38.5 | Baguñà et al. 1999 |
| Menorca 3n | Chr4'' | 40.1 | Baguñà et al. 1999 |
| Barcelona 2n | Chr1 | 41.95 | Ribas 1990 |
| Barcelona 2n | Chr1' | 29.38 | Ribas 1990 |
| Barcelona 2n | Chr2 | 33.85 | Ribas 1990 |
| Barcelona 2n | Chr2' | 33.85 | Ribas 1990 |
| Barcelona 2n | Chr3 | 27.86 | Ribas 1990 |
| Barcelona 2n | Chr3' | 15.41 | Ribas 1990 |
| Barcelona 2n | Chr4 | 8.82 | Ribas 1990 |
| Barcelona 2n | Chr4' | 8.82 | Ribas 1990 |
| Menorca 3n | Chr1 | 42.19 | Ribas 1990 |
| Menorca 3n | Chr1' | 30.49 | Ribas 1990 |
| Menorca 3n | Chr2 | 33.37 | Ribas 1990 |
| Menorca 3n | Chr2' | 33.37 | Ribas 1990 |
| Menorca 3n | Chr3 | 27.78 | Ribas 1990 |
| Menorca 3n | Chr3' | 17.78 | Ribas 1990 |

| | | | |
|---|---|---|---|
| Menorca 3n | Chr4 | 8.71 | Ribas 1990 |
| Menorca 3n | Chr4' | 8.71 | Ribas 1990 |
| Mallorca 2n | Chr1 | 37.74 | De Vries 1984 |
| Mallorca 2n | Chr2 | 31.77 | De Vries 1984 |
| Mallorca 2n | Chr3 | 29.5 | De Vries 1984 |
| Mallorca 2n | Chr3' | 16.18 | De Vries 1984 |
| Mallorca 2n | Chr4 | 7.6 | De Vries 1984 |
| Barcelona 2n | Chr1 | 34.96 | De Vries 1984 |
| Barcelona 2n | Chr2 | 31.76 | De Vries 1984 |
| Barcelona 2n | Chr3 | 28.31 | De Vries 1984 |
| Barcelona 2n | Chr3' | 18.08 | De Vries 1984 |
| Barcelona 2n | Chr4 | 10.55 | De Vries 1984 |

Table 2 Size of chromosome scaffolds, the scaffold names, and the assigned chromosome to allow comparisons to data in Table 1.

| population | chr | scaffold | scaffold length (bp) | publication |
|---|---|---|---|---|
| Barcelona CIW4 | Chr1 | Chr1_h1 | 327,432,606 | This study |
| Barcelona CIW4 | Chr1' | Chr3_h2 | 224,520,763 | This study |
| Barcelona CIW4 | Chr2 | Chr2_h1 | 272,298,851 | This study |
| Barcelona CIW4 | Chr2' | Chr2_h2 | 267,951,672 | This study |
| Barcelona CIW4 | Chr3 | Chr3_h1 | 119,429,788 | This study |
| Barcelona CIW4 | Chr3' | Chr1_h2 | 219,263,528 | This study |
| Barcelona CIW4 | Chr4 | Chr4_h1 | 46,130,935 | This study |
| Barcelona CIW4 | Chr4' | Chr4_h2 | 46,645,998 | This study |

Table 3 Relative differences (∆) between Chr1 and Chr3 homologs in the asexual strain, based on estimates from the literature and the genome assembly presented in this study. Our estimate agrees well with the literature.

| population | publication | ∆ Chr1/Chr1' | ∆ Chr3/Chr3' (%) |
|---|---|---|---|
| Barcelona 2n | Ribas 1990 | 30 | 44.7 |
| Barcelona 2n | Baguñà et al. 1999 | 28.8 | 49.9 |
| Barcelona 2n | De Vries 1984 | - | 36.1 |
| **Barcelona 2n** | **This study** | **31.4** | **45.5** |
| Mallorca 2n | De Vries 1984 | - | 45.2 |
| Menorca 3n | Ribas 1990 | 27.7 | 36 |
| Menorca 3n | Baguñà et al. 1999 | 30.7 | 45.5 |

### 4 Hi-C analyses

#### 4.1 Hi-C quality control

Table 4 Mapping statistics of Hi-C data from the sexual lab strain (S2F18), the asexual lab strain (CIW4), three isolines of the Menorca population (MEN 9a, MEN 9f, MEN 5+6), and all Menorca isolines combined (MEN all). Mapping was performed with the sexual (schMedS3) and asexual genome (schMedA2) as a reference.

| schMedS3 haplotype 1 | S2F18 | CIW4 | MEN all | MEN 9a | MEN 9f | MEN 5+6 |
|---|---|---|---|---|---|---|
| Input (M reads) | 1138.4 | 584.7 | 482.6 | 153.5 | 144.7 | 184.4 |
| Mean insert size (bp) | 308 | 345 | 401 | 359 | 443 | 403 |
| SE mapped (%) | 92 | 89.2 | 79.2 | 74.4 | 79.8 | 82.6 |
| SE truncated (%) | 29.6 | 36.4 | 20.7 | 23.9 | 16.7 | 21 |
| Mapped (M reads) | 222.4 | 49 | 38 | 10.1 | 11.9 | 15.9 |
| Duplicate (M reads) | 33.5 | 9.8 | 8.5 | 2.2 | 2.8 | 3.5 |
| Unique (M reads) | 159.7 | 56.1 | 24.9 | 7.2 | 7 | 10.8 |
| Cis >15kb (M reads) | 89.7 | 32.3 | 14 | 3.9 | 4.1 | 6.1 |
| Trans (M reads) | 51.1 | 16.9 | 7.5 | 2.6 | 1.8 | 3.1 |
| Restriction site (M reads) | 42.2 | 3.3 | 7.4 | 2 | 2.3 | 3 |
| Restriction site (%) | 12.8 | 4.2 | 12.1 | 11.9 | 12.3 | 12.1 |
| Cis/Trans | 1.8 | 1.9 | 1.9 | 1.5 | 2.3 | 1.9 |

| schMedA2 haplotype 1 | S2F18 | CIW4 | MEN all | MEN 9a | MEN 9f | MEN 5+6 |
|---|---|---|---|---|---|---|
| Input (M reads) | 1138.4 | 584.7 | 482.6 | 153.5 | 144.7 | 184.4 |
| Mean insert size (bp) | 310 | 345 | 397 | 352 | 440 | 401 |
| SE mapped (%) | 88.4 | 92 | 83.5 | 78.7 | 84.3 | 86.7 |
| SE truncated (%) | 29.6 | 36.4 | 20.7 | 23.9 | 16.7 | 21 |
| Mapped (M reads) | 112.1 | 82.3 | 43.8 | 11.2 | 13.1 | 19.4 |
| Duplicate (M reads) | 15.1 | 17.2 | 10.1 | 2.5 | 3.1 | 4.5 |
| Unique (M reads) | 71.2 | 98.9 | 29.9 | 8.1 | 7.7 | 14.1 |
| Cis >15kb (M reads) | 37 | 57.9 | 17 | 4.3 | 4.5 | 8.1 |
| Trans (M reads) | 24.6 | 29.5 | 8.9 | 3 | 1.9 | 4.1 |
| Restriction site (M reads) | 24.6 | 4.6 | 8.3 | 2.2 | 2.6 | 3.5 |
| Restriction site (%) | 14.8 | 3.5 | 11.7 | 11.9 | 12.3 | 11.2 |
| Cis/Trans | 1.5 | 2 | 1.9 | 1.5 | 2.3 | 2 |

#### 4.2 Hi-C comparison S2F18 vs CIW4

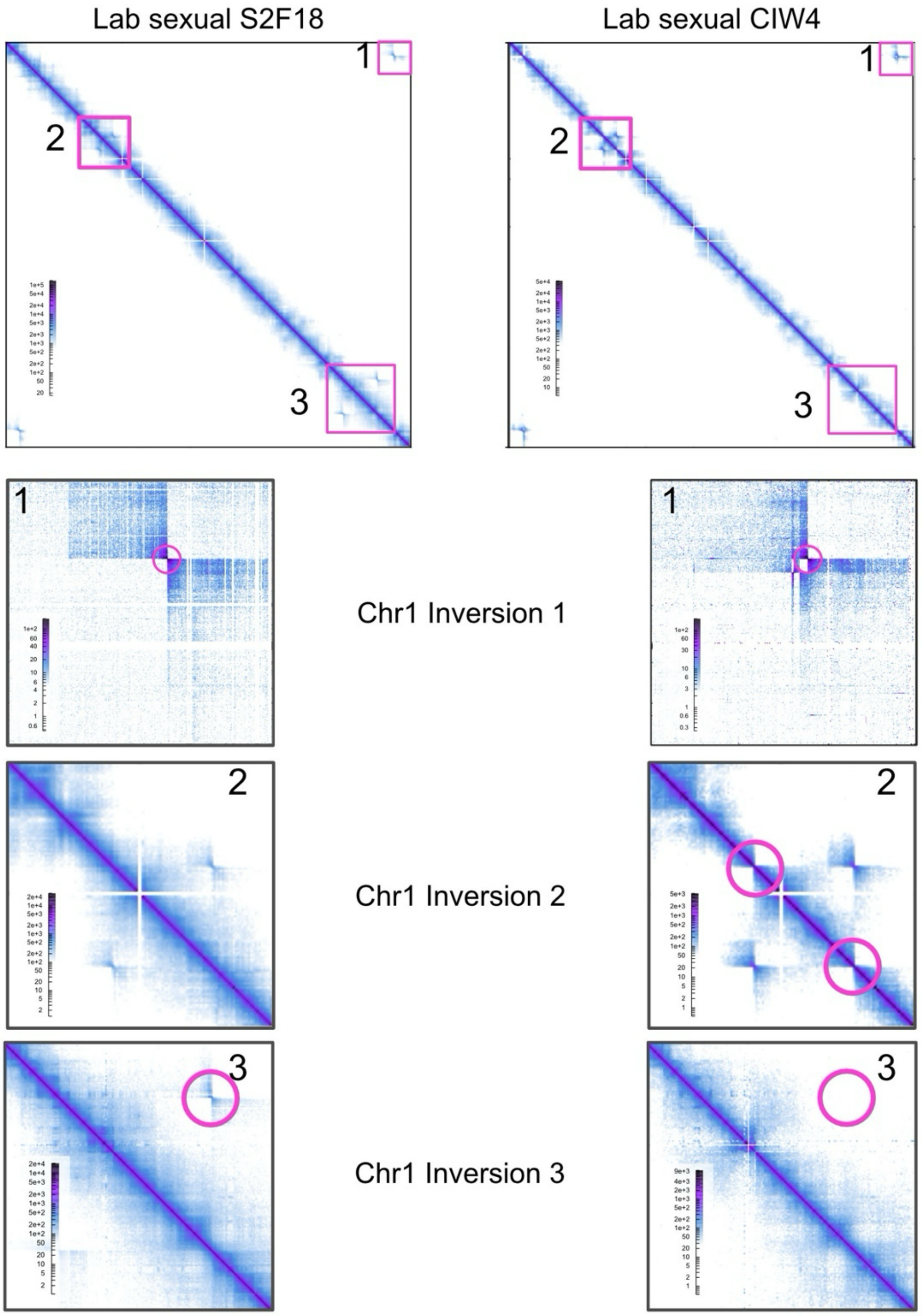

Figure 4 Hi-C signal of the sexual laboratory strain (S2F18) and the asexual laboratory strain (CIW4) when mapped onto Chr1 of the sexual reference genome (schMedS3h1). Pink boxes highlight regions of interest that are enlarge below. The enlarged regions of interest, showing Hi-C signal in the sexual strain S2F18 (left) and asexual strain CIW4 (right). The signal indicates that the asexual strain shares the large inversion (1) but does not have the two smaller inversions (2,3). Note, that asexual haplotype 1 is inverted in relation to the sexual reference chromosome leading to the off-diagonal signal but a lack of signal on the diagonal in (2). Further note that the signal in 1 is duplicated in CIW4 due to a duplication of that region in the asexual strain.

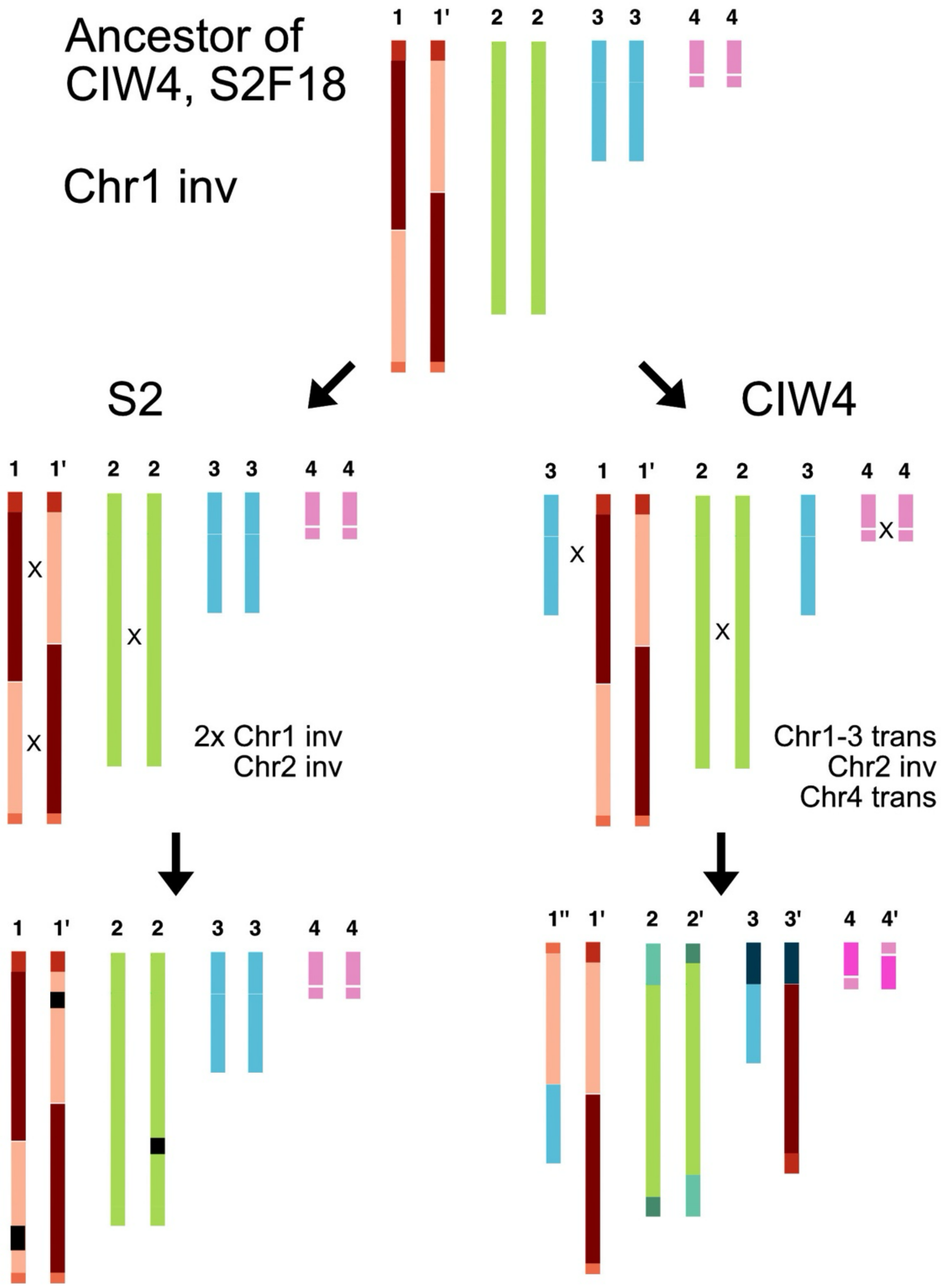

Figure 5 Most parsimonious history of chromosomal rearrangements explaining the observed structural rearrangements between CIW4 and S2F18. The Chr1 inversion is shared, while all other rearrangements have occurred independently.

#### 4.3 Hi-C Chromosome 4

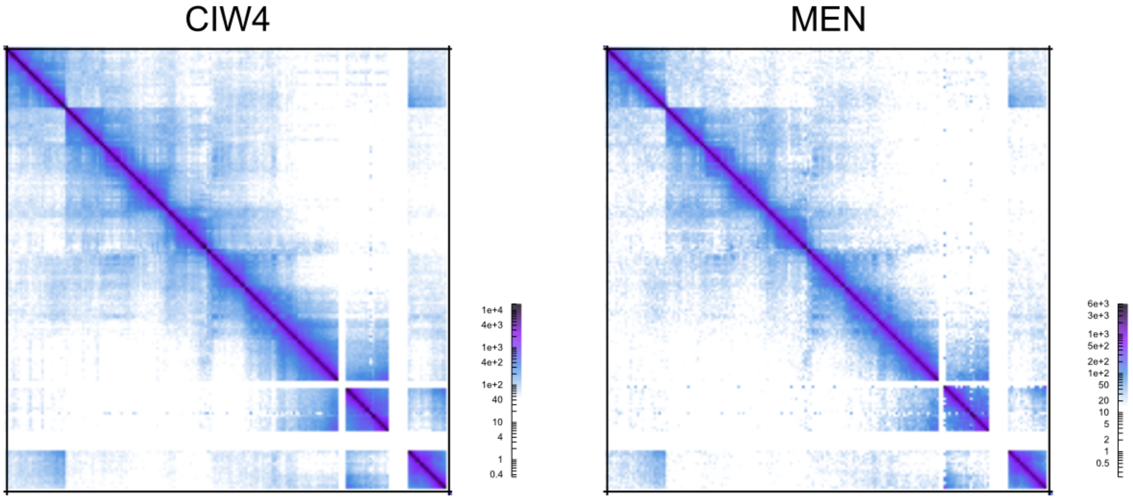

Figure 6 Hi-C contact map of Chr4 of the schMedA2h1 assembly, showing the Hi-C signal from the asexual laboratory strain (CIW4) and from a combination of three clonal lines generated from wild asexual individual from Menorca (MEN). Note the highly repetitive region towards the end of the scaffold, indicated by large stretches of no contact and the intense off-diagonal signal between the start and the end. Resolution of the contact map is 250k bp per pixel.

#### 4.4 Hi-C Menorca isolines

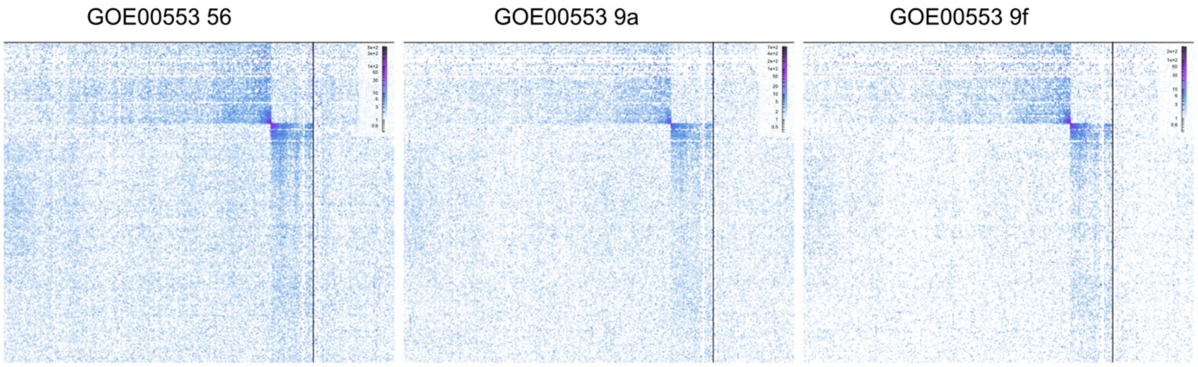

Figure 7 Hi-C signal of three clonal lines derived from wild asexuals from Menorca. Shown is the signal for the Chr1 inversion 1. Resolution of the contact map is 250k bp per pixel.

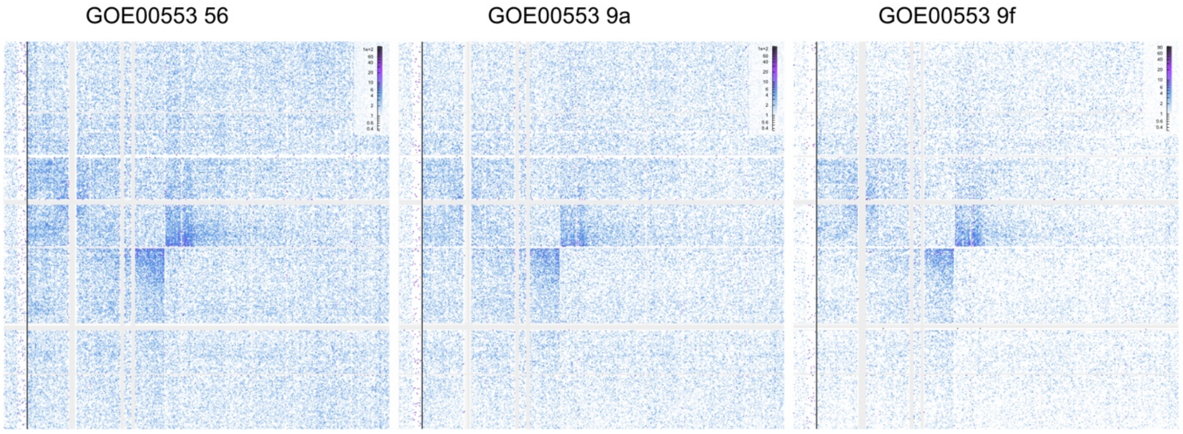

Figure 8 Hi-C signal of three clonal lines derived from wild asexuals from Menorca. Shown is the signal for the Chr1/3 translocation. Resolution of the contact map is 250k bp per pixel.

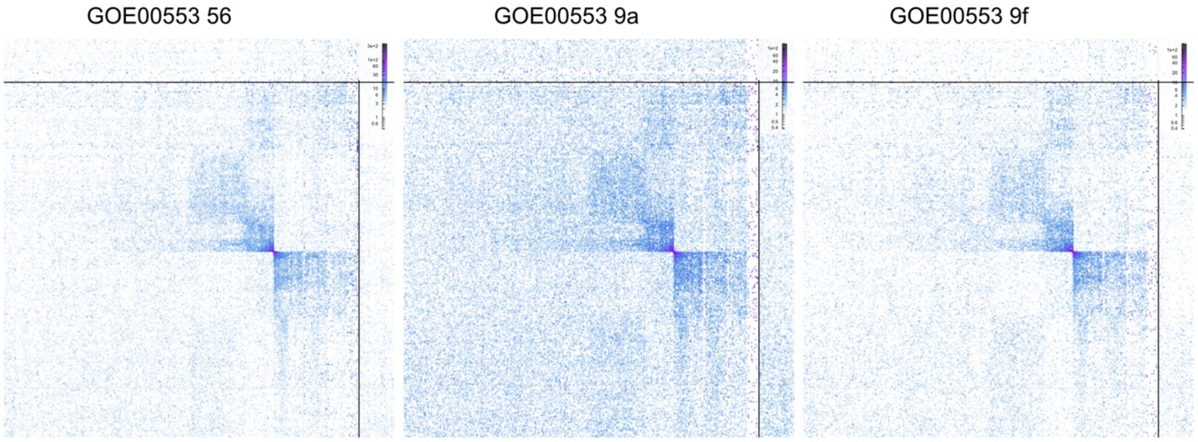

Figure 9 Hi-C signal of three clonal lines derived from wild asexuals from Menorca. Shown is the signal for the Chr2 inversion. Resolution of the contact map is 250k bp per pixel.

### 5 Chromosome FISH

#### 5.1 Generation of DNA probes specific to the predicted centromeric satellite DNA repeats

Total gDNA was isolated from a pool of 10 worms. Before DNA extraction, worms were washed in 0.05% NAC solution (N-acetyl-L-cysteine) for 1 min. Worms were disrupted by homogenization, and DNA was extracted with the phenol-chloroform procedure followed by ethanol precipitation. The extracted gDNA was used as a template in the following PCR.

Amplification of fragments of centromeric DNA repeats was performed using standard PCR with specific forward and reverse primers (Table X). Then PCR products were labeled in an additional 20 cycles of PCR using corresponding specific primers in the presence of Flu-12-dUTP, fluorescein-5(6)-carboxamidocaproyl-[5(3-aminoallyl)2′-deoxyuridine-5′-Triphosphate)] (Biosan, Novosibirsk, Russia), or TAMRA-5-dUTP, 5-tetramethylrhodamine-dUTP (Biosan, Novosibirsk, Russia). Modified oligonucleotides (CEN-SmedSat1_158 and CEN-SmedSat2_159) were ordered in the laboratory of synthetic biology (ICBFM SB RAS, Novosibirsk, Russia). All primers and oligo DNA probes used in this study are listed in Table S2.

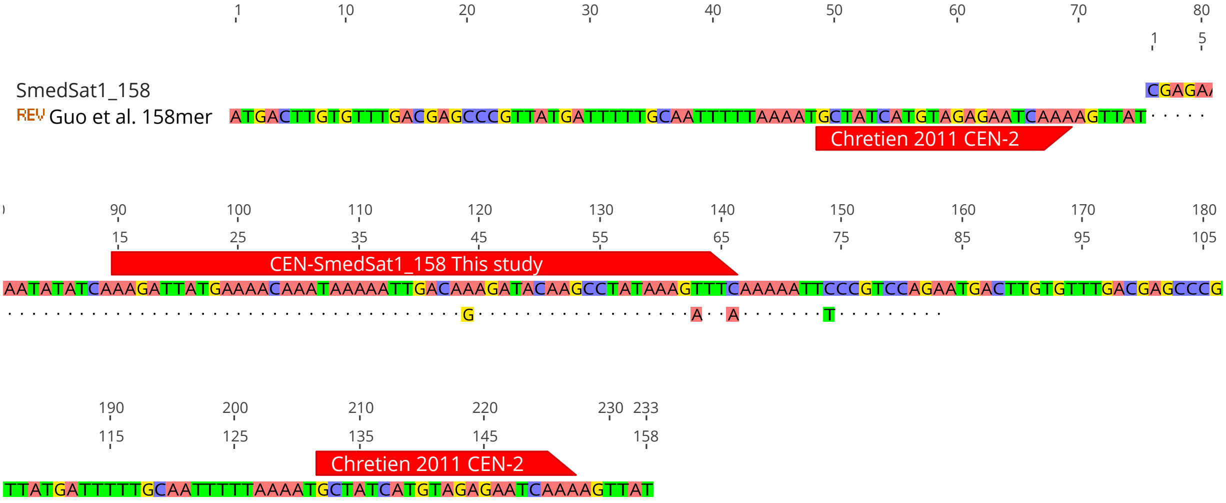

Figure 10 Similarity of the SmedSat 158mer repeat targeted in this study to the k-mer studied by Guo et al., showing the predicted probe binding location of the probe used in this study and the one used by Chretien 2011.

#### 5.2 Chromosome slide preparation

Metaphase spreads were obtained from dividing cells from regenerated tissue. Worms of *S. mediterranea* from both sexual and asexual strains were cut and left to regenerate in 6-well plates (1 individual per well) containing 1XPW for two days. Then the regenerated tissue of the worm was isolated and used for the following slide preparation. For that, the pieces of tissue were placed into 0.2% colchicine (in 1XPW) for 1 h at RT. Immediately after, they were transferred into 0.2% KCl (in 1XPW) for 1 h at RT. After that, the treated material was carefully placed onto the clean, dry slides, and residues of the solution were removed. Immediately after, a small amount (about 20 µL) of the fixative solution I (glacial acetic acid:ethanol:bidist H2O, 3:3:4) was added, and the material was macerated in the fixative with a glass needle. Then the slide with the macerated material was placed into water bath vapors, and the fixatives II and III (glacial acetic acid:ethanol, 1:1, and glacial acetic acid, correspondingly) were placed onto the material and left for 2-3 min. Then the prepared chromosome slides were air-dried at RT and used for the following procedures.

#### 5.3 Fluorescent *in situ* hybridization

FISH experiments with DNA probes generated using PCR were carried out as was described earlier (Zadesenets et al., 2020). Briefly, after a pretreatment step including RNA-seq, pepsin, and fixation, the DNA of chromosomal material and generated DNA probes was denatured simultaneously at 72°C for 2 min. For hybridization, chromosome slides were incubated overnight at 37°C. After slides were washed using 50% formamide in 2×SSC (three times for 5 min), 2×SSC (two times for 5 min), and 0.2×SSC (two times for 5 min). All washings were done at 45°C.

Oligo-FISH was performed on metaphase chromosomes of *S. mediterranea* (sexual and asexual strains). Pre-treatment of chromosome slides was done according to the standard FISH technique (Zadesenets et al., 2020). Oligo-FISH probes (CEN-SmedSat1_158, 22 mkM, and CEN-SmedSat2_159, 67 mkM) were dissolved in hybridization mixture (50% formamide, 10% dextran sulfate, 4×SSC) until the final concentration (1:500 and 1:1000, correspondingly). After that, the simultaneous hybridization of DNA of chromosome slides and oligo-FISH probes was done at 72°C for 2 min and followed by hybridization for two days at 37°C in a dark place. Then the slides were washed with 2× SSC, 0.5× SSC, and 1× TNT (100 mM Tris-HCl, 150 mM

NaCl, 0.1% Tween 20, pH 7.4) for 15 min each at RT. After chromosome slides were dehydrated in 70%, 90%, and 100% graded ethanol for 2 min each at RT and air-dried. Metaphase chromosomes were counterstained with 4',6-diamidino-2-phenylindole solution (DAPI) (VectaShield, USA) according to standard protocol.

#### 5.4 Quantification of FISH signal in CIW4 and S2F18

Table 5 Signal quantification of oligo-FISH signal of the 158mer and 159mer based on 50 metaphase plates from different specimens of the asexual strain (CIW4). When it was cytologically possible to separate the homologs, they were labeled accordingly. Otherwise, they were grouped by signal intensity.

| Oligo | Chr. 1 | | Chr. 2 | | Chr. 3 | | Chr. 4 | |
|---|---|---|---|---|---|---|---|---|
| | Chr1 | Chr1' | hom. 1 | hom. 2 | Chr3 | Chr3' | hom. 1 | hom. 2 |
| 158mer | 2.84±0.618 | 2.43±0.5 | 2 | 2 | 1.02±0.143 | 1.65±0.48 | 0 | 0 |
| 159mer | 0 | 0 | 0 | 0 | 1 | 1.65±0.52 | 2 | 1 |

Table 6 Signal quantification of oligo-FISH signal of the 158mer and 159mer based on metaphase plates from different specimens of the sexual strain (S2F18). Homologous chromosomes were grouped by signal intensity.

| Oligo | Chr. 1 | | Chr. 2 | | Chr. 3 | | Chr. 4 | |
|---|---|---|---|---|---|---|---|---|
| | hom. 1 | hom. 2 | hom. 1 | hom. 2 | hom. 1 | hom. 2 | hom. 1 | hom. 2 |
| 158mer | 2.60±0.599 | 2.53±0.50 | 2.23±0.42 | 2.17±0.38 | 1.75±0.43 | 1.55±0.5 | 0 | 0 |
| 159mer | 0 | 0 | 0 | 0 | 1 | 1 | 2 | 2 |

#### 5.5 FISH images

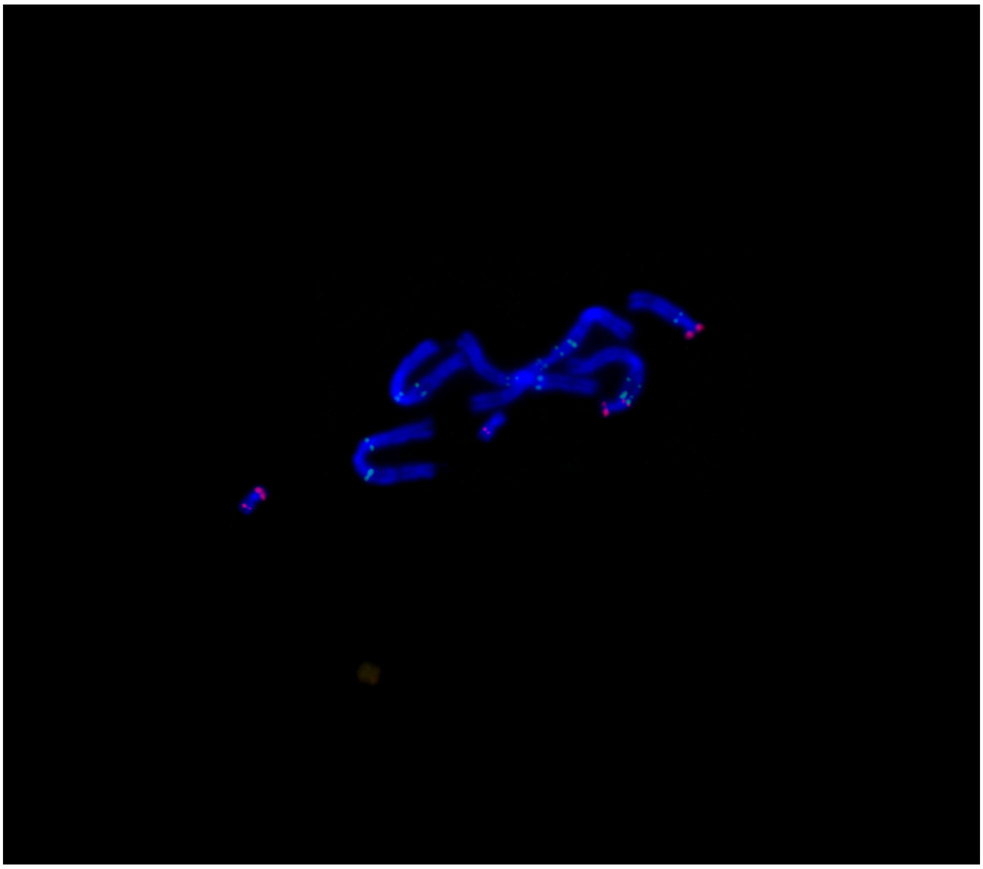

Figure 11 Raw metaphase spread used for visualization in Figure 1d of the main text. Note that overlapping chromosomes had to be digitally separated.

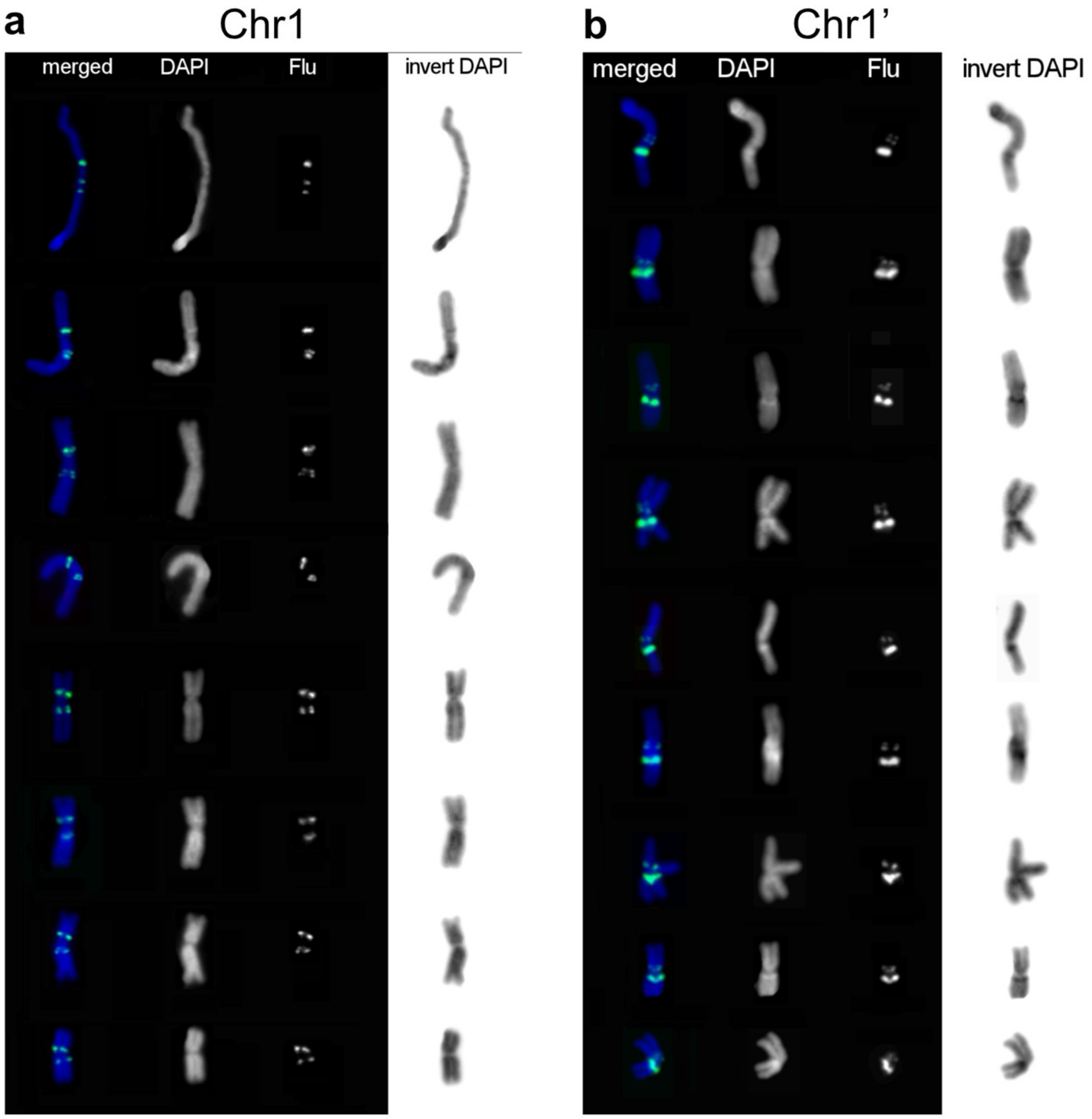

Figure 12 Example variation in chromosome FISH of Chr1 (a) and the rearranged Chr1’ (b) in the asexual strain (CIW4). Flu signal shows labelling of the 158mer.

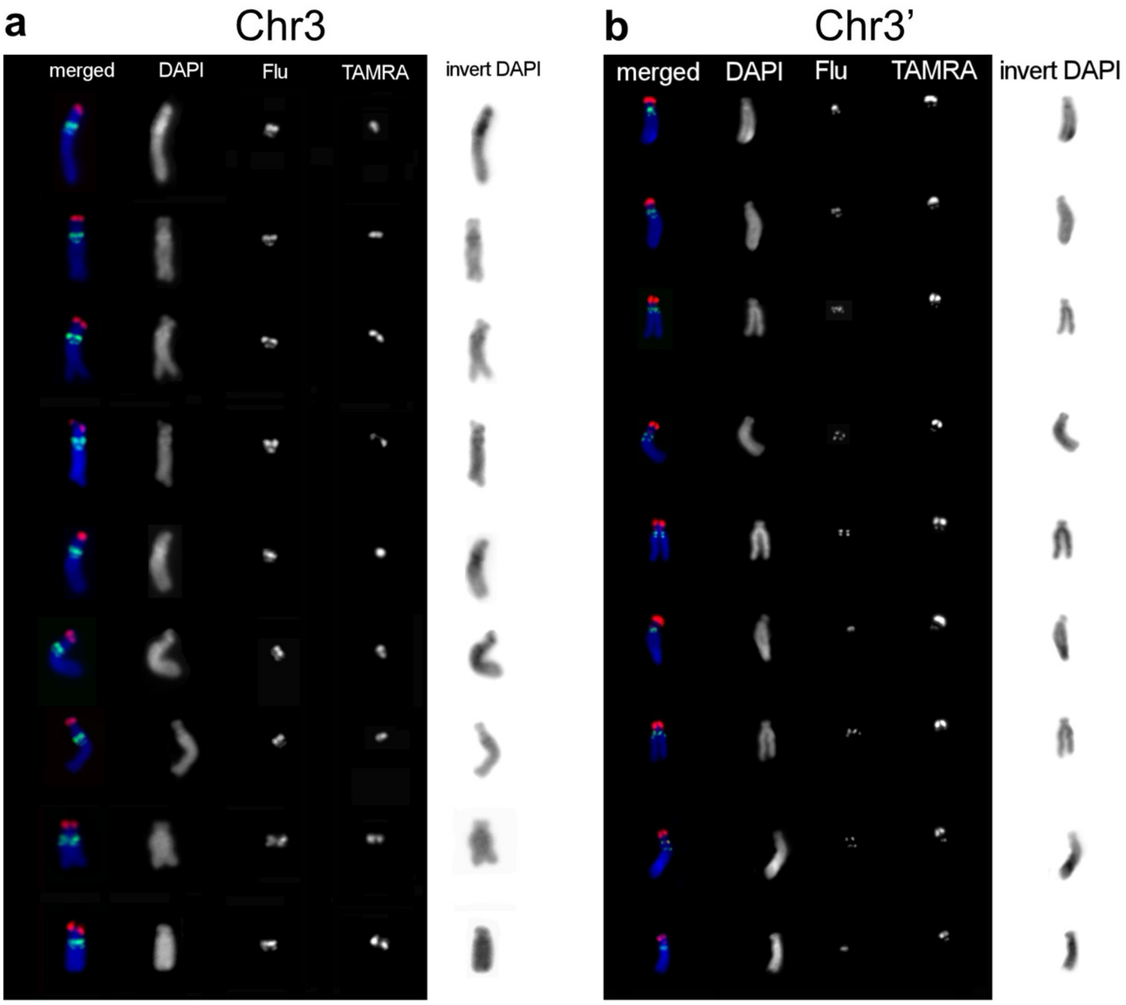

Figure 13 Example variation in chromosome FISH of Chr3 (a) and the rearranged Chr3’ (b) in the asexual strain (CIW4). Flu signal shows labelling of the 158mer and TAMRA shows labelling of 159mer.

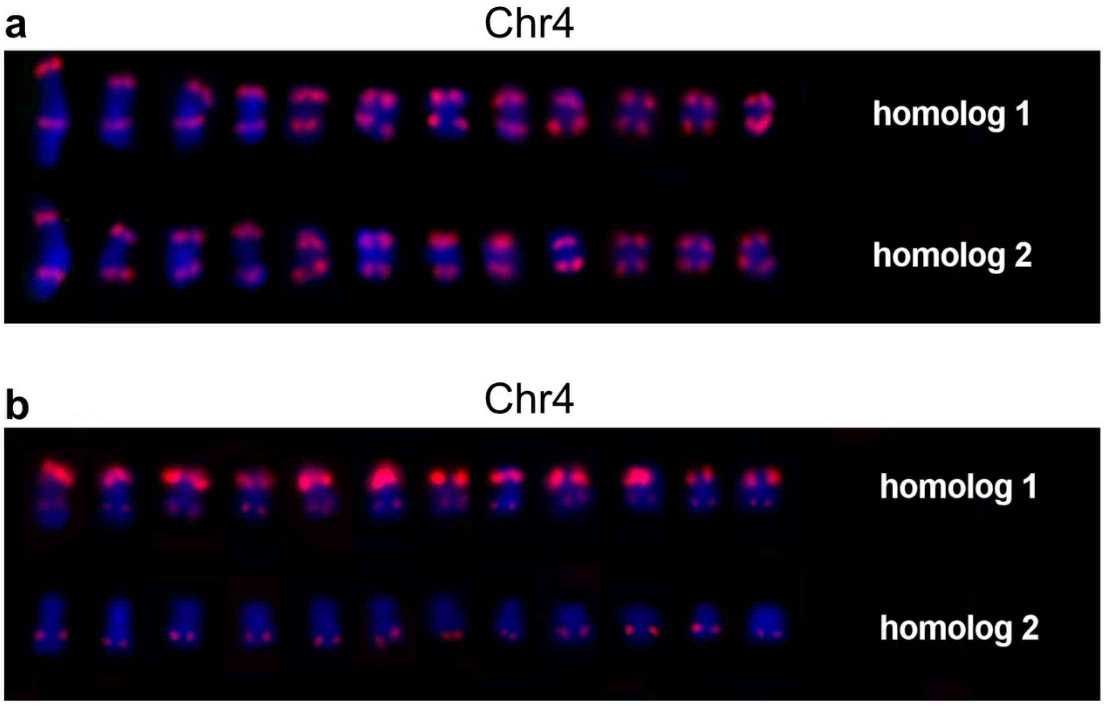

Figure 14 Example variation in chromosome FISH of Ch4 and Chr4’ in the sexual strain (S2F18, a) and Chr1 and asexual strain (CIW4, b). Signal shows labelling of the 159mer.

### 6 Strata liftover

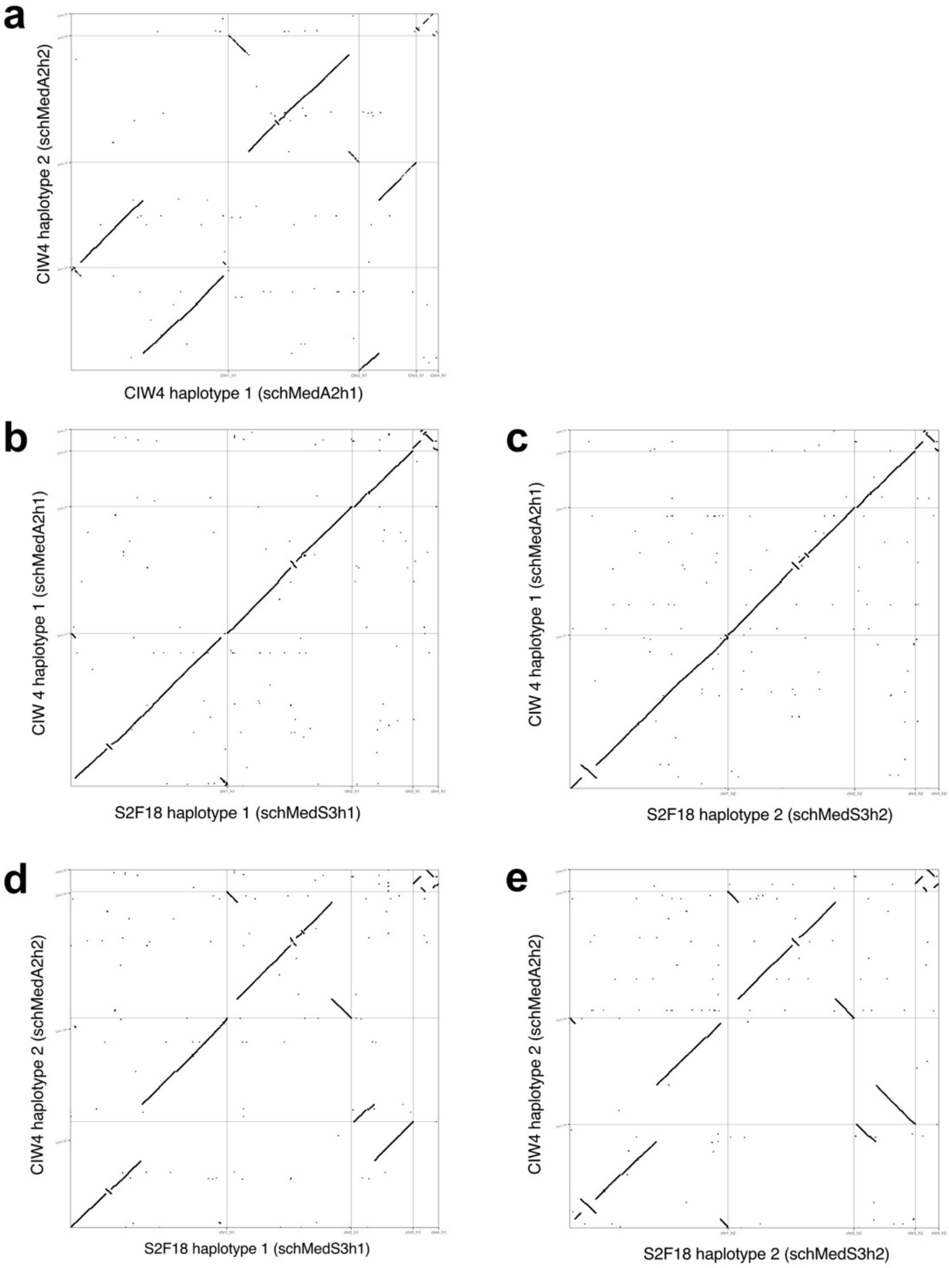

Figure 15 Dotplots representing whole genome alignments between (a) asexual (CIW4) haplotype 1 and CIW4 haplotype 2; (b) sexual (S2F18) haplotype 1 and CIW4 haplotype 1; (c) S2F18 haplotype 2 and CIW4 haplotype 1; (d) S2F18 haplotype 1 and CIW4 haplotype 2; (e) S2F18 haplotype 2 and CIW4 haplotype 2. Dots represent minimap2 alignments with a minimum length of 1 kb and a mapping quality >30.

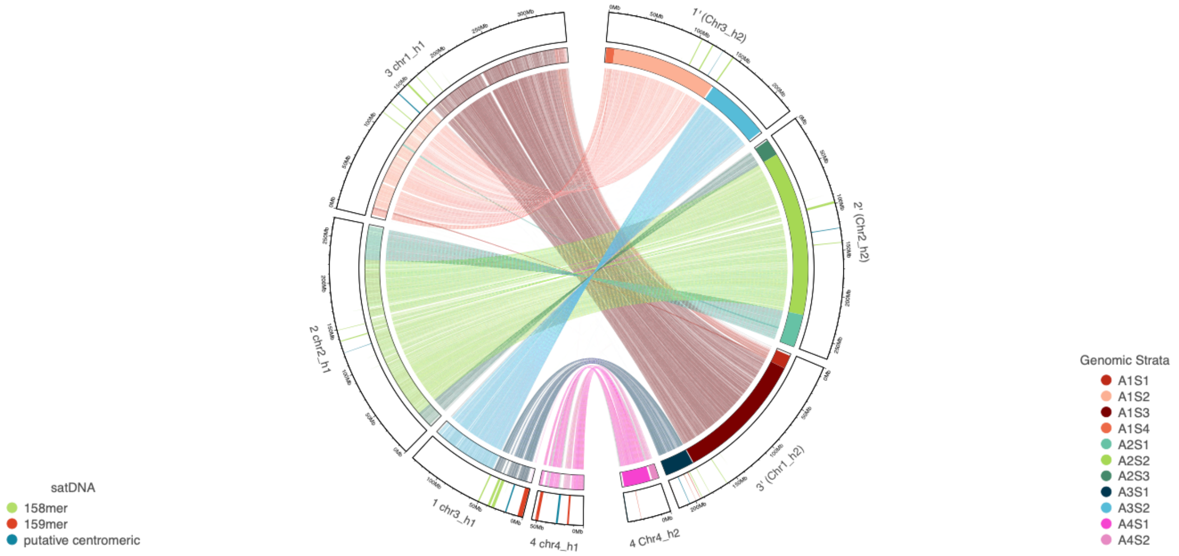

Figure 16 Whole-genome alignment based liftover of CIW4 haplotype 2 (schMedA2h2) genomics strata, shown on the right, onto S2F18 haplotype 1 (schMedS3h1) shown on the left.

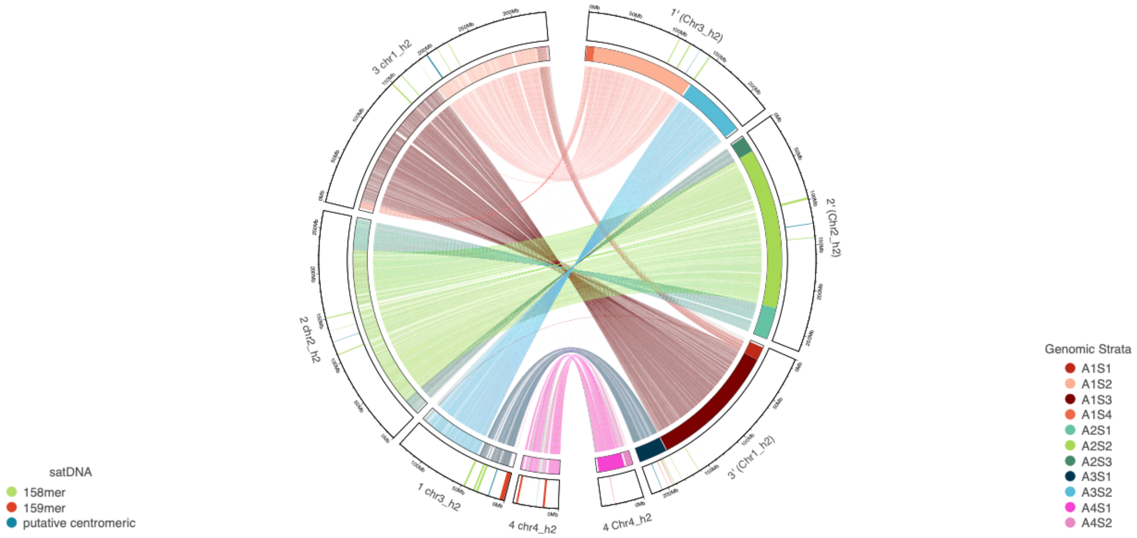

Figure 17 Whole-genome alignment based liftover of CIW4 haplotype 2 (schMedA2h2) genomics strata, shown on the right, onto S2F18 haplotype 2 (schMedS3h2) shown on the left.

### 7 Reproduction-related genes

To gain functional information about gene loss candidate genes, we established a detailed annotation of reproduction-related genes. We generated gene expression profiles from sexual and asexual wild-type animals, as well as from sexual control *RNAi* (eGFP) and *ophis* RNAi, which is known to block the development of all reproductive system components at an early stage[5,6]. Using these data, we annotated 2,136 protein-coding genes as high-confidence reproduction-related if they were significantly down-regulated in *ophis* (RNAi) and asexual wild-type compared to sexual control (see Methods, Main text Fig. 3d-f, Supporting Tables S4-6). In this section, we will detail the experimental design and the results of this analysis.

#### 7.1 Sample size

To determine the appropriate sample size and read depth we performed a power calculation using ssizeRNA single function from the R package ssizeRNA. We oriented our projected number of differentially expressed genes based on the comparison of sexual and asexual wild-type data from Davies et al., [7] where, with an average read count of 20 million per library, 52% of transcripts were significantly differentially expressed and 60% had a log2fold-change of >2. This determined that increasing the coverage to 40 million reads per library and a sample size of six would result in an acceptable beta of 0.72, assuming an alpha of 0.05.

#### 7.2 Confirmation of RNAi via phenotyping

We confirmed the phenotype of sexual animals by observing a gonopore and laid cocoons in the eGFP RNAi and sexual wild-type group. While in the ophis RNAi treatment only two animals had a slightly light spot in the area where a gonopore might appear. Size was assessed six days before collection and revealed no significant size difference between RNAi groups, but slightly larger size in wild-type, likely because they received *ad-libitum* food (Figure 18). We also confirmed the loss of major reproductive structures in *ophis* RNAi via whole-mount DAPI staining of animals in the treatment groups (Figure 19).

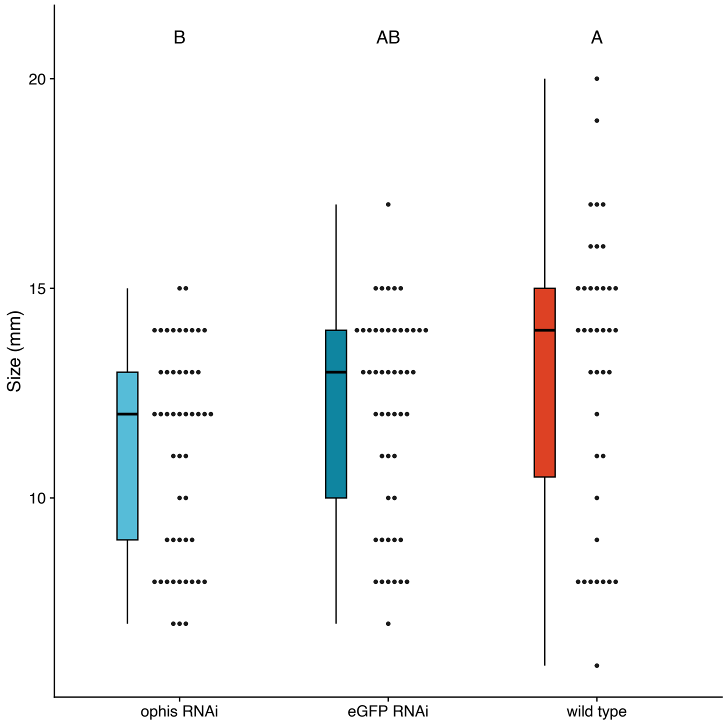

Figure 18 Body length of the three sexual treatment groups six days before collection. Boxes indicate the interquartile range (IQR), with the median as a horizontal line. Whiskers extend to 1.5×IQR, and individual points represent raw data. Different letters indicate statistically significant differences between groups (Tukey HSD, $p < 0.05$).

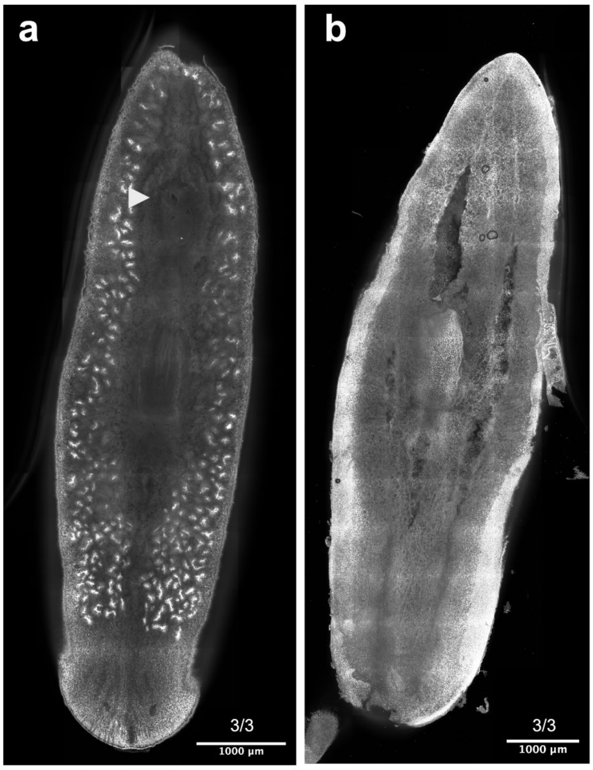

Figure 19 DAPI staining of representative samples from the (a) *eGFP* RNAi and (b) *ophis* RNAi treatments. Three specimens were imaged from each condition. (a) shows a single slice of the image stack with a view of the abundant testes. (b) is a maximum intensity projection of the entire stack showing the lack of testes. White arrow in (a) highlights the copulatory apparatus, which is absent in (b).

#### 7.3 Differential expression analysis

We observed that, as expected, the *eGFP* RNAi control and the sexual wild-type clustered in the PCA (

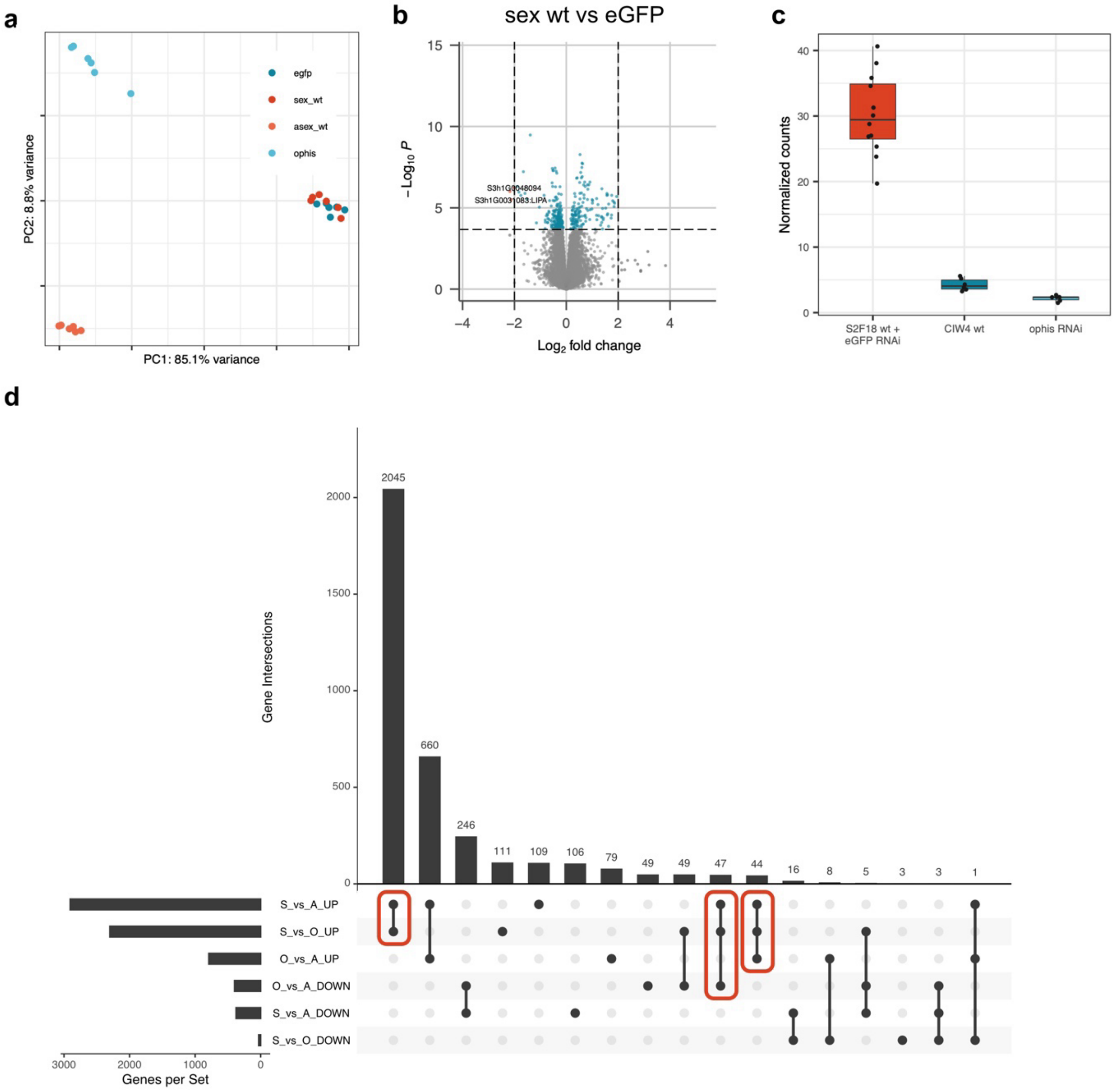

Figure 20a). Only three genes were detected as significantly down-regulated and six as significantly up-regulated in the eGFP RNAi vs sexual wild-type contrast (

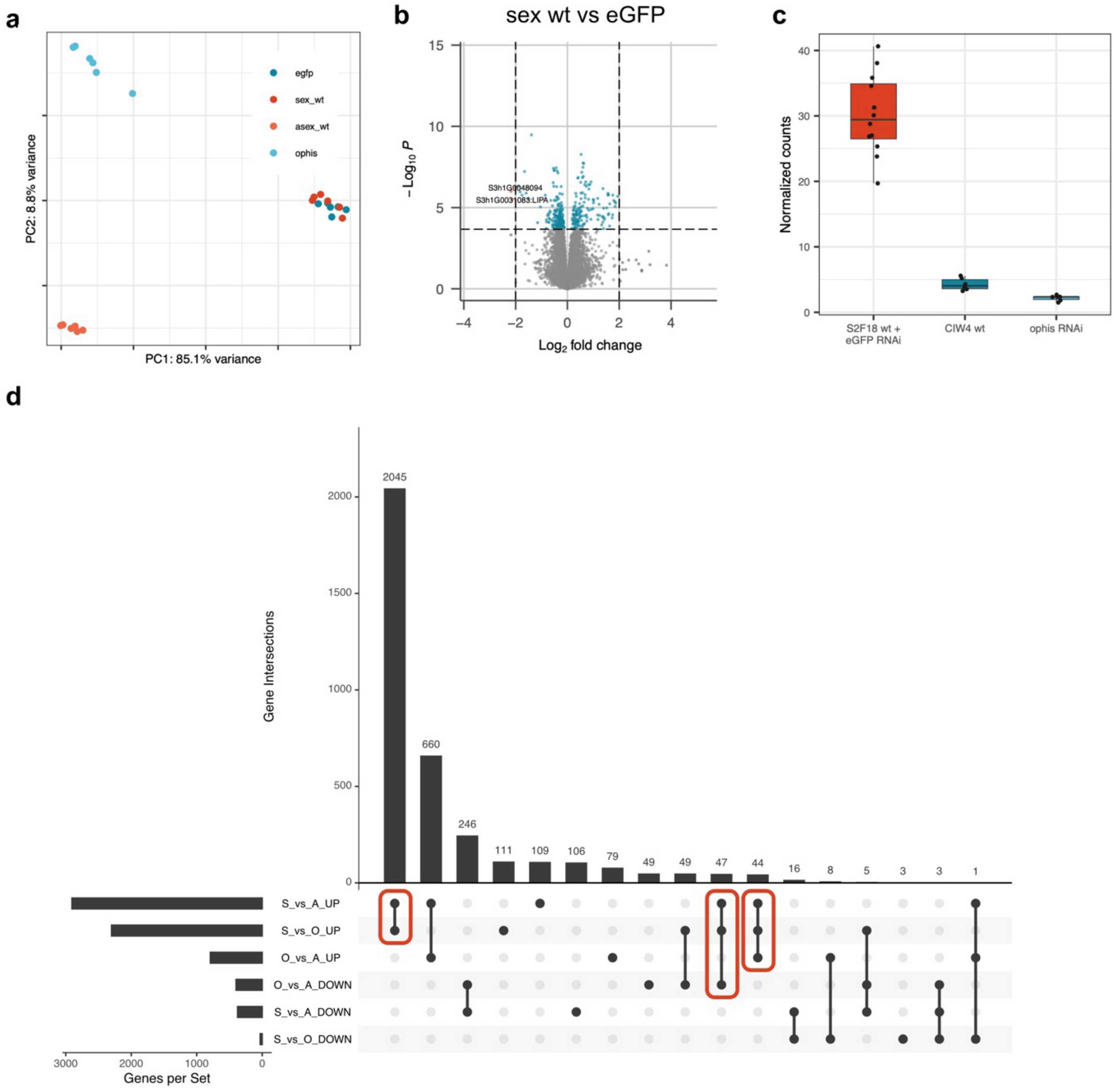

Figure 20b). Therefore, we grouped these two controls for the entire analysis. Next, we confirmed that *ophis* RNAi indeed knocked down the gene (

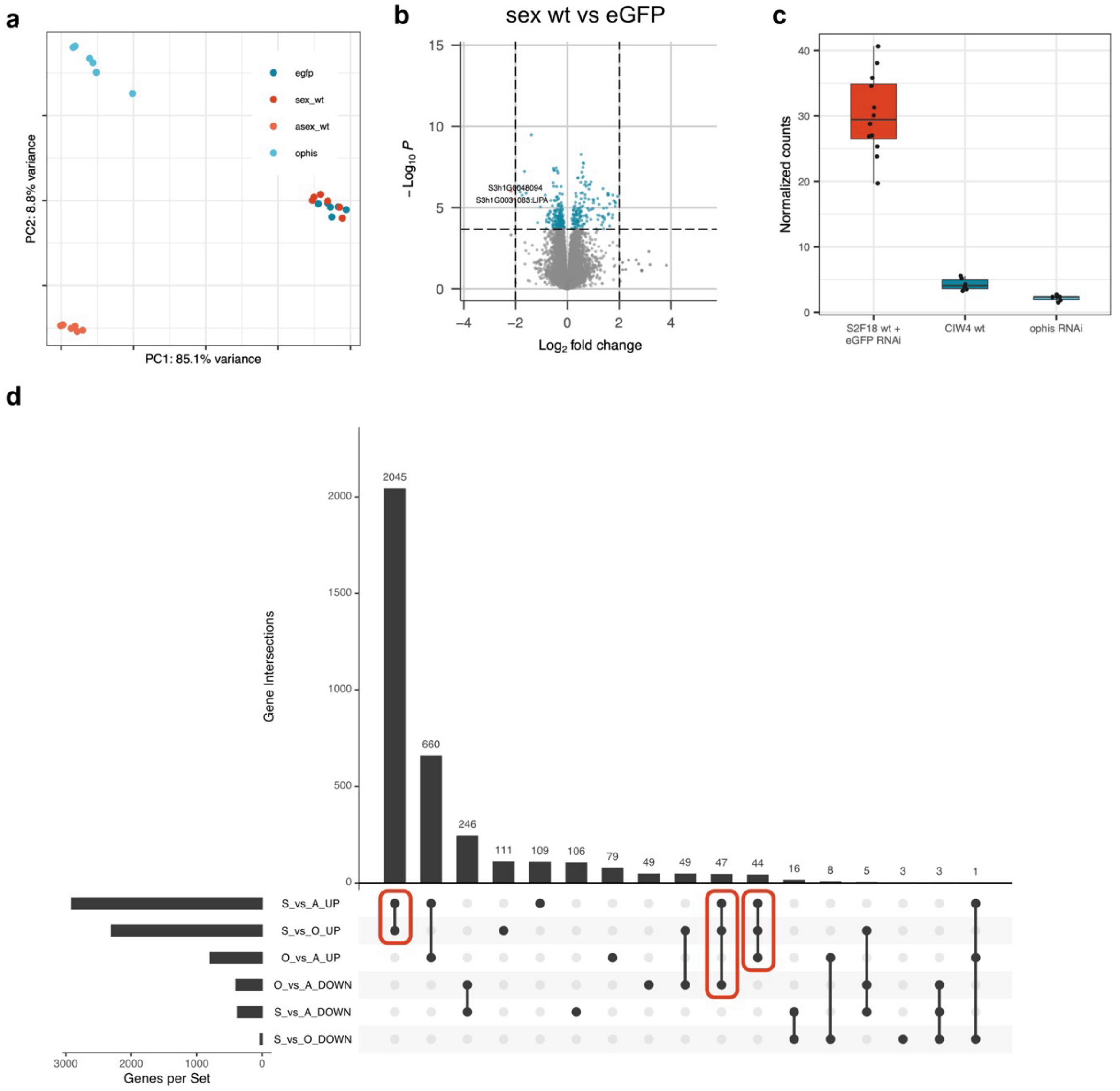

Figure 20c, Supporting Table S8) and then defined genes upregulated in the sexual control (wild-type & eGFP RNAi) vs. *ophis* RNAi and sexual control vs. asexuals wild-type (

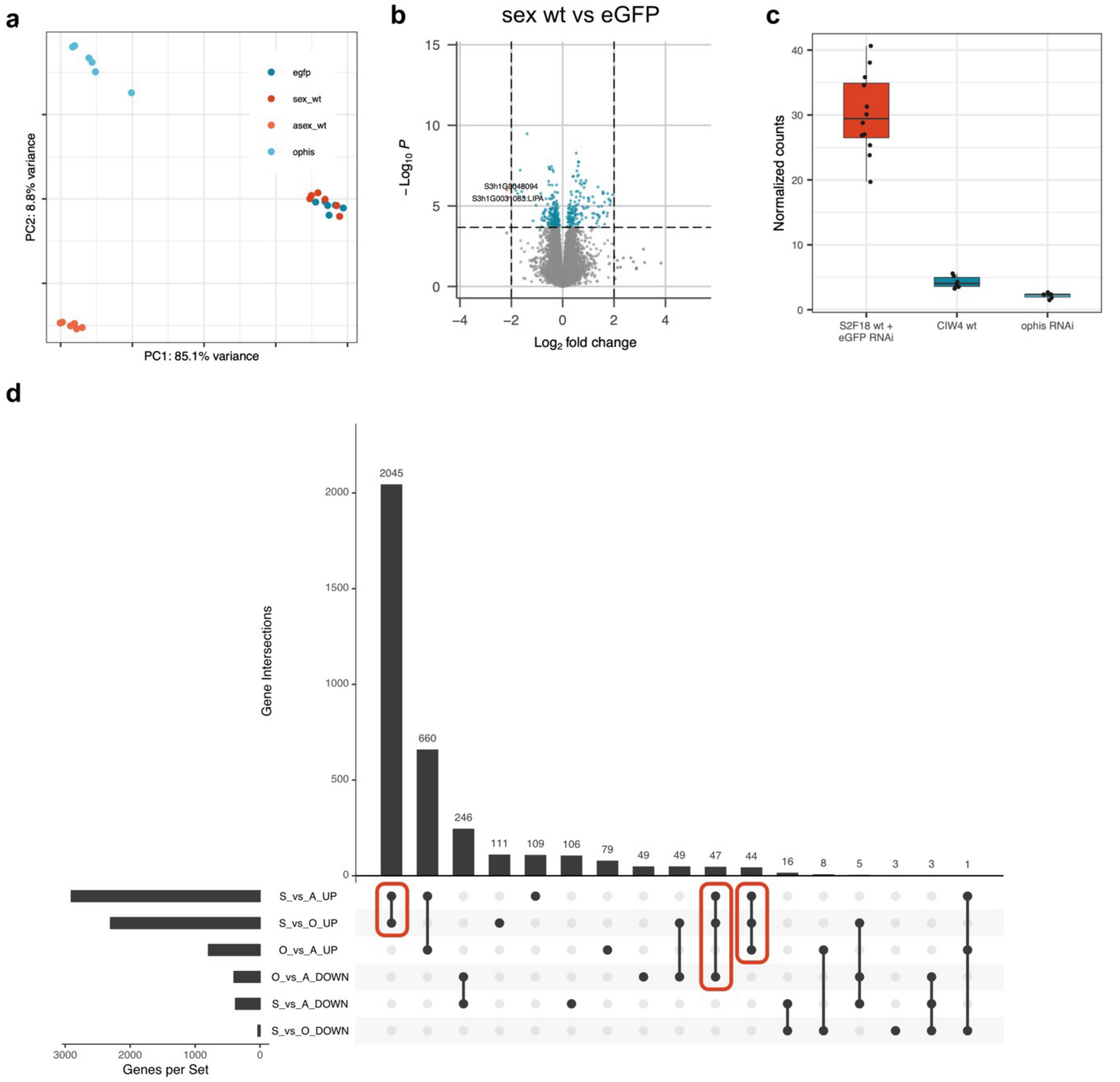

Figure 20d-g) as high-confidence reproduction-related genes.

Enrichment analysis of these 2,136 reproduction-related genes revealed 30 GO terms that were statistically significantly enriched. 9 of these terms were related to cilium assembly or other cilium related terms, likely due to an enrichment in sperm-related processes in animals with developed testes (Figure 21, Figure 22). Using these data we clearly see that identification of reproduction-related genes via simple comparison of sexual and asexual wild-type would lead to the identification of genes that are also differentially expressed between ophis RNAi and asexual wild type. I.e., genes that are likely different between strains – largely independent of their role in the reproductive system. Indeed, we find 1,829 genes that are upregulated in both these contrasts, but also 1,213 that are downregulated. Enrichment analysis of these strain-enriched genes, showed enrichment in GO terms related to immunity (Figure 23, Figure 24).

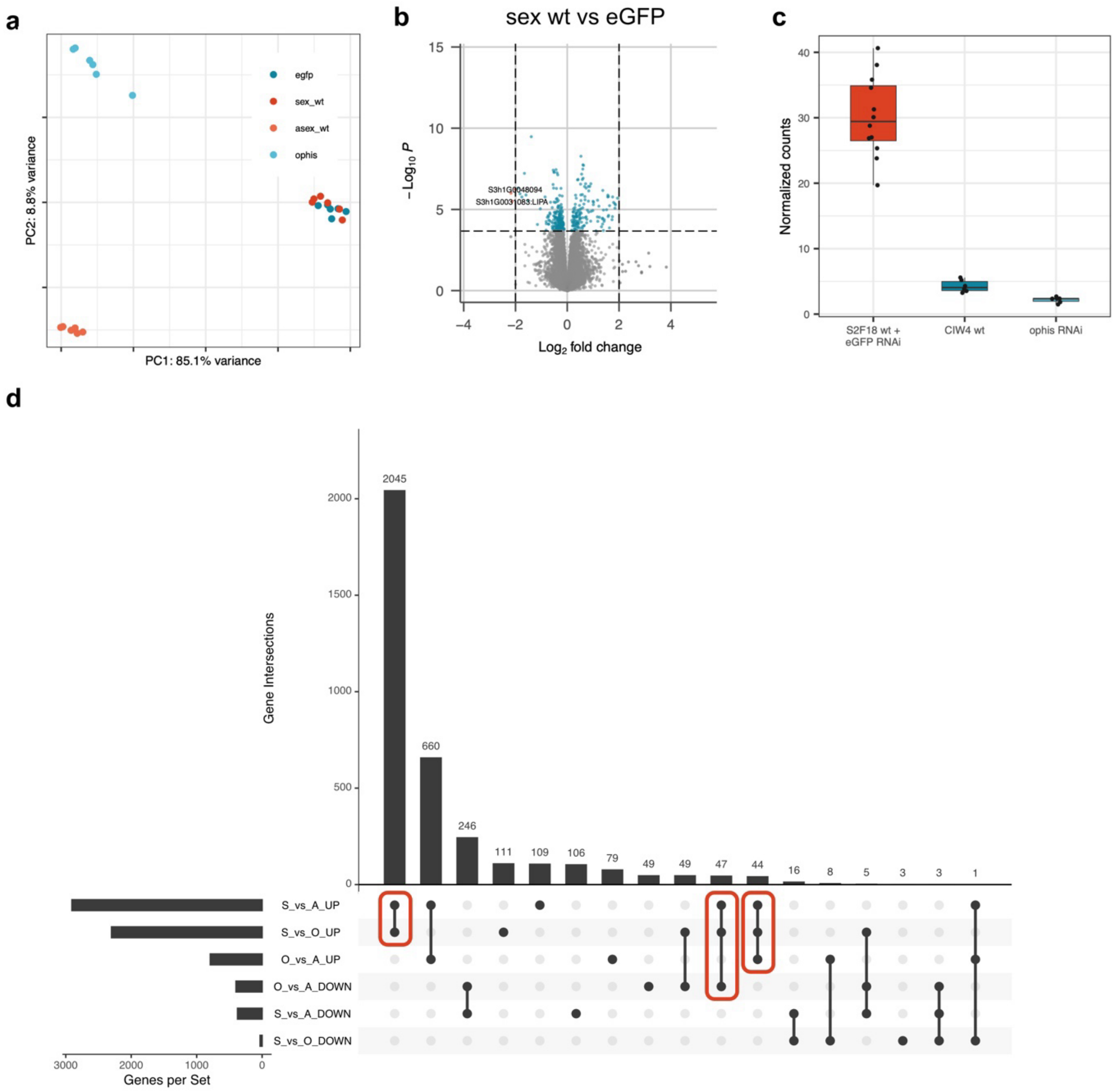

Figure 20 Identification of reproduction-related genes using RNA interference (RNAi) and differential expression analysis. **a** Principal component analysis of the expression of the four treatment groups: *eGFP* RNAi, *ophis* RNAi, sexual S2F18 wild-type, and asexual CIW4 wild-type, each with six replicates. Minimum expression filtered counts were normalized, and log2 transformed using the trimmed mean of the M-values (TMM) method and regression using the voom function of the R package limma. **b** Volcano plot from differential expression analysis between sexual wild-type and *eGFP* RNAi, showing minimal differences. **c** Normalized read counts for the RNAi target *ophis* (S3h1G0033988) in sexual control, asexual wild type, and *ophis* RNAi, indicating successful knock-down. **d** Upset plot summarizing the results from differential expression calls in Fig. 2d-f of the main text. Red boxes outline the sets designated as high-confidence reproduction-related transcripts.

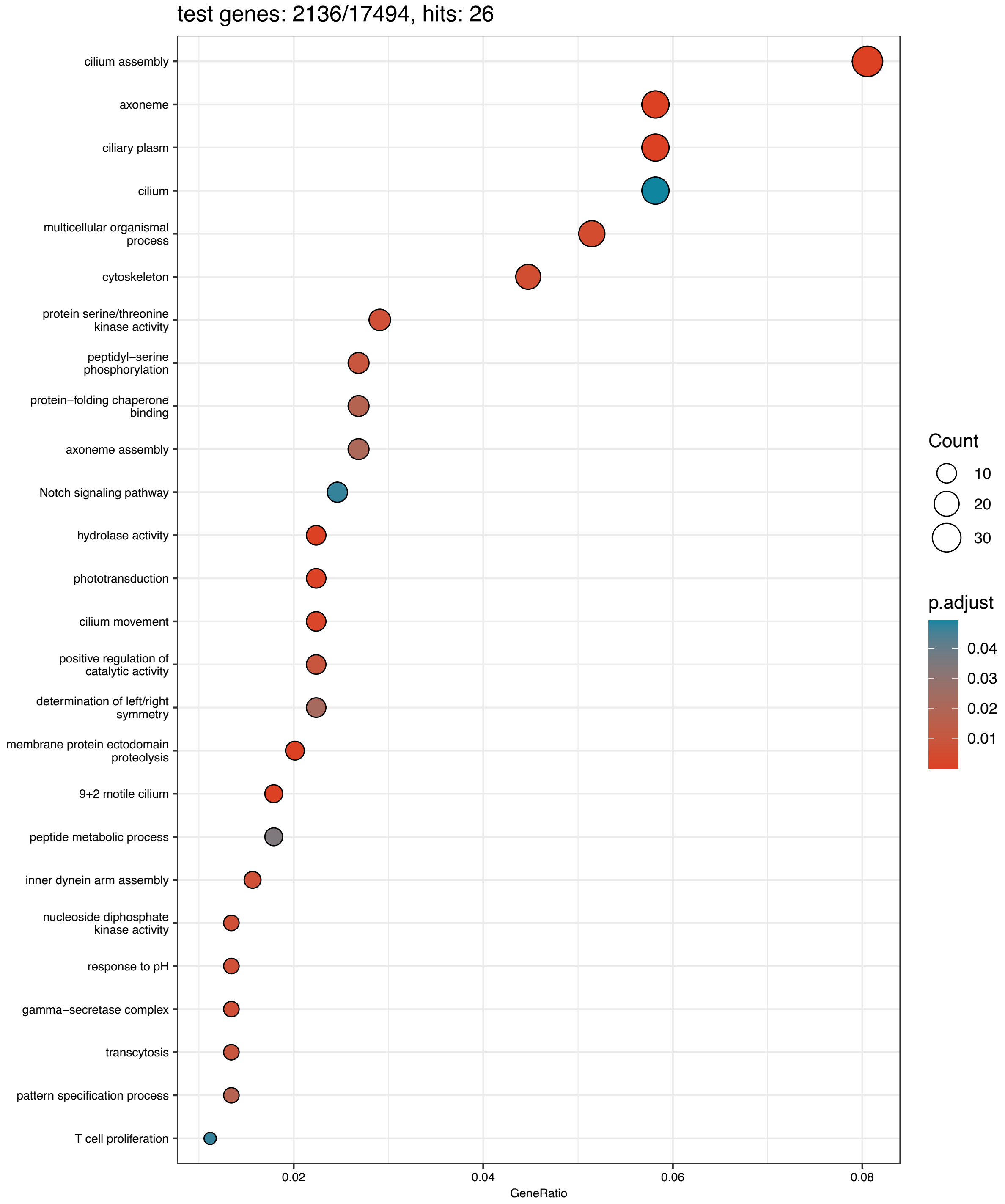

Figure 21 Enrichment dotplot showing GeneRatio and number of enriched genes for the 30 statistically significantly enriched GO terms in the set of high-confidence reproduction related genes.

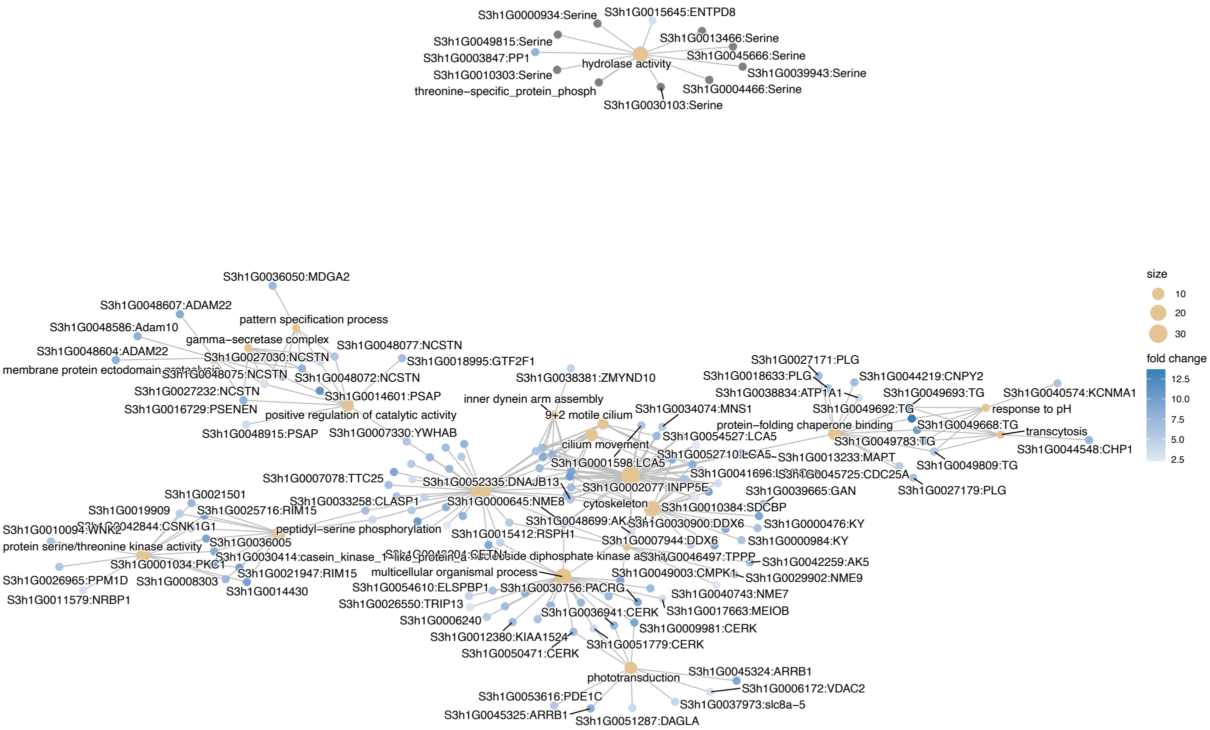

Figure 22 Gene-Concept Network plot showing the enriched GO terms and the associated genes for the set of high-confidence reproduction-related genes.

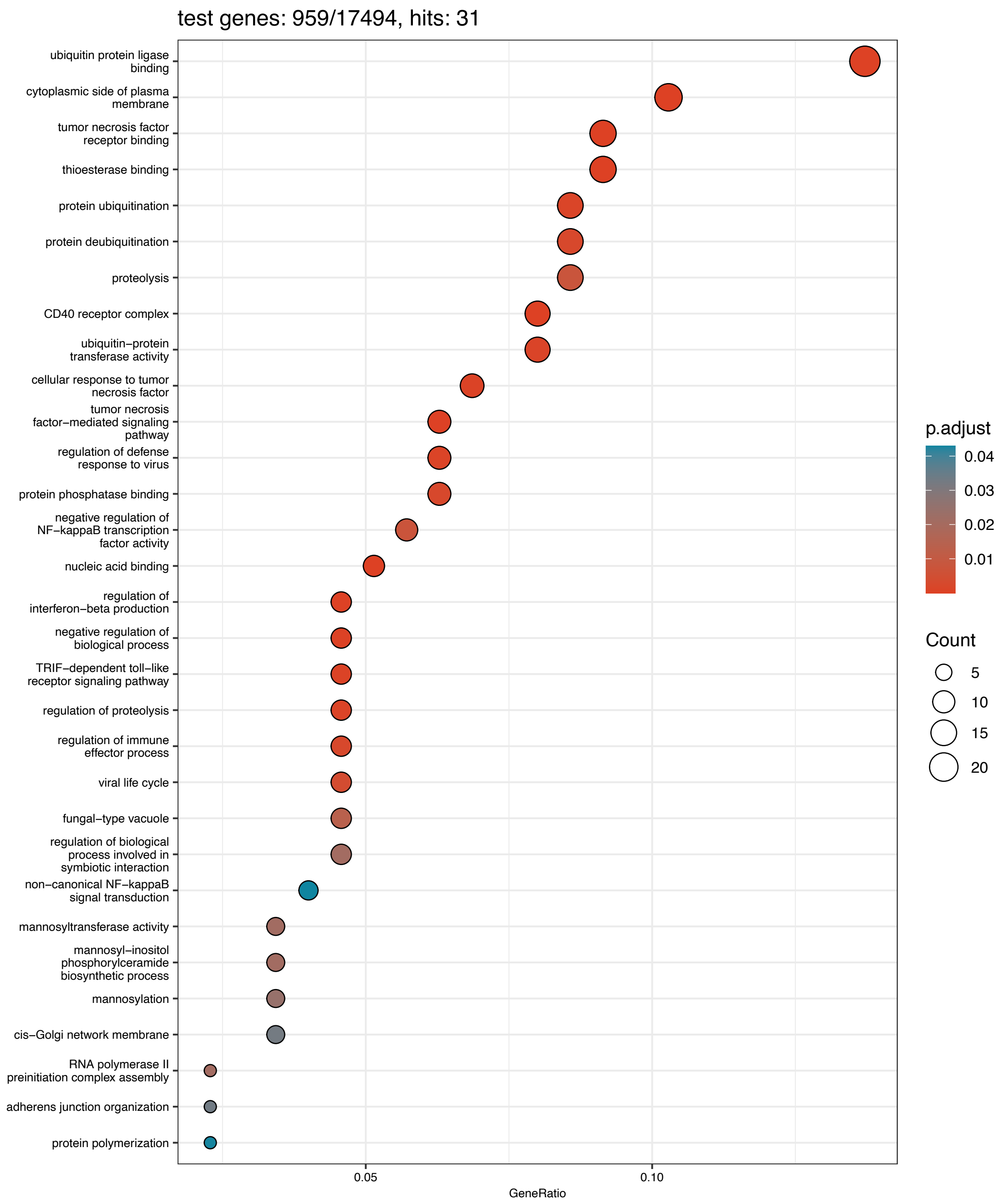

Figure 23 GO enrichment analysis of transcripts identified as enriched in one strain, but not due to the reproductive system.

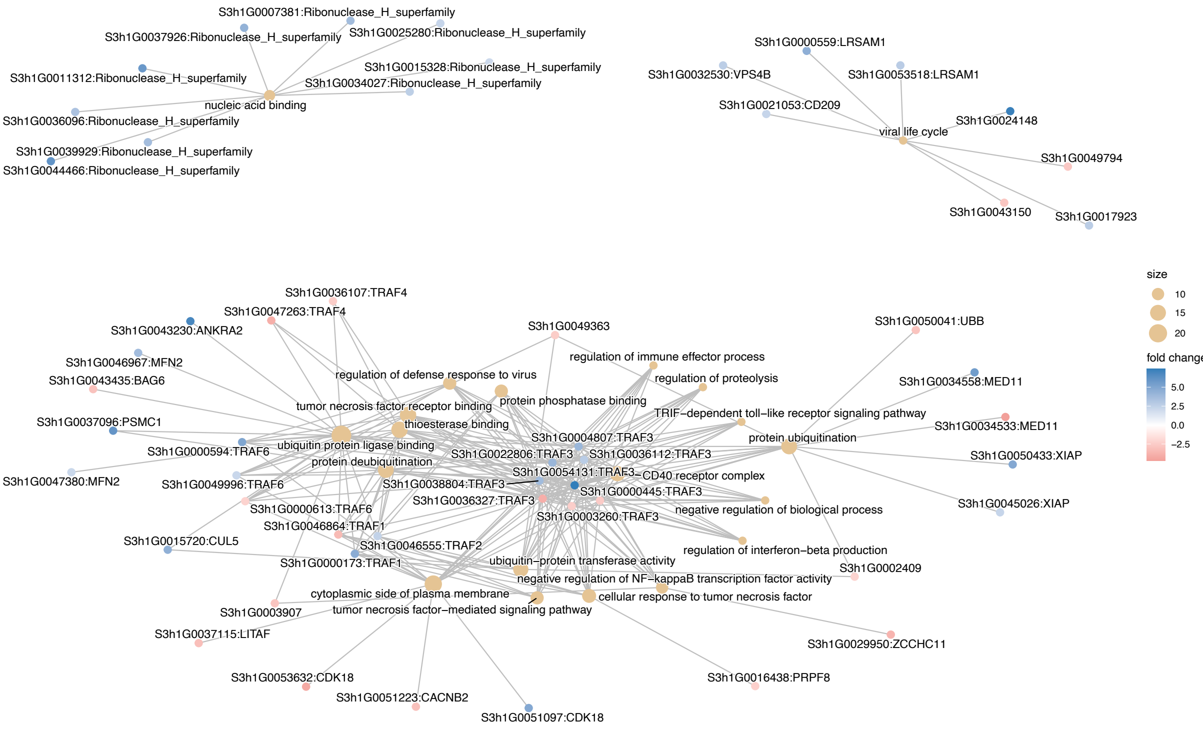

Figure 24 Gene-Concept Network plot showing the 31 enriched GO terms and the associated genes for strain-enriched genes.

### 8 Gene loss

To probe for potential gene losses in the vicinity of the breakpoints or for possible proximity effects on the expression of important reproductive system development genes, we analyzed the 112 gene models within a 1 Mb window flanking the Chr1/3 translocation breakpoints. To systematically analyze potential gene losses, we used the whole-genome alignment-based annotation pipeline TOGA[8]. Since TOGA is reference annotation centric, we again chose haplotype 1 of the sexual genome (schMedS3) as our reference and improved upon our previous annotation of the sexual genome via a liftover of haplotype 2 (Supporting Table S3). We then used our whole-genome alignments to identify genes that were flagged as lost or potentially lost in the asexual schMedA2 genome. In addition, we assessed the general conservation of genes using whole-genome alignments of the high-quality genomes of the three most closely related *Schmidtea* species—*S. polychroa*, *S. lugubris*, and *S. nova*[9].

We then intersected the TOGA results with our RNAi-based annotation of reproduction-related genes (see section reproduction-related genes). In addition, we annotated 76 genes known from the literature to be important for germline development to assess if they were in the vicinity of the genomic rearrangements (Supporting Table S8).

With these data sets in hand, we analyzed 112 gene models within a 1 Mb window flanking the breakpoint. Of those, 9 were flagged as lost or potentially lost in both haplotypes of the asexual strain. However, the flagged genes were scattered rather than forming a contiguous block, none of them were conserved in the three sister species of *S. mediterranea* (Figure 25; Supporting Table S9), and 7/9 overlapped repetitive regions. Together, these data suggest that the ”lost” genes are likely repetitive elements and that the Chr1/3 translocation did not lead to additional gene deletion in the vicinity (Supporting Table S9). In terms of reproduction-related genes, 13 mapped to within 1 Mb of the break point.

Because the Chr1/3 translocation was not associated with loss of reproduction-related genes, we next investigated whether the other large structural variations were accompanied by gene disruptions. Specifically, we examined the large inversions on Chr1 and Chr2, while ignoring the rearrangement on Chr4 due to possible issues with the scaffolding quality (Supporting Information: Hi-C Chromosome 4). In the 1 Mb regions flanking the breakpoint of inv(1)(10M_325M), we identified 2/23 genes as potentially lost, but neither one was included in the set of reproduction-related genes (Supporting Table S9). One of the lost genes was not conserved in *Schmidtea*, while the other was conserved but occurred within a tandem array of three gene copies, suggesting a recent duplication (Supporting Table S9; Figure 26). Examining the asexual-specific Chr2 inversion, we found 5/65 genes near the breakpoints flagged as lost or potentially lost. One of them was annotated as reproduction-related but none were conserved across the genus, and three were in regions highly enriched for repeats. In addition to the potential gene losses, we detected an exon loss in one gene within CIW4 haplotype 2 (Supporting Table S9; Figure 26). Two candidates are currently under investigation via knock-down assays.

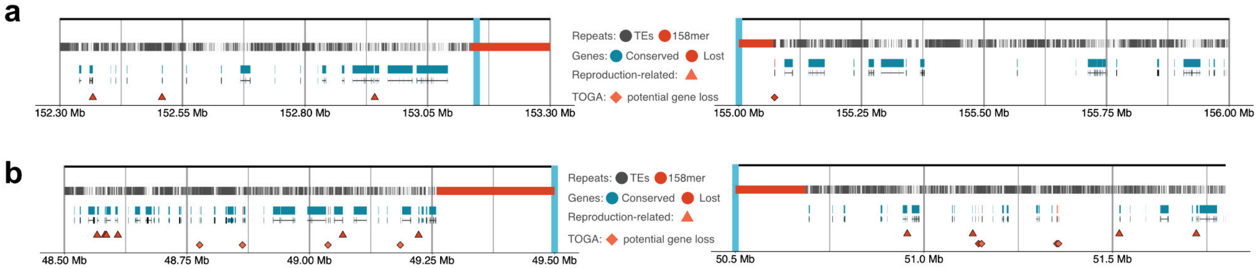

Figure 25 Gene loss analysis in the homologous regions of the Chr1/3 translocation breakpoints in the sexual genome assembly **a-b** Enlarge view of the regions outlined in Figure 3 a-b of the main text, showing the distribution of transposable elements (TEs), the 158mer repeats (158mer), and gene annotations. Triangles indicate high-confidence reproduction-related genes, and diamonds mark genes flagged as potentially lost in both haplotypes of the asexual schMedA2 assembly using the gene annotation pipeline TOGA.

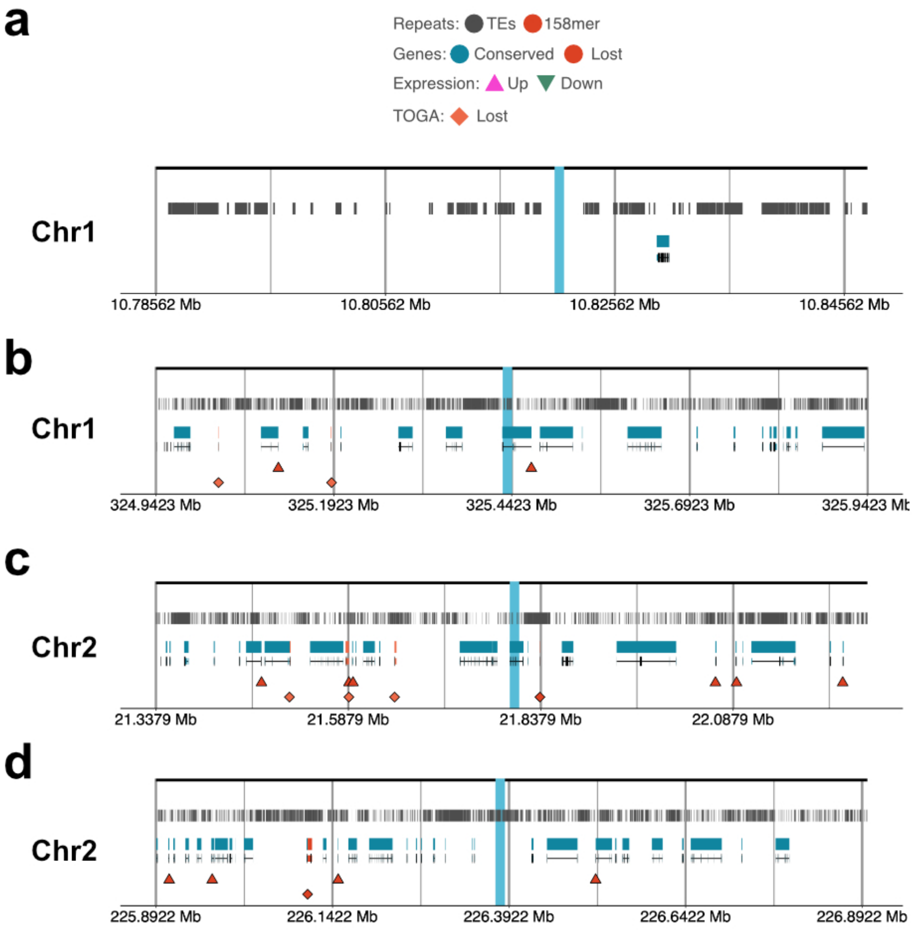

Figure 26 Genomic context on the sexual genome assembly (schMedS3) haplotype 1 of chromosome inversion breakpoints of the Chr1 Inversion 1 (a-b) and Chr2 inversion (c-d). Shown are the distribution of transposable elements (TEs) and gene annotations. Triangles indicate high-confidence reproduction-related genes, and diamonds mark genes flagged as potentially lost in both haplotypes of the asexual schMedA2 assembly using the gene annotation pipeline TOGA.

### 9 Enrichment analyses

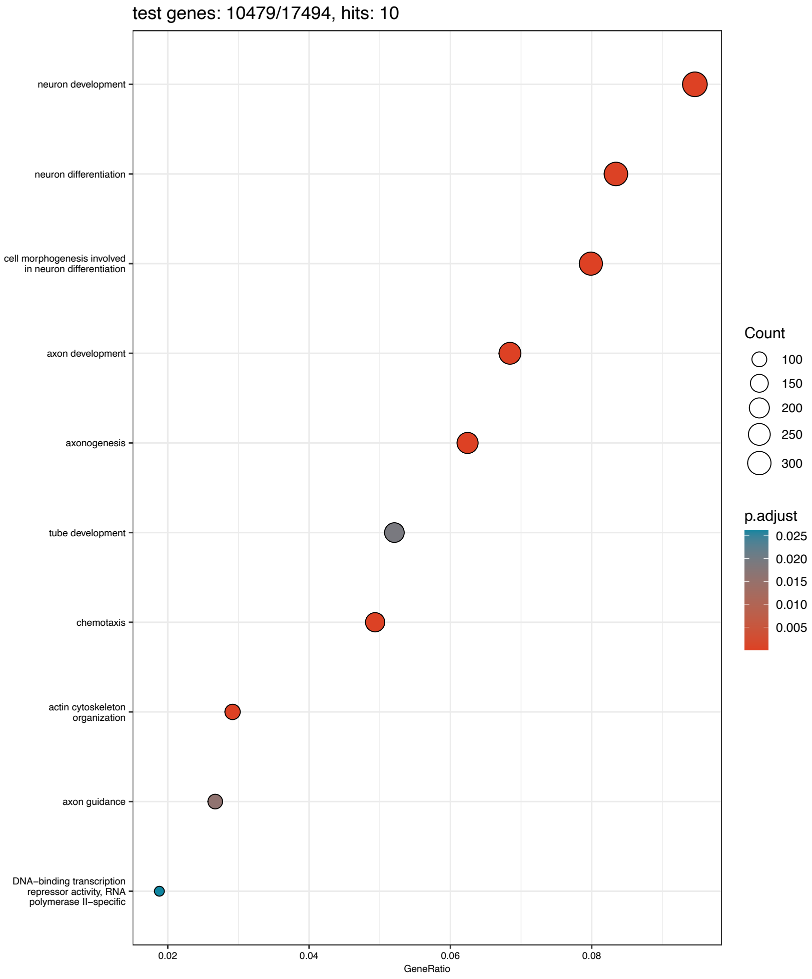

Figure 27 Enrichment dotplot showing GeneRatio and number of enriched genes for the 10 statistically significantly enriched GO terms in the set of single-copy genes used for the $d_N/d_S$ analysis.

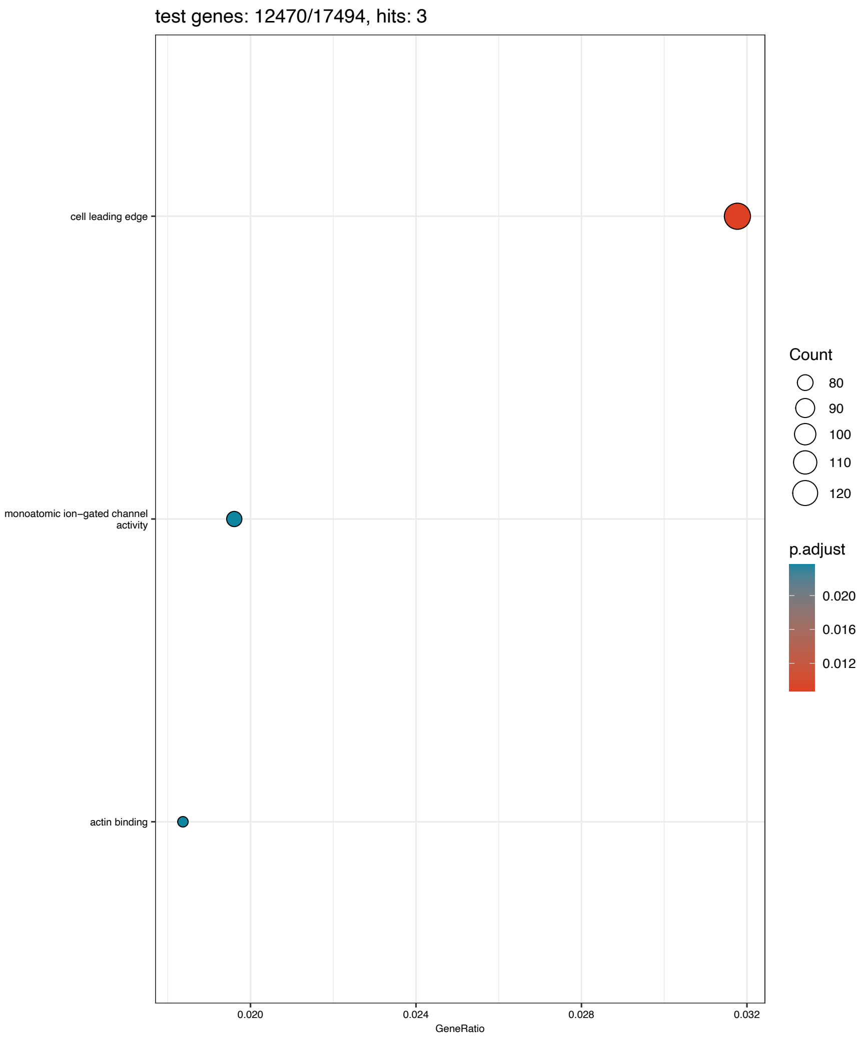

Figure 28 Enrichment dotplot showing GeneRatio and number of enriched genes for the 3 statistically significantly enriched GO terms in genes used for the piN/piS analysis.

### 10 Dataset intersection

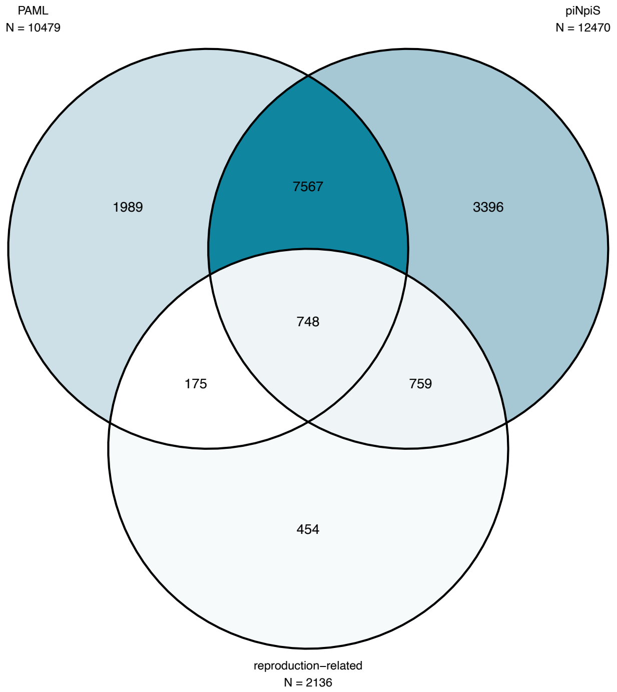

Figure 29 Venn diagram showing the overlap between genes used in the dN/dS analysis (PAML), the piN/piS analysis(piNpiS), and genes annotated as reproduction-related.

### 11 Population genomics

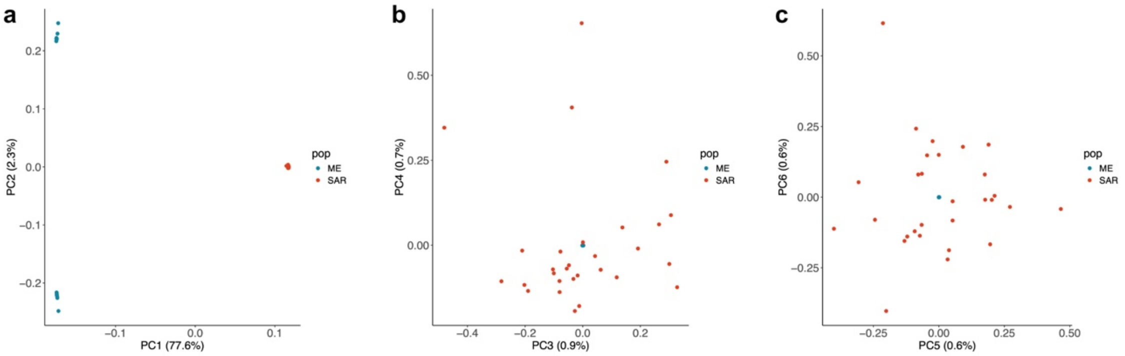

Figure 30 Principal component (PC) analysis of whole-genome sequencing data for Menorca (ME) and Sardinia (SAR) populations. Shown are (a) PC1 & PC2, (b) PC3 & PC4, and (c) PC5 & PC6.

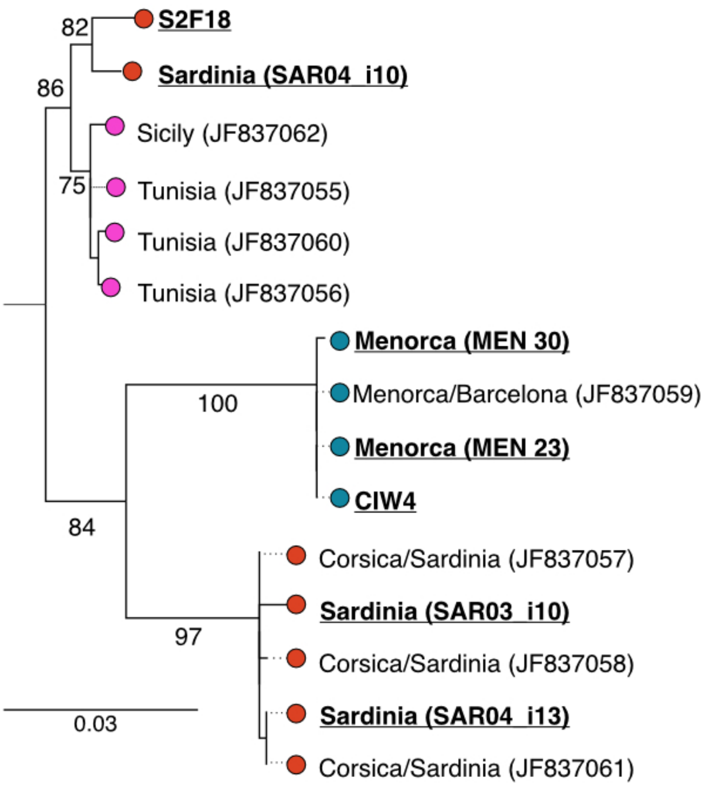

Figure 31 Maximum-likelihood phylogeny of Sardinia (SAR), Menorca (MEN), the asexual laboratory strain from Barcelona (CIW4), the sexual laboratory strain from Sardinia (S2F18), and publicly available sequences of a fragment of the COI barcoding gene. We find high similarity between MEN and CIW4 and more diversity within SAR, which does not appear monophyletic. However, note the low node support, which is based on 100 non-parametric bootstraps.

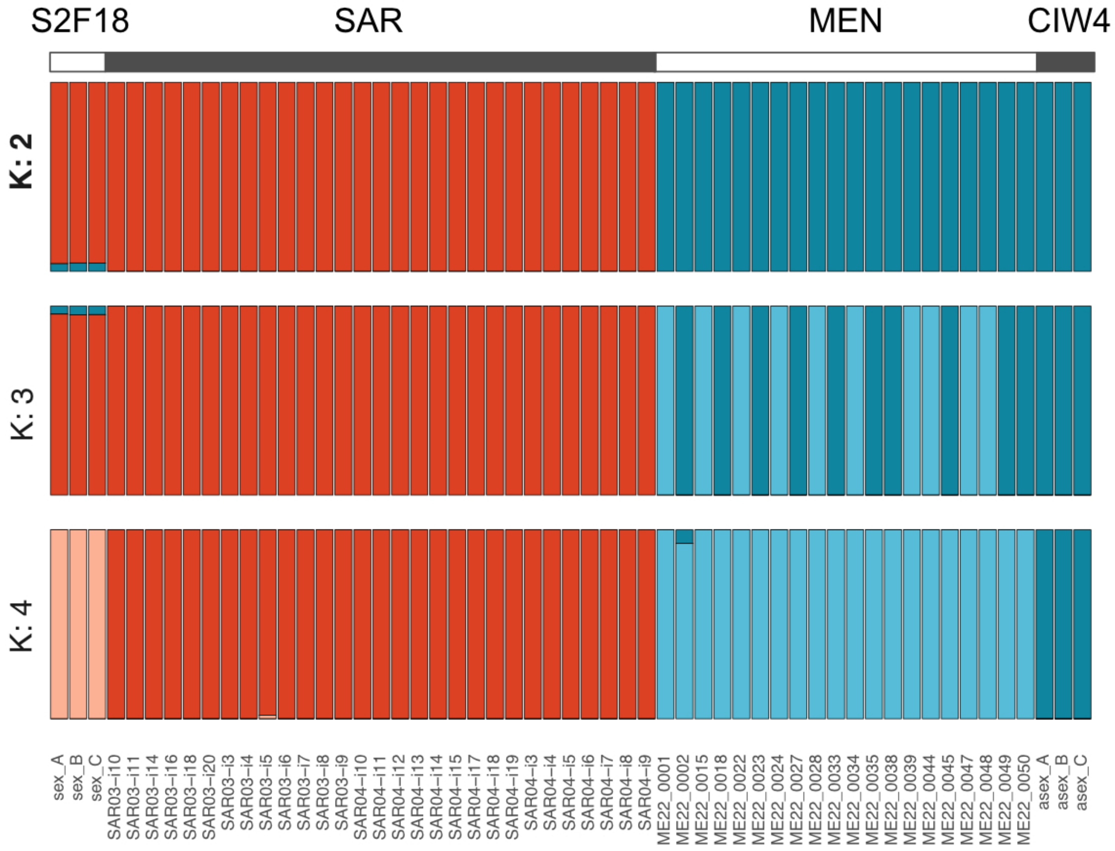

Figure 32 Population structure inferred using ngsAdmix. Best K = 2 was identified using Evanno's method.

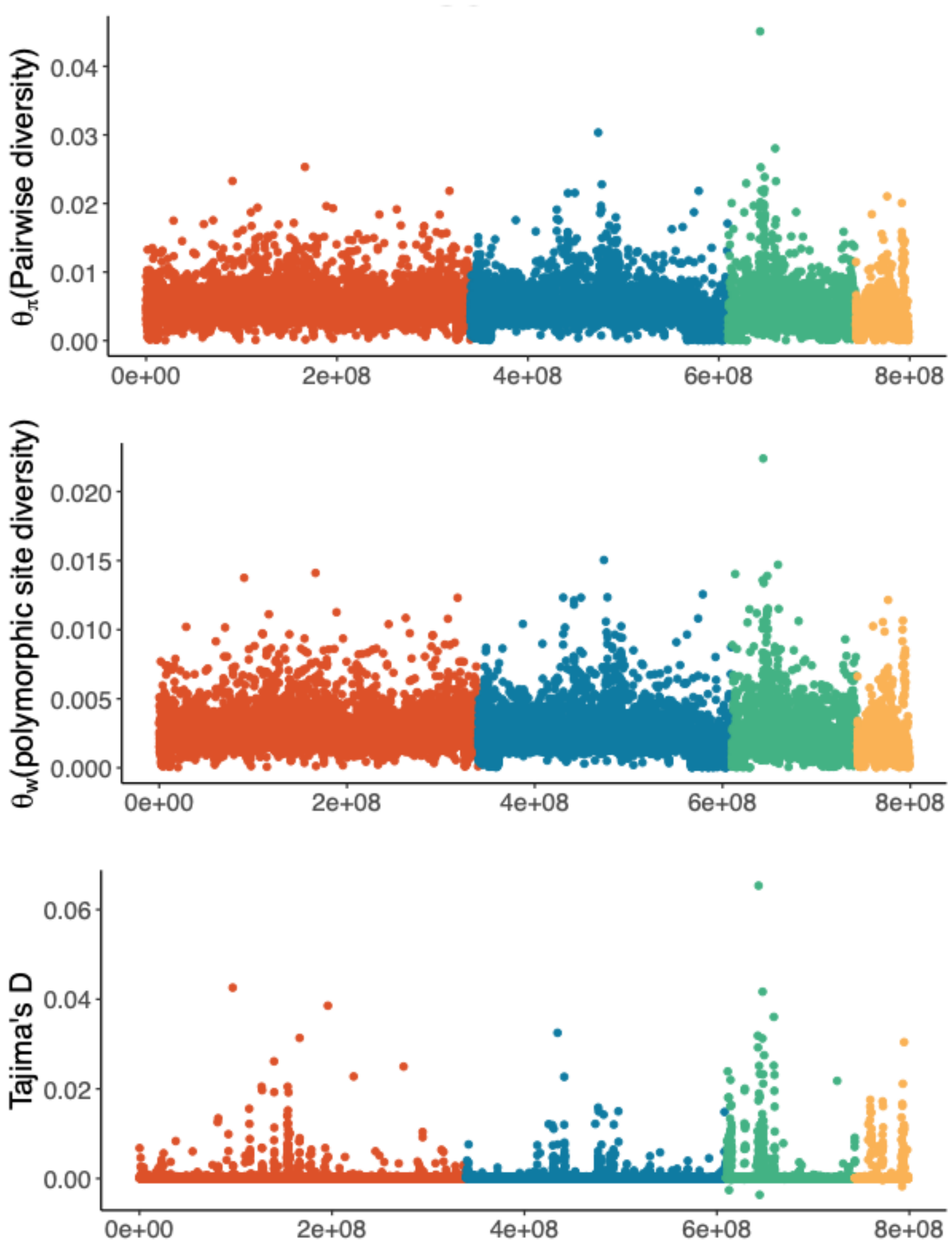

Figure 33 Population genomic parameters for the asexual population from Menorca (MEN). Show are theta estimates based on nucleotide diversity, polymorphic site distribution and Tajima's D inferred for 50Kb windows across the chromosome scaffolds.

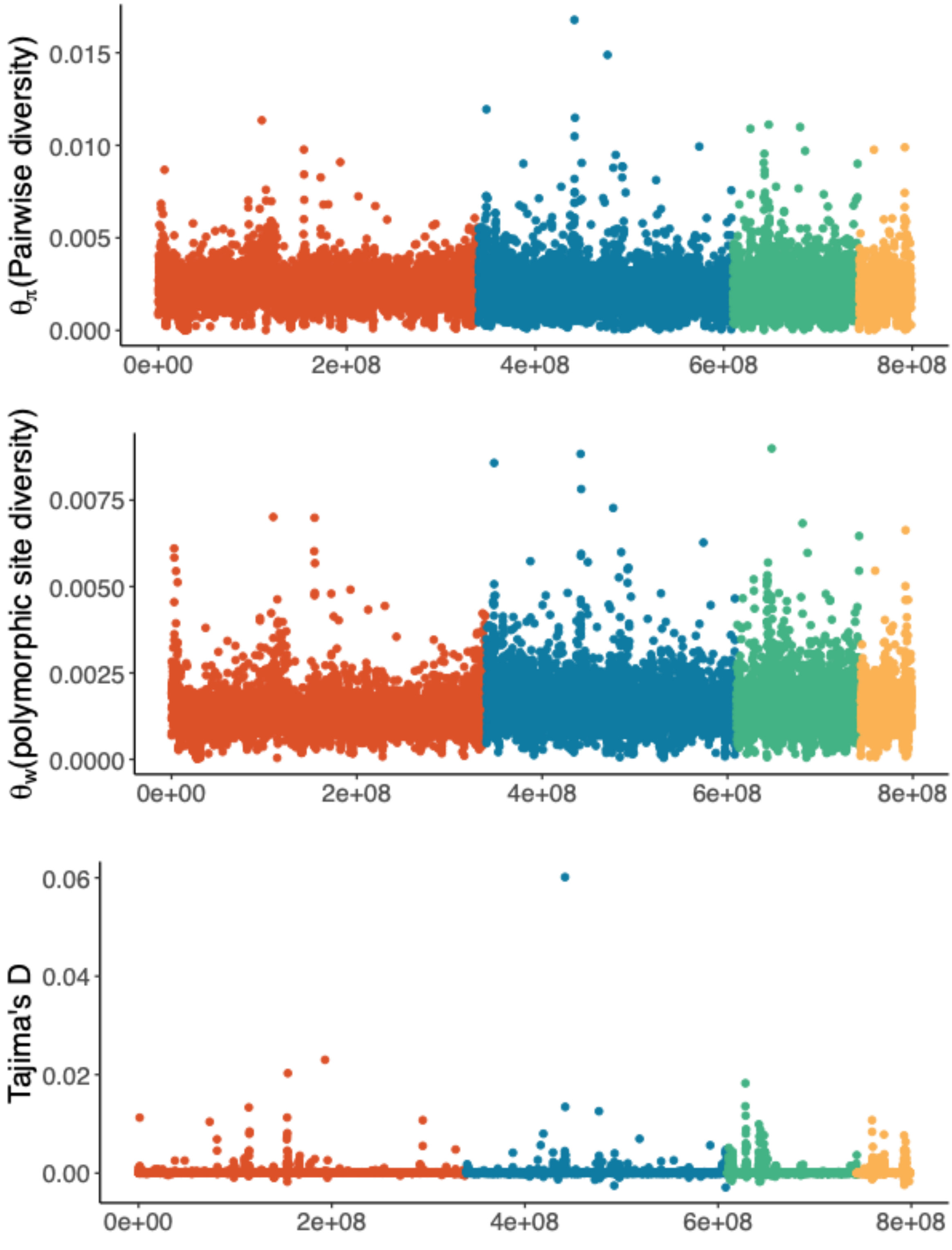

Figure 34 Population genomic parameters for the asexual population from Sardinia (SAR). Show are theta estimates based on nucleotide diversity, polymorphic site distribution and Tajima's D inferred for 50Kb windows across the chromosome scaffolds.

Table 7 Mean Diversity ($\pi$) and Watterson ($W$) estimators of the population parameter theta ($\theta$) in SAR and MEN, both genome-wide and for each chromosome scaffold. The last column indicates the grouping of chromosomes according to the test presented in the table below.

| $\theta$ estimator | population | median | mean | SD | chromosome | Wilcoxon group |
|---|---|---|---|---|---|---|
| $\pi$ | SAR | 0.00229 | 0.00233 | 0.00087 | chr1 | A |
| $\pi$ | SAR | 0.00209 | 0.00219 | 0.00110 | chr2 | B |
| $\pi$ | SAR | 0.00208 | 0.00222 | 0.00120 | chr3 | B |
| $\pi$ | SAR | 0.00213 | 0.00218 | 0.00105 | chr4 | B |
| $\pi$ | SAR | 0.00219 | 0.00225 | 0.00103 | genome-wide | |
| $\pi$ | MEN | 0.00463 | 0.00504 | 0.00219 | chr1 | A |
| $\pi$ | MEN | 0.00415 | 0.00449 | 0.00246 | chr2 | B |
| $\pi$ | MEN | 0.00383 | 0.00441 | 0.00316 | chr3 | C |
| $\pi$ | MEN | 0.00268 | 0.00319 | 0.00270 | chr4 | D |
| $\pi$ | MEN | 0.00429 | 0.00463 | 0.00254 | genome-wide | |
| $W$ | SAR | 0.00121 | 0.00129 | 0.00053 | chr1 | A |
| $W$ | SAR | 0.00148 | 0.00157 | 0.00066 | chr2 | B |
| $W$ | SAR | 0.00153 | 0.00163 | 0.00074 | chr3 | C |
| $W$ | SAR | 0.00144 | 0.00151 | 0.00065 | chr4 | D |
| $W$ | SAR | 0.00136 | 0.00146 | 0.00064 | genome-wide | |
| $W$ | MEN | 0.00239 | 0.00264 | 0.00125 | chr1 | A |
| $W$ | MEN | 0.00213 | 0.00238 | 0.00137 | chr2 | B |
| $W$ | MEN | 0.00204 | 0.00240 | 0.00174 | chr3 | C |
| $W$ | MEN | 0.00154 | 0.00185 | 0.00156 | chr4 | D |
| $W$ | MEN | 0.00221 | 0.00246 | 0.00142 | genome-wide | |

Table 8 Wilcoxon test showing SAR has significantly lower values of $\theta_\pi$ and $\theta_W$ than MEN. Tests are two-sided.

| $\theta$ estimator | group1 | group2 | n1 | n2 | statistic | p |
|---|---|---|---|---|---|---|
| $\pi$ | SAR | MEN | 15752 | 15463 | 36511700 | < 2.2e-16 |
| $W$ | SAR | MEN | 15752 | 15463 | 53283438 | < 2.2e-16 |

Table 9 All pairwise Wilcoxon tests for differences in $\theta_\pi$ and $\theta_W$ between chromosomes. All tests are two-sided and corrected for multiple testing using the Benjamini-Hochberg procedure.

| $\theta$ estimator | population | group1 | group2 | n1 | n2 | statistic | p | p.adj | signif |
|---|---|---|---|---|---|---|---|---|---|
| $\pi$ | SAR | chr1 | chr2 | 6717 | 5307 | 20158263 | 4.7E-35 | 2.82E-34 | **** |
| $\pi$ | SAR | chr1 | chr3 | 6717 | 2649 | 9956697 | 2.37E-19 | 7.11E-19 | **** |
| $\pi$ | SAR | chr1 | chr4 | 6717 | 1079 | 4038495 | 1.52E-09 | 3.04E-09 | **** |
| $\pi$ | SAR | chr2 | chr3 | 5307 | 2649 | 7018898 | 0.916 | 0.916 | ns |
| $\pi$ | SAR | chr2 | chr4 | 5307 | 1079 | 2830526 | 0.555 | 0.833 | ns |
| $\pi$ | SAR | chr3 | chr4 | 2649 | 1079 | 1417973 | 0.708 | 0.85 | ns |
| *W* | SAR | chr1 | chr2 | 6717 | 5307 | 12440432 | 2E-178 | 1.2E-177 | **** |
| *W* | SAR | chr1 | chr3 | 6717 | 2649 | 6013554 | 3.6E-132 | 1.1E-131 | **** |
| *W* | SAR | chr1 | chr4 | 6717 | 1079 | 2740903 | 6.98E-38 | 1.4E-37 | **** |
| *W* | SAR | chr2 | chr3 | 5307 | 2649 | 6735214 | 0.002 | 0.003 | ** |
| *W* | SAR | chr2 | chr4 | 5307 | 1079 | 3006429 | 0.009 | 0.009 | ** |
| *W* | SAR | chr3 | chr4 | 2649 | 1079 | 1555912 | 0.000021 | 3.15E-05 | **** |
| $\pi$ | MEN | chr1 | chr2 | 6654 | 5255 | 20512606 | 1.88E-59 | 2.82E-59 | **** |
| $\pi$ | MEN | chr1 | chr3 | 6654 | 2548 | 10576677 | 1.06E-75 | 2.12E-75 | **** |
| $\pi$ | MEN | chr1 | chr4 | 6654 | 1006 | 5179419 | 6.8E-173 | 4.1E-172 | **** |
| $\pi$ | MEN | chr2 | chr3 | 5255 | 2548 | 7258113 | 1.58E-09 | 1.58E-09 | **** |
| $\pi$ | MEN | chr2 | chr4 | 5255 | 1006 | 3685260 | 1.38E-87 | 4.14E-87 | **** |
| $\pi$ | MEN | chr3 | chr4 | 2548 | 1006 | 1673004 | 8.91E-46 | 1.07E-45 | **** |
| *W* | MEN | chr1 | chr2 | 6654 | 5255 | 20207816 | 1.96E-48 | 2.94E-48 | **** |
| *W* | MEN | chr1 | chr3 | 6654 | 2548 | 10177919 | 2.64E-50 | 5.28E-50 | **** |
| *W* | MEN | chr1 | chr4 | 6654 | 1006 | 4842587 | 7.6E-116 | 4.6E-115 | **** |
| *W* | MEN | chr2 | chr3 | 5255 | 2548 | 7053488 | 0.000122 | 0.000122 | *** |
| *W* | MEN | chr2 | chr4 | 5255 | 1006 | 3454871 | 7.28E-54 | 2.18E-53 | **** |
| *W* | MEN | chr3 | chr4 | 2548 | 1006 | 1599602 | 8.46E-31 | 1.02E-30 | **** |

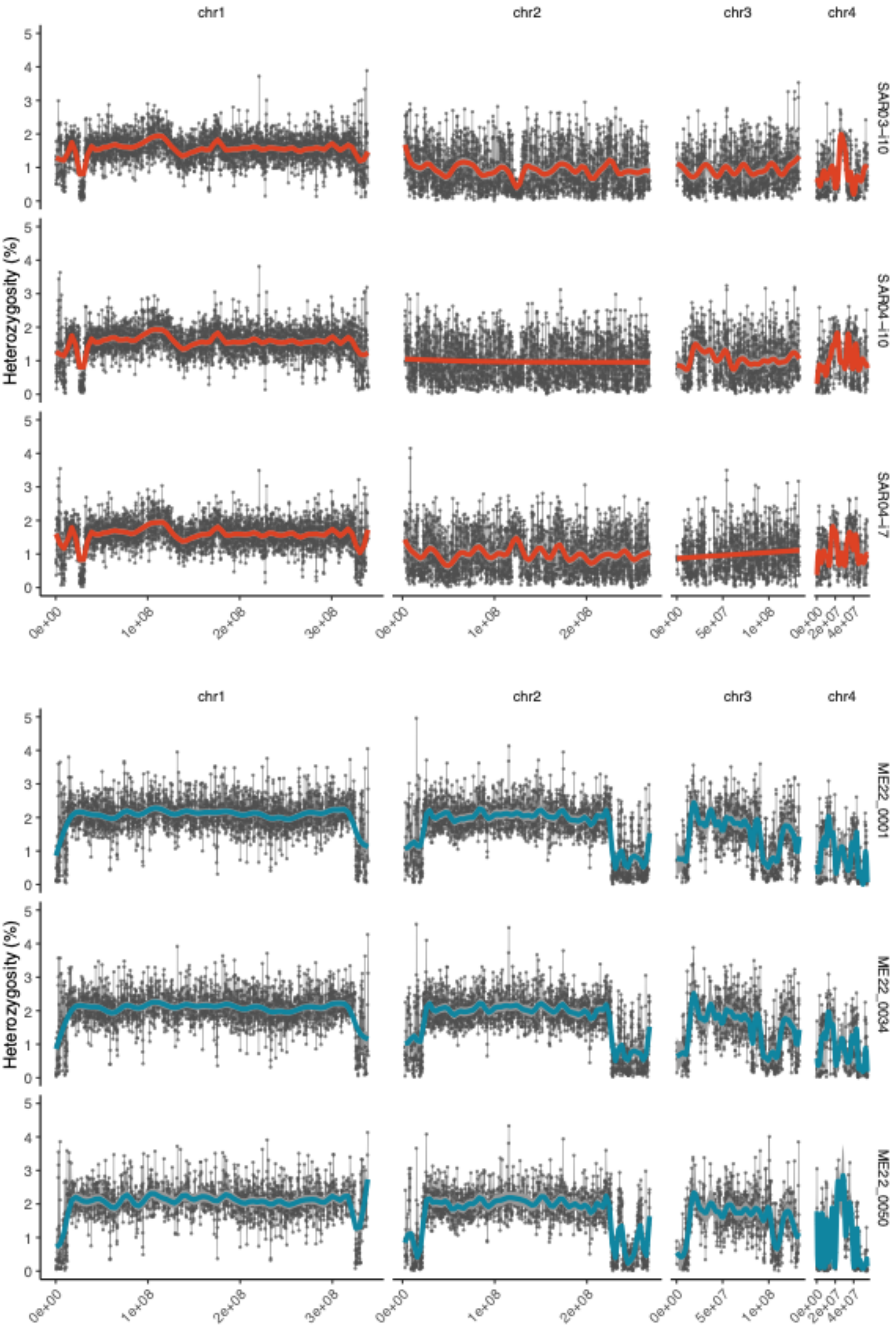

Figure 35 Observed heterozygosity calculated as the percentage of heterozygous sites per called site (min 10 reads, FS < 30). Points show value within a 100kb window and are connected by thin lines. The colored lines show the fit of a general additive model (GAM) the r package ggplot2. The fit was calculated using the formula ‘y ~ s(x, k = 50)’. Envelope around the colored lines represent 95% confidence intervals. Red: three samples from the sexual SAR population; blue: three samples from the asexual MEN population.

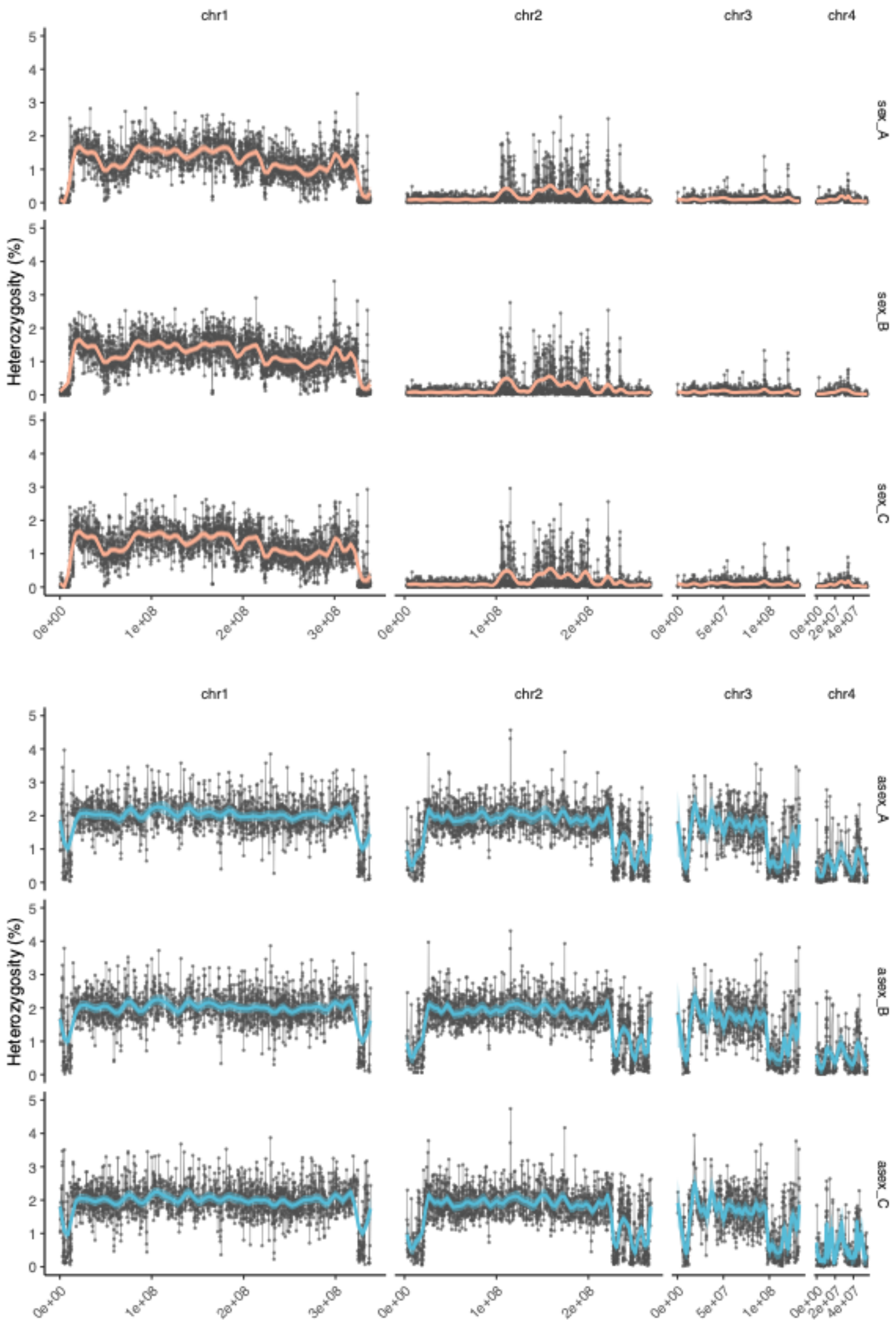

Figure 36 Observed heterozygosity calculated as the percentage of heterozygous sites per called site (min 10 reads, FS < 30). Points show value within a 100kb window and are connected by thin lines. The colored lines show the fit of a general additive model (GAM) fit via the r package ggplot2. The fit was calculated using the formula ‘y ~ s(x, k = 50)’. Envelope around the colored lines represent 95% confidence intervals. Red: three samples from the sexual lab strain (S2F18); blue: three samples from the asexual lab strain (CIW4).

### 12 LTR insertion age

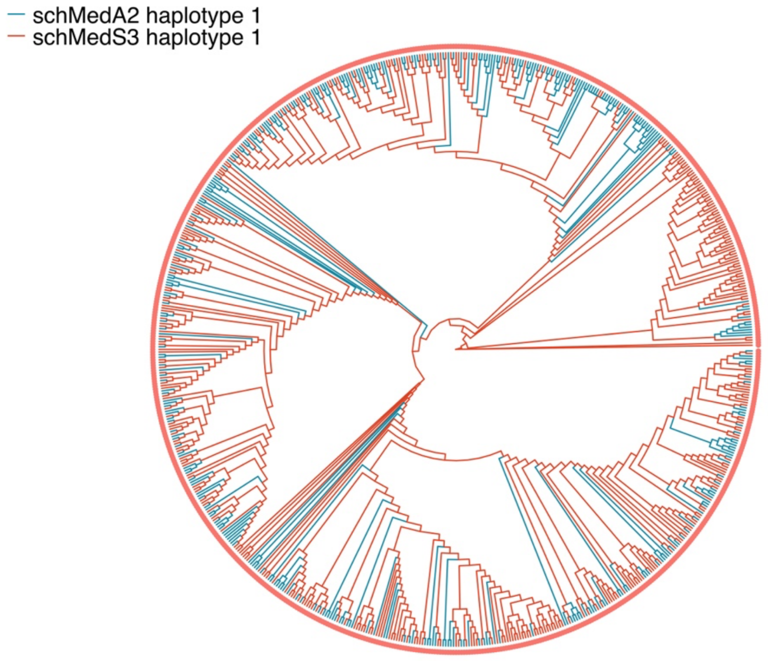

Figure 37 Phylogenetic tree of LTR transposons identified to have inserted exclusively in the asexual (schMedA2) and sexual (schMedS3) genomes.

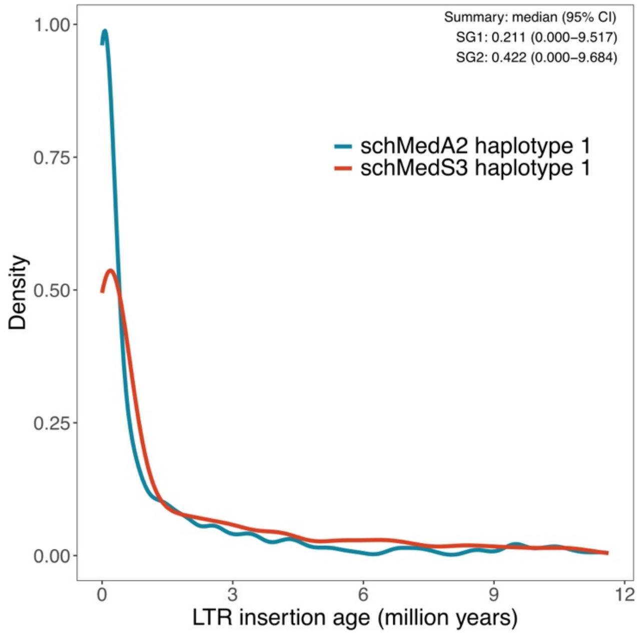

Figure 38 Density plot showing the distribution of divergence between LTR copies. Median insertion age is 0.211 Ma and 0.422 Ma in the asexual and sexual genome, respectively.
